# Support for the social buffering hypothesis, especially under unpredictable precipitation regimes

**DOI:** 10.1101/2025.04.30.651380

**Authors:** Roberto Salguero-Gómez, Gabriel S Santos, Lauren Brent, Samuel J L Gascoigne, Michael B Bonsall, Josh A Firth, Gregory Albery, Rahul Mondal, Maja Kajin, Katrina J Davis

**Affiliations:** Department of Biology, University of Oxford, South Parks Road, Oxford, OX1 3RB, United Kingdom; Vale Institute of Technology - Sustainable Development, Rua Boaventura da Silva 955, Belém, Pará 66055-090, Brazil; Department of Biology, Espirito Santo Federal University, Avenida Fernando Ferrari, 514 Vitória, Espírito Santo 29075-910, Brazil; Centre for Research in Animal Behaviour, University of Exeter, Exeter, EX4 4QG, United Kingdom; School of Biological Sciences, University of Aberdeen,Tillydrone Ave, Aberdeen, AB24 2TZ, United Kingdom; School of Biology, Faculty of Biological Sciences, University of Leeds, Leeds, UK; School of Natural Sciences, Trinity College Dublin, Dublin, Republic of Ireland; Department of Biostatistics and Epidemiology, International Institute for Population Sciences, India

**Keywords:** comparative demography, cooperative breeding, demographic buffering, environmental stochasticity, phylogenetic analysis, sociality continuum

## Abstract

Sociality is hypothesised to buffer populations against environmental stochasticity by stabilising key vital rates, yet this has rarely been tested across diverse taxa. We compiled demographic time series from 87 populations for 66 animal species spanning 12 taxonomic classes to test whether sociality promotes demographic buffering and whether this relationship depends on environmental context. Using stochastic elasticities and phylogenetically comparative analyses, we find broad support for the social buffering hypothesis: more social species exhibit lower sensitivity of population growth to vital rate variance (lower stochastic elasticity to vital rate variances, |*T*_σ_|), particularly in juvenile survival and reproduction. Contrary to expectations, juvenile vital rates, but not adult survival, showed the strongest buffering patterns in social taxa, suggesting life-stage-specific social mechanisms. Furthermore, we show that social species invest more in maximising the average performance of adult survival and reproduction (higher stochastic elasticity to vital rate means, *T*_μ_), consistent with adaptive strategies favouring canalisation of fitness-related traits. This buffering benefit is especially relevant in highly unpredictable environments: in climates with erratic precipitation, social species have greater demographic advantages, exhibiting even lower *T*_σ_ values. Together, these findings provide support for the social buffering hypothesis, especially in highly stochastic environments. Social traits confer some demographic stability under moderate climatic conditions, but may become more relevant under extreme stochasticity—an insight particularly important for understanding population resilience in a rapidly changing world.

## Introduction

Sociality is crucial for the persistence of natural populations. The way in which individuals interact within social groups influences not only individual fitness (Silk, 2007b; Snyder-Mackler et al., 2020), but also shapes population-level processes such as survival (Creel & Christianson, 2008; Salguero-Gómez, 2024), reproduction (Lukas & Clutton-Brock, 2012; Salguero-Gómez, 2024), dispersal (Greenwood, 1980), and resistance to environmental pressures (Oro, 2020; Rubenstein & Lovette, 2007; Salguero-Gómez, 2024). Social organisations, from ephemeral aggregations to highly coordinated closed social groups, have evolved across diverse taxa as adaptive responses to predation (Krause & Ruxton, 2002), resource limitation (Koenig & Dickinson, 2016), or reproductive constraints (Emlen, 1982). Interactions that occur between members of a social organisation often mediate access to resources and mates (Clutton-Brock, 2009), affect stress physiology (Young & Bennett, 2013), and enable individuals to cope with environmental uncertainty (Bourke, 2011). As such, sociality is a central feature of the viability of natural populations.

Social organisation is a multifaceted trait. However, this characteristic has been historically often boiled down to a binary trait that is mostly examined in a few taxonomic groups. Indeed, while much of the previous literature distinguishes species as either social or non-social (Lukas & Clutton-Brock, 2012), this dichotomy is now often considered as insufficient for capturing the complexity and gradation of social behaviours observed across the animal kingdom. For instance, some spiders exhibit facultative sociality, forming colonies under high resource availability but remaining solitary otherwise (Avilés et al., 2012). Moreover, most comparative research in this area has focused on birds (Jetz et al., 2012) and mammals (Lukas & Clutton-Brock, 2012), where social systems have been extensively studied. This narrow focus has left a gap regarding the generality of how social organisations influence demography across animals, including in less well-studied taxa such as reptiles, amphibians, invertebrates (but see (Crump, 2015; Doody et al., 2013; Wong & Balshine, 2011)), and marine organisms. A growing consensus now acknowledges the need for integrative frameworks that account for the dimensionality of sociality (Albery et al., 2024; Doody et al., 2013; Firth et al., 2024; Salguero-Gómez, 2024), including the kinds, intensity, and frequency of said interactions.

There are clear implications regarding how sociality should allow populations to persist via the social buffering hypothesis (Klug & Bonsall, 2014; Rubenstein & Lovette, 2007; Shine, 1978). This hypothesis posits that social interactions, particularly those involving cooperative care, resource sharing, or communal defence, can mitigate the negative demographic impacts of environmental fluctuations. By pooling resources, enhancing survival, or smoothing reproductive success over time, sociality may reduce demographic variance and stabilise population growth. However, the strength and consistency of such buffering effects are expected to vary depending on the type of social organisation, with cooperative breeders hypothesised to benefit more strongly from such mechanisms than solitary, loosely gregarious, or communal species (Salguero-Gómez, 2024).

Population ecologists have extensively examined the drivers of demographic buffering (J.-M. Gaillard & Yoccoz, 2003; Morris et al., 2008; Pfister, 1998; Tuljapurkar, 1982), a life-history strategy whereby natural selection reduces temporal variability in vital rates that most affect population fitness. This strategy is predicted to evolve under environmental stochasticity, as a means of stabilising population growth in the face of unpredictable ecological pressures (Bruijning et al., 2020; Hilde et al., 2020; Le Coeur et al., 2022; Tuljapurkar, 1982). Traditionally, demographic buffering has been assessed, among other ways (Lewontin & Cohen, 1969), by examining the coefficient of variation in vital rates or their temporal variance (Hilde et al., 2020; Pfister, 1998). However, recent theoretical and empirical work has emphasised the utility of stochastic elasticity analysis in explicitly quantifying how changes in the mean and variance of vital rates affect long-term population growth rate (λ) (Ezard & Coulson, 2010; S. Gascoigne et al., 2024; S. J. L. Gascoigne et al., 2023; Giaimo & Traulsen, 2023; Haridas & Tuljapurkar, 2005).

Stochastic elasticities directly quantify the buffering capacity of a population due to its vital rates. In this framework, the stochastic elasticity to the mean of vital rates (*T*_μ_) reflects the relative importance of selection for improving average demographic performance, whereas the stochastic elasticity to vital rate variance (*T*_σ_) captures the sensitivity of λ to interannual variability in those same vital rates. Importantly, *T*_μ_ and *T*_σ_ probe alternative buffering mechanisms by which populations can persist in variable environments (S. J. L. Gascoigne et al., 2024). Across species, the impacts of mean vital rates on population growth rate in variable environments covary with life history strategies. For example, mean adult survival is especially important in species with longer generation times (Oli & Dobson, 2003; Saether & Bakke, 2000). Furthermore, *T*_σ_ also covaries with life history strategies. Specifically, the quantification of *T*_σ_ across taxa has demonstrated the increased sensitivity of species with shorter lifespans to variation in demographic rates relative to those with longer lifespans (Morris et al., 2008). Together, these metrics provide a powerful and evolutionarily relevant view of how organisms manage trade-offs between maximising fitness and buffering environmental uncertainty. Despite their theoretical relevance, *T*_μ_ and *T*_σ_ have rarely been applied in macroecological or behavioural studies—particularly not to test the role of sociality in modulating selection on demographic stability. Few studies have explicitly linked behavioural traits such as social organisation to either of these components of population dynamics, despite predictions that social behaviour may act as a buffering mechanism (Rubenstein & Lovette, 2007; Salguero-Gómez, 2024). Bridging this gap is essential to understand whether social systems merely enhance survival and reproduction, or also reduce vulnerability to environmental stochasticity—a distinction with major implications for forecasting population resilience under global change.

Here, we collate data on 87 populations and 66 animal species spanning 12 taxonomic classes, thus encompassing a wide diversity of life histories and social systems, to examine the demographic consequences of sociality through the lens of demographic buffering. We test four key hypotheses. (H1) Social species should exhibit stronger demographic buffering, operationalised as lower temporal variance vital rates, relative to solitary taxa, in line with the social buffering hypothesis (Rubenstein & Lovette, 2007). This buffering is predicted to be especially pronounced in cooperative breeders, where group-living and alloparental care redistribute energetic costs and reduce reproductive uncertainty. (H2) This buffering effect should be primarily mediated via adult stages, particularly adult survival, which tends to be more canalised in long-lived species (J.-M. Gaillard & Yoccoz, 2003; Morris et al., 2008; Pfister, 1998), while juvenile rates should contribute minimally to buffering patterns. (H3) More social species should exhibit higher stochastic elasticity to the mean of adult survival and reproduction than to juvenile survival and maturation, indicating stronger selection for maximising the average performance of key adult vital rates in social species compared to solitary ones. Finally, (H4) the hypothesised relationship between sociality and demographic buffering should be modulated by environmental stochasticity, such that buffering effects in social species diminish under highly variable conditions, *e.g.,* in arid regions with high rainfall variability. Indeed, recent evidence has emerged suggesting that demographic buffering can be ineffective in extremely stochastic environments (Rodriguez-Caro et al., 2021; Santos et al., 2024). To test these four hypotheses, we develop a comparative phylogenetic framework that incorporates time series of demographic information, species-specific climatic variability, and a recently introduced sociality continuum, which classifies species from more solitary to fully social (Salguero-Gómez, 2024).

## Methods

To test the hypotheses that sociality modulates demographic buffering in animals via adult survival and its interaction with environmental stochasticity, we performed a suite of phylogenetically-informed comparative analyses. Our analyses build on and extend two recent methodological frameworks: one developing novel demographic metrics to quantify the degree of demographic buffering in natural populations using matrix population models (MPMs, hereafter) (Santos et al., 2023), and another one introducing a continuum of social organisation for animal species, from more solitary to more social (Salguero-Gómez, 2024). By incorporating longitudinal demographic data specifically designed to capture temporal variation in survival and reproduction, and linking them to the climatic patterns associated with each studied location, we also test the hypothesis that the expected relationships between sociality and demography break down in extreme environments.

### Demographic data

To quantify the strength of demographic buffering across species, we first compiled demographic time series data for animal populations from the COMADRE Animal Matrix Database (v.4.32.3.1) (Salguero-Gomez et al., 2016), a global open-access repository of age- and stage-structured matrix population models (MPMs) for animals. We complemented this database with demographic data on *Suricata suricatta* (meerkat) from (Conquet et al., 2023), which also includes an annual long-term series of MPMs.

To ensure the reliability, comparability, and biological realism of these MPMs, we applied a stringent series of selection criteria. Indeed, in its version 4.23.3.1, COMADRE contains 3,488 MPMs from 429 populations across 415 peer-reviewed studies. These studies, though peer-reviewed, were not necessarily built with comparative demography in mind by each single set of authors (Salguero-Gomez et al., 2021). As such, we imposed the following set of selection criteria using R (v. 4.3.2) and the R packages Rcompadre and Rage (Jones et al., 2022), and popbio (Stubben & Milligan, 2007). We retained only MPMs that satisfied the following conditions:

- Wild populations only: We excluded MPMs parameterised with data from laboratory settings, captive populations (*e.g.*, zoos), or experimentally manipulated populations by filtering the metadata of the COMADRE R object by the variables ‘Captivity’ and ‘MatrixTreatment’. This step ensured that emergent demographic traits would reflect natural ecological and social dynamics.
- Proper matrix decomposition: We required that each MPM be decomposed into its survival-development (***U***) and reproduction (***F***) submatrices such that the overall MPM ***A*** = ***U*** + ***F*** (Caswell 2001). This structure allowed us to calculate standard life history traits and to perform matrix algebra reliably across species.
- Biological plausibility of vital rates: Using the ‘cdb_flag’ function in the R package Rcompadre, we excluded matrices with missing values or with biologically implausible values (*i.e.*., survival probabilities 0 < σ ≤ 1). MPMs containing ***F*** submatrices solely composed of zero-values were also excluded.
- Animals only: We retained only MPM associated with the kingdom *Animalia* by filtering the ‘Kingdom’ metadata, and thus excluding any entries from *Bacteria* or *Fungi*.
- Time series completeness: We grouped MPMs by species, study (based on the ‘Authors’, ‘Source’ and ‘YearPublication’ metadata), and population (‘MatrixPopulation’), and retained only those populations for which at least three annual MPMs were available. This step allowed us to estimate the impact of interannual variability in vital rates on population growth while minimising bias due to small sample sizes. We note that the primary results we show are largely insensitive to the available data duration of each study (Table S1).
- Population selection for multi-population species: For species represented by multiple studies (3 species in this study), we retained the studies containing the longest and best spatially replicated data.
- Humans: In COMADRE, we have access to temporally replicated MPMs for humans (*Homo sapiens sapiens*) for 42 countries (Nicol-Harper et al., 2018). However, to strike a compromise between having humans represented in this analysis, but not overwhelm the analyses with highly unbalanced population replications for any one species, and also due to the difficulties in assigning buffering abilities against environmental stochasticity to humans (Mondal et al., 2024), we retained only one human population: Spain.
- Standardising projection intervals: Because MPMs in COMADRE span different projection intervals (*e.g*., 6 months, 5 years; Figure S1), we back-transformed all MPMs to a 1-year time step using the methods by elevating each matrix element a_ij_ to the power of the frequency of study, following Salguero-Gómez and Gamelon (Salguero-Gomez & Gamelon, 2021) and Salguero-Gómez (Salguero-Gómez, 2024).

This set of selection criteria resulted in a subset of COMADRE for the next analytical steps, comprising 87 populations from 66 animal species, and totalling 955 MPMs (Figure S2).

### Estimates of demographic buffering

To quantify demographic buffering across the resulting subset of animal species, we focused on interannual variability in stage-specific survival (σ), maturation (γ), and reproduction (φ). However, as the dimension of the MPMs varied considerably (Figure S3), we collapsed all MPMs with dimension >2 (*i.e.*, representing more than two stages in the life cycle of the species) down to 2×2 MPMs, where the first stage represents juveniles (*J*) and the second stage adults (*A*). To do so, we implemented the collapsing criterion developed by Salguero-Gómez and Plotkin (2010), which allows for the collapsing of any MPM while retaining its eigenstructure. Following suggestions from Salguero-Gómez and Plotkin (2010), we kept the first stage unaltered, and collapsed into the adult stage from the second life cycle stage onwards. As such, the overall structure of the resulting MPMs contain four vital rates, shown in Eq. 1: juvenile survival (σ*_J_*), juvenile maturation (γ), adult survival (σ*_A_*), and reproduction (φ).

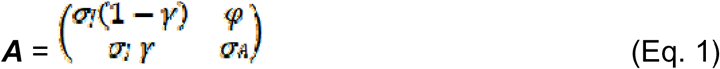

Next, we quantified the stochastic elasticities of population growth rate (λ*_s_*) to both changes in the mean and in the variance of vital rates across each population’s time series of MPMs. Briefly, stochastic elasticities describe the proportional change in long-term stochastic growth rate in response to small, proportional changes in a given vital rate, thus making them ideal tools to examine how natural populations respond to environmental stochasticity (Haridas & Tuljapurkar, 2005; Tuljapurkar et al., 2003). Following Haridas and Tuljapurkar (2005), we distinguished between two forms of elasticity: (i) the elasticity of the stochastic population growth rate (λ*_s_*) to mean of vital rates, *E_vr_*^μ^, which isolates the impact of changing the mean value of a given vital rate *vr* in the MPM (Eq. 1) while holding its variability constant; and (ii) the elasticity of λ*_s_* to the changes in the variance of a given vital rate, *E_vr_*^σ^, which isolates the impact of changing variability while holding the mean of said vital rate (and all others) constant. These two types of stochastic elasticities are critical for evaluating demographic buffering, which is defined as selection to reduce the sensitivity of λ_s_ to variability in vital rates (Hilde et al., 2020; Pfister, 1998; Santos et al., 2023).

We calculated these elasticities using population-specific MPMs constructed from each of the four annual vital rate estimates shown in Eq. 1: juvenile survival (σ*_J_*), juvenile maturation (γ), adult survival (σ*_A_*), and reproduction (φ). For each population, we implemented the approach detailed in Haridas and Tuljapurkar (Haridas & Tuljapurkar, 2005), which expresses λ*_s_* as a function of the full time series of MPMs and uses perturbation analysis to compute the stochastic elasticities. Specifically, we calculated *E_vr_*^μ^ as the derivative of log(λ*_s_*) with respect to the mean of the vital rate *vr*, and *E_vr_*^σ^ as the derivative with respect to its standard deviation, both obtained via numerical approximations using finite perturbations that are relative to the value of the vital rates under examination (S. J. L. Gascoigne et al., 2023; Haridas & Tuljapurkar, 2005). By separating effects of means and variances, this approach allows us to estimate the total elasticity of λ*_s_* to changes in the mean of vital rates (*T*_μ_) and the total elasticity of λ*_s_*to changes in the variance of vital rates (*T*_σ_) by summing across all vital rate elasticities in each population:

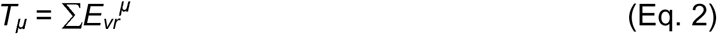

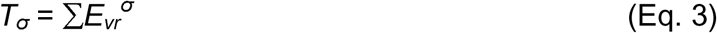

To compute these elasticities empirically, we used the ‘stoch.elas’ function in the popbio R package (Stubben & Milligan, 2007) as a baseline, which we then modified to generate separate elasticities for the mean and standard deviation of each vital rate using Monte Carlo simulation across MPM resamples. Specifically, for each population, we calculated the stochastic growth rate λ*_s_* from 10,000 iterations of matrix multiplication through the random time series of MPMs (one per year). Then, we introduced a small perturbation (1%) to either the mean or the variance of each vital rate independently while keeping all others fixed. We then recalculated λ*_s_* and took the finite-difference approximation of each elasticity (Haridas & Tuljapurkar, 2005). The separate components of the sum of mean elasticities (*T*_μ_) thus measures the aggregate selective importance of vital rate averages and helps us test (H3) whether more social species show higher stochastic elasticity to adult survival and reproduction than to juvenile rates, suggesting stronger selection on adult vital rate averages in social than solitary species. In contrast, the sum of variance elasticities (*T*_σ_) measures the aggregate selective importance of vital rate variances, and as such is directly relevant to our test (H1) of whether more social species exhibit reduced sensitivity to vital rate variance. As the distributions of *T*_μ_ and *T*_σ_ were right-skewed, we log_10_-transformed them before using them as response variables in our phylogenetic comparative analyses (below). We also applied a log_10_-transformation to the four vital rate stochastic elasticities to the mean (*E* ^μ^*, E* ^μ^, *E* ^μ^ and *E*_φ_ ^μ^) and to the variance (*E* ^σ^*, E* ^σ^, *E* ^σ^ and *E* ^σ^).

It is worth noting that, ecologically and evolutionarily, *T*_μ_ and *T*_σ_ (and their respective underlying vital rate components: *E_vr_*^μ^ and *E_vr_*^σ^) capture complementary forces. The stochastic elasticity to changes in mean vital rates reflects selection for life history traits that increase average performance (*e.g*., survival, reproduction), while the stochastic elasticity to changes in the variance of vital rates reflects selection against variability (*i.e*., demographic buffering). Under the demographic buffering hypothesis, we expect a higher degree of sociality to correlate with lower *T*_σ_, indicating reduced exposure of λ*_s_*to environmental noise. The framework from Haridas and Tuljapurkar (Haridas & Tuljapurkar, 2005) also contains a unique property associated with elasticity analyses, shown in Eq. 4. Specifically, with equal proportional changes in mean and variance in vital rates, the stochastic elasticity *E_vr_* is the summation of *E_vr_*^μ^ and *E_vr_*^σ^. Since, with all levels of variability in vital rates the sum of stochastic elasticity *E_vr_* sums to one, the summations of elasticity values *T*_μ_ and *T*_σ_ also equal to one. This property holds for all possible life histories. In turn, just like their deterministic counterparts (Takada et al., 2018), *T*_μ_ and *T*_σ_ are valuable tools for comparative analysis as their values are unbiased by the matrix dimensionality (*i.e.*, the number of stages in the MPM prior to downscaling to a 2×2 MPM, as done here), life history complexity (*e.g*., iteroparous *vs.* semelparous), or asymptotic properties (*e.g.*, declining, stable or increasing populations as per λ*_s_*) of the MPMs.

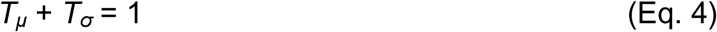

Importantly, *T*_μ_ and *T*_σ_ inhabit different numeric domains. *T*_μ_ is invariably positive as the summed impacts of minor increases in the mean of all vital rates yield an increase to the stochastic population growth rate λ*_s_* (Haridas & Tuljapurkar, 2005). On the other hand, *T*_σ_ is invariably negative as the summed impacts of minor increases in the variance of all vital rates lead to a decrease in λ*_s (Haridas & Tuljapurkar, 2005)_*. In turn, to allow for the log_10_-transformation of *T*_σ_, *E*_σ*J*_^σ^*, E*_γ_^σ^, *E*_σ*A*_^σ^ and *E*_φ_^σ^, we used absolute values for comparative analysis as in Santos et al. (2023). Consequently, the absolute-value transformation yields values that negatively relate to the degree of demographic buffering, with higher values of *T*_σ_ and |*E* ^σ^| identifying less buffered populations.

### Sociality classification

To classify the degree of sociality across the 66 animal species included in this study, we employed a five-level sociality continuum previously introduced and justified by Salguero-Gómez (2024). This continuum captures gradations in social organisation across taxa by integrating spatial cohesion, temporal stability, and the frequency of social interactions. The five ordinal levels are: (1) Solitary: individuals live alone except during brief breeding encounters (*e.g*., tigers, some wasps); (2) Gregarious: individuals form temporary or fluid groups but engage in limited social coordination (*e.g.*, wildebeest, schooling fish); (3) Communal: individuals cohabit or share nesting sites but do not engage in cooperative care (*e.g.*, purple martins, some reef fish); (4) Colonial: individuals consistently share nesting or living spaces, often in dense aggregations (*e.g.*, seabirds, coral polyps); and (5) Social: individuals form stable, cooperative groups with persistent social bonds and behavioural coordination, including cooperative breeding or hierarchical organisations (*e.g*., meerkats, female elephants, baboons). These categories were designed to be taxonomically agnostic, allowing consistent application across a wide range of animal taxa.

We assigned sociality scores based on species-level information gathered from a combination of curated databases (*e.g*., Animal Diversity Web (Dewey et al., 2010), FishBase (Froese & Pauly, 2002), IUCN Red List (Gearty & Chamberlain, 2025)), peer-reviewed literature, and expert consultation. When available, we relied on published ethological reviews and species-specific studies documenting group size, social behaviour, and breeding systems. Each classification was independently validated by at least one taxonomic expert. In cases of ambiguity, we adopted a conservative approach by assigning the species to the lower of two adjacent categories unless strong evidence supported otherwise. See Salguero-Gómez (2024) and Table S2 for further details on our scoring criteria.

### Adult body mass

As life history traits, vital rates, and demographic buffering metrics often covary allometrically with body size (Calder 1984; Charnov 1993; Blueweiss et al. 1978), we explicitly discounted the effect of body mass in our phylogenetic comparative models. For each species in our dataset, we obtained adult body mass estimates from curated trait databases specific to major vertebrate and invertebrate taxa. Mammalian and avian body mass data were extracted primarily from the AnAge (de Magalhães & Costa, 2009) and AVONET databases (Tobias et al., 2022), respectively, while data for reptiles, amphibians, and fish were sourced from the Amniote Database (Myhrvold et al., 2015) and FishBase (Froese & Pauly, 2002), the latter using the rfishbase R package (Boettiger et al., 2012). For invertebrates and less-studied groups, we relied on MOSAIC (Bernard et al., 2023), data from Healy et al. (2019), and primary literature searches via Web of Science using species names and the keywords “adult body mass” or “adult weight.” All mass values were converted to grams and log_10_-transformed prior to analysis to meet assumptions of linearity and homoscedasticity. When multiple mass estimates were available for a species, we took the mean of adult female body mass when reported, as this tends to better reflect demographic contributions in iteroparous animals (J.-M. Gaillard et al., 2005; Isaac et al., 2007).

### Phylogenetic tree

To account for non-independence due to shared evolutionary history, we constructed a phylogenetic tree spanning all 66 species in our dataset using the Open Tree of Life as implemented in the rotl R package (Michonneau et al., 2016). First, we matched species names to OTL taxonomy using the ‘tnrs_match_names’ function, resolving any synonyms or misspellings manually. Using the matched taxon IDs, we retrieved a synthetic, ultrametric phylogeny. This resulting tree incorporates curated backbone information from published phylogenies across major animal clades. Branch lengths were scaled using divergence time estimates where available in the OTL backbone. For compatibility with downstream comparative models, we ensured the tree was fully bifurcating and resolved any polytomies using the ‘multi2di’ function from the ape R package (Paradis et al., 2004).

For species represented by multiple populations (n = 12 species, Table S2), we modified the tree above to create population-level phylogenetic tips by duplicating the species-level branch and assigning each population a unique identifier (e.g., *Species_1_Pop_1*, *Species_1_Pop_2*). These within-species duplicates were assigned to nearly zero-length terminal branches, thereby preserving the species-level topology and divergence times while accommodating population-level replication in the data. This approach allowed us to test hypotheses using the full population-level dataset (n = 87 populations) while still accounting for phylogenetic structure at the species level. The order of introduction of the populations within the tip of each species does not affect assessments in macroecological studies using COMADRE (Merrien et al., 2021). The resulting population-expanded tree was used for all comparative analyses, including phylogenetic ANOVAs, PGLS models, and estimation of phylogenetic signal (Pagel’s λ), using the R packages caper (Orme et al., 2013), nlme (Heisterkamp et al., 2017), and phytools (Revell, 2012).

### Climatic data and environmental stochasticity

To assess whether (H4) the demographic buffering effects of sociality vary across environments with differing climatic regimes, we obtained long-term climate data for each population using the NASA POWER (Prediction of Worldwide Energy Resources) database (*NASA Prediction Worldwide Energy Resources (POWER) Climate Data: Data Access Viewer*, 2020). This resource offers high resolution climatic products at approx. 4 km^2^ resolution from January 1958. For each of the 87 populations in our dataset, for which we have GPS coordinates in COMADRE, we extracted monthly mean temperature (T2M) and total precipitation (PRECTOTCORR) over the duration of the corresponding demographic time series and the 30 years preceding the start of the study, using the nasapower R package (Sparks, 2018). We then computed interannual summary statistics for each location and variable, including mean, maximum, minimum, and variance of precipitation and temperature, which formed the basis for estimating environmental predictability. Here, in regards to aquatic species, it is important to note that, even though species such as fish and corals are buffered on the short-term from fast fluctuations in precipitation, this abiotic factor remains a strong proxy for key hydrological processes—such as stream flow, turbidity, nutrient input, and water temperature, which in turn directly influence vital rates in freshwater and coastal environments (Buisson et al., 2008; Comte & Grenouillet, 2013; Fabricius, 2005).

We calculated three standardised metrics of climatic predictability following Colwell’s framework (1974), adapted for time series climate data: constancy, contingency, and predictability. Constancy quantifies the extent to which a climatic variable remains stable over time (1 - variance/mean), while contingency reflects the extent to which fluctuations are structured and recurrent. Predictability is then defined as the sum of constancy and contingency. These metrics were calculated separately for our records of monthly temperature and precipitation. We merged these values with each population’s demographic data, allowing us to explore how environmental predictability modulates the relationship between sociality and demographic buffering.

To construct a parsimonious and interpretable climatic PCA, we first reduced collinearity among the aforementioned environmental variables. Starting with a comprehensive set of temperature and precipitation-derived metrics (e.g. mean, variance, predictability), we computed all pairwise Spearman correlation coefficients to identify highly correlated variables. We visualised these correlations using custom pairwise scatterplots (Figure S4) and used a threshold of |ρ| > 0.85 to flag and remove strongly collinear variables using the function ‘findCorrelation’ in the caret R package (Irizarry, 2019). This threshold balances the need to retain meaningful ecological variation while minimizing redundancy and inflation of variance along principal component axes (Dormann et al., 2013). This step dropped off the following variables from our next steps: contingency and predictability of temperature, as well as constancy and contingency of precipitation. Next, for the climatic variables that we retained (mean, variance, and constancy of temperature, as well as mean, variance, and predictability of precipitation) we conducted a PCA using the ‘prcomp’ function in base R, centering and scaling all variables. We retained only the first two principal component (PC) axes, as only they raised associated eigenvalues > 1 (Figure S5) in agreement with Kaiser’s criterion (Legendre & Legendre, 2012).

This multivariate approach revealed how our 87 examined populations across 66 animal species are organised along two axes of climatic variability. The climatic principal component analysis reveals two axes (Figure 2) whose associated eigenvalue > 1 (Figure S5), and thus we retained these axes to test (H4) that the relationship between degree of sociality and demographic buffering would break apart in more stochastic environments. The first principal component (PC1), which explains 51% of the variance, positions populations along a continuum of climates with more variable precipitation (right) *vs.* more predictable precipitation (left). The second principal component (PC2), explaining 18% of variance, places populations along a continuum of higher mean annual precipitation (bottom) *vs*. climates with higher constancy in temperature (top).

**Figure 1.**
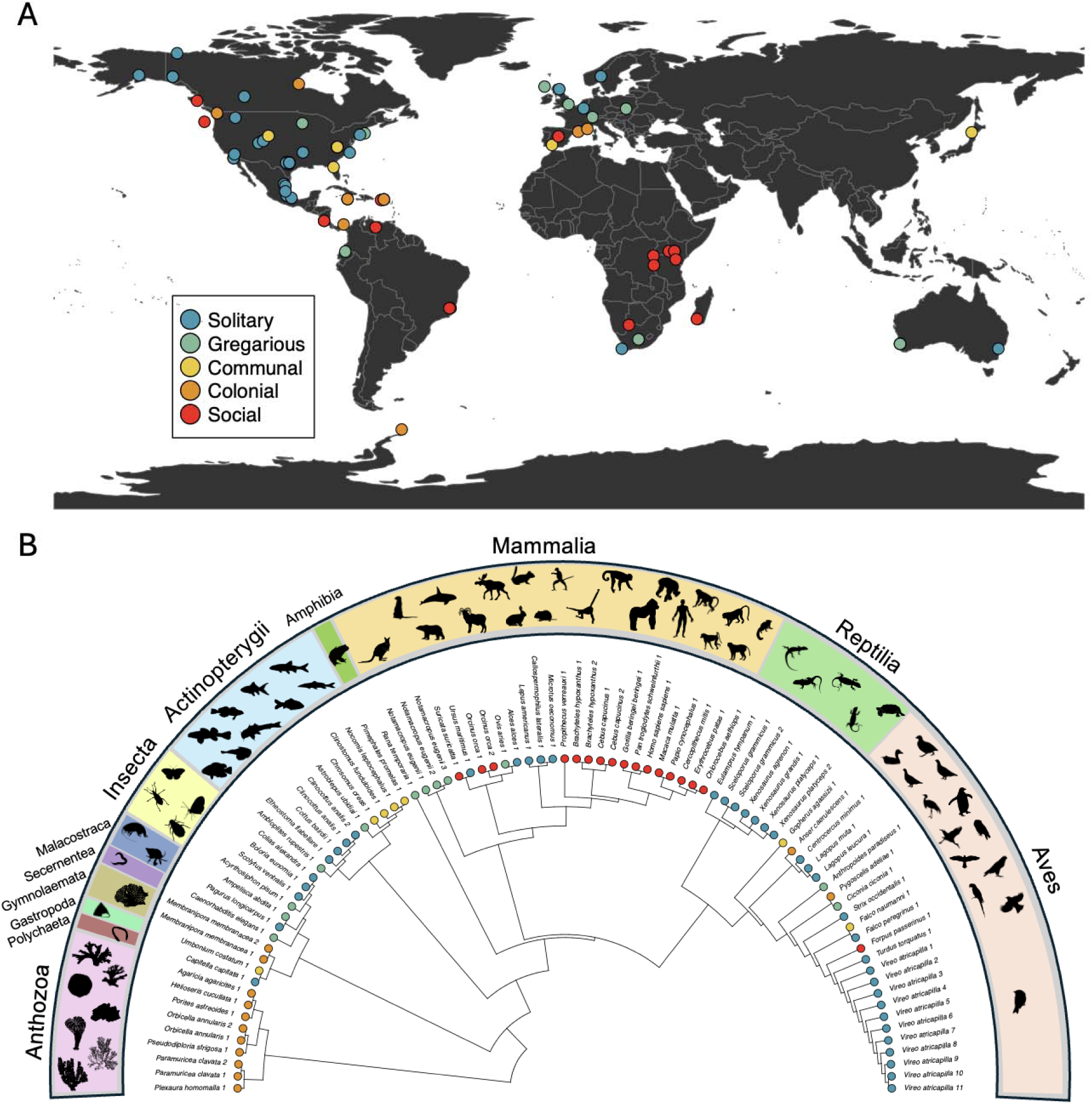
Representation of the 87 natural populations of the 66 animal species examined in this study to test the social buffering hypothesis. **A**. Geographic location of the populations. B. Phylogenetic relationships of the species. The phylogeny was modified to accommodate intra-specific spatial replication, indicated by the name of the species followed by the number of the population. In total, we examined patterns of demographic buffering across 12 animal taxonomic classes. Each species was classified into one of the five levels of sociality shown in the insert in panel A, following Salguero-Gómez (Salguero-Gómez, 2024).

**Figure 2.**
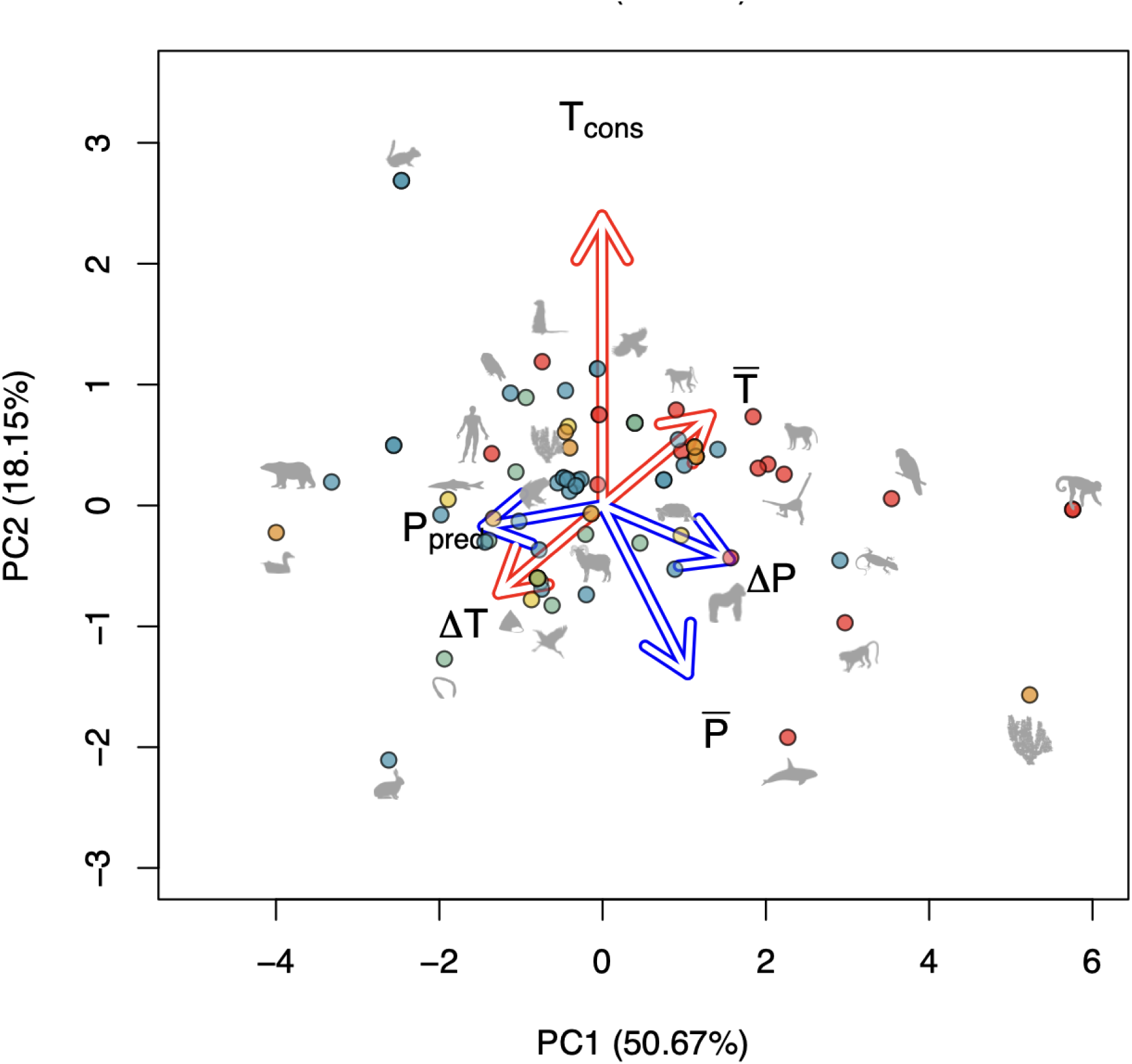
The examined 87 animal populations are organised along two axes of climatic variation: (PC1) predictability in precipitation and (PC2) constancy of temperature. Principal component analyses of the first two principal components, displaying the loadings of climatic products derived from the NASA POWER database. Climatic variables are: (1) Mean annual total precipitation (P^-^), (2) Variance annual total precipitation (Δ*T*), (3) Predictability of annual precipitation (*P_pred_*), (4) Mean annual temperature (T), (5) Variance annual temperature (Δ*T*); and (6) Constancy of temperature (*T_cons_*). The first principal component (PC1), which explains 50% of the variance, positions populations along a continuum of climates with more variable precipitation (right) *vs.* more predictable precipitation (left). The second principal component (PC2), explaining 18% of variance, places populations along a continuum of higher mean annual precipitation (bottom) *vs*. climates with higher constancy in temperature (top). Constancy quantifies the extent to which a climatic variable remains stable over time. Predictability is the sum of constancy and contingency, where contingency reflects the extent to which fluctuations are structured and recurrent. The colours of each dot represent the classification of sociality (See Figure 1). Silhouettes represent a subset of the animal species shown on the PCA, obtained from phylopic (Keesey, 2020).

### Comparative phylogenetic models

To test our four main hypotheses, we implemented a suite of phylogenetically-informed comparative models that account for the shared evolutionary history of the species in our dataset. As a first step, to discount the effect of adult body mass on the stochastic elasticities, a standard approach in comparative demography (J. M. Gaillard et al., 1989; Healy et al., 2019), we constructed a comparative data object using the ‘comparative.data’ function from the caper R package (Orme et al., 2013) to obtain the residuals of the phylogenetic generalised least squares (PGLS) models between each of the stochastic elasticities (Total: *T*, *T* ; and vital rate specific: |*E*_vr_ ^μ^|, |*E* ^σ^|) and body mass. We then used these residuals for the next analytical steps.

To evaluate H1, which posits that more social species exhibit stronger demographic buffering, we examined variation in the sum of stochastic elasticities to variance (*T*_σ_) and in the coefficients of variation of individual vital rates. We ran phylogenetic ANOVAs with sociality (ordinal, 1–5) as the predictor and each buffering metric as a response. These phylogenetic ANOVAs were implemented by fitting linear models followed by post hoc Tukey HSD tests, and their outputs were interpreted within a phylogenetic context based on trait alignment to the tree. To ensure results were phylogenetically robust, we repeated these tests in a PGLS framework, comparing results across both model types.

To test H2, which predicts that demographic buffering is primarily driven by adult vital rates, we used both phylogenetic ANOVAs and PGLS models with stochastic elasticities of the stochastic population growth rate to changes in the temporal variance of specific vital rates, separately. Specifically, we modelled the interannual CV of juvenile survival (|*E*^σ^ |), maturation (|*E*^σ^ |), adult survival (|*E*^σ^ |), and reproduction (|*E*^σ^ |) separately, using the degree of sociality as a predictor. Following significant ANOVA results, we carried out post-hoc Tukey test comparisons to determine which sociality levels differed most in their buffering patterns. By combining these models with the analysis of stochastic elasticities to variance (*E*^σ^) by life cycle stage, we further tested whether adult rates were the primary targets of buffering in more social taxa.

For H3, which posits that social species exhibit greater stochastic elasticity to the mean of adult vital rates, we analysed both total elasticity to the mean (*T*_μ_) and vital-rate-specific (|*E_vr_*^μ^|) values as response variables. We used PGLS models with the degree of sociality as predictor, and again followed up with phylogenetic ANOVAs and Tukey tests where model residuals supported group-wise comparisons. This allowed us to assess whether more social species rely more heavily on maximising the mean of critical demographic parameters—particularly adult survival and reproduction—rather than buffering their variance compared to less social species.

To test H4, that social buffering is weaker in highly stochastic environments, we extended the above model. Specifically, we ran two separate models evaluating the interaction between the degree of sociality and each of the two principal component axes of our climatic PCA above—our proxy for environmental stochasticity. These were implemented in PGLS models where response variables included either *T* or |*E* ^σ^|. We centred and scaled both predictors and examined interaction terms to assess whether the buffering benefit of sociality diminished as environments became more variable. We also conducted model comparisons with and without interaction terms using AIC to evaluate support for climate-modulated effects. A significant interaction term between the degree of sociality and a climatic driver would indicate that any potential relationship between demographic buffering and sociality is indeed modulated by the climate. A positive effect of this interaction would imply that climatic extremes render the demographic buffering of more social species less effective, as demographic buffering is highest when its symbol is smallest.

Across all analyses, we tested assumptions of model normality, checked for heteroscedasticity, and confirmed the presence of phylogenetic signal in the residuals. When strong effects were detected, we visualised results using standardised model coefficients and plotted predicted means from Tukey post-hoc tests to highlight contrasts across the sociality continuum. This multi-model approach ensured that our conclusions were robust to model structure and phylogenetic correction, and allowed us to rigorously test how sociality interacts with demography and environment across animal taxa.

## Results

Our results support the hypothesis (H1) that more social species exhibit enhanced demographic buffering compared to more solitary species. Indeed, more social species show a lower stochastic elasticity to vital rate variance (*T*_σ_) (Figure 3A). We found significant differences in *T*_σ_ across the sociality continuum, with social species exhibiting the lowest *T*_σ_ values, followed by communal, and then colonial, gregarious, and solitary (P = 0.0049). Solitary species consistently showed the highest sensitivity to changes in the variance of all vital rates.

**Figure 3.**
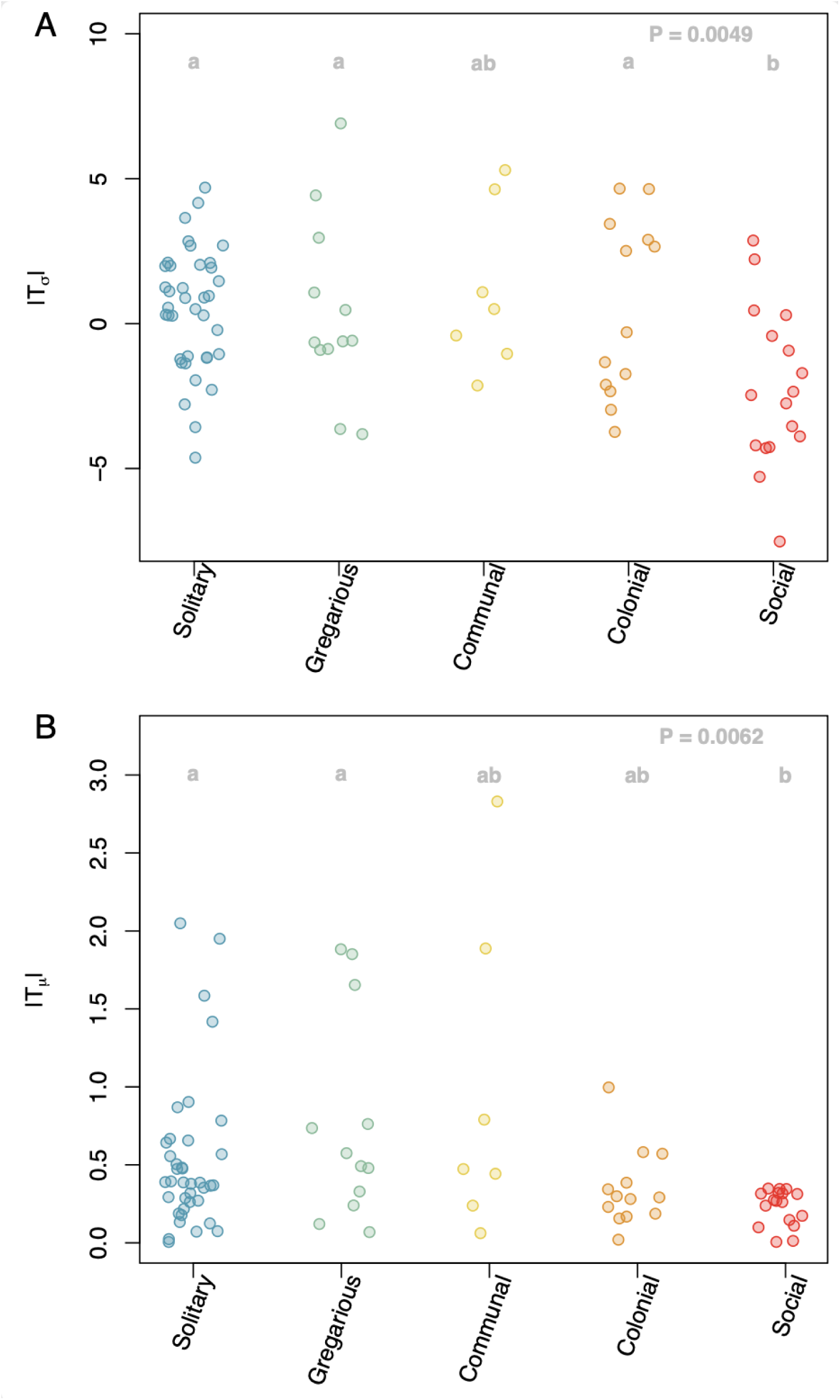
Increases in sociality are associated with canalisation in the selective pressures of demographic performance and increases in demographic buffering. Sum of stochastic elasticities of population growth rate (λ*_s_*) to changes in the (**A**) variance (*T*_σ_) and (**B**) mean (*T*_μ_) of all vital rates in a given population, classified by degree of sociality. Response variable is the absolute value of the residuals of *T*_σ_ and *T*_μ_ against adult body mass of each animal species. *T*_σ_ is log_10_-transformed. P values (top-right) correspond to a phylogenetic ANOVA, and post-hoc Tukey test letters positioned on top of each group indicate whether groups are significantly different, after phylogenetic corrections.

We found partial support for the prediction (H2) that adult stages drive demographic buffering patterns in more social species. Across the full dataset, stochastic elasticities to the variance of reproduction (|*E*^σ^ |) were consistently lower in more social species (Figure 4D, P = 0.0012). However, we found no significant difference across our social continuum regarding the stochastic elasticity to the variance of adult survival (|*E*^σ^ |; Figure 4C, P = 0.6666). Unexpectedly, we found a lower value of the stochastic elasticity to variance in juvenile survival (|*E*^σ^ |) in more social species compared to less social ones (Figure 4A, P = 0.0128), implying that more social species buffer more against stochastic environments in early stages of their development. The stochastic elasticity to variance in maturation (|*E*^σ^ |) showed no significant differences across sociality groups.

**Figure 4.**
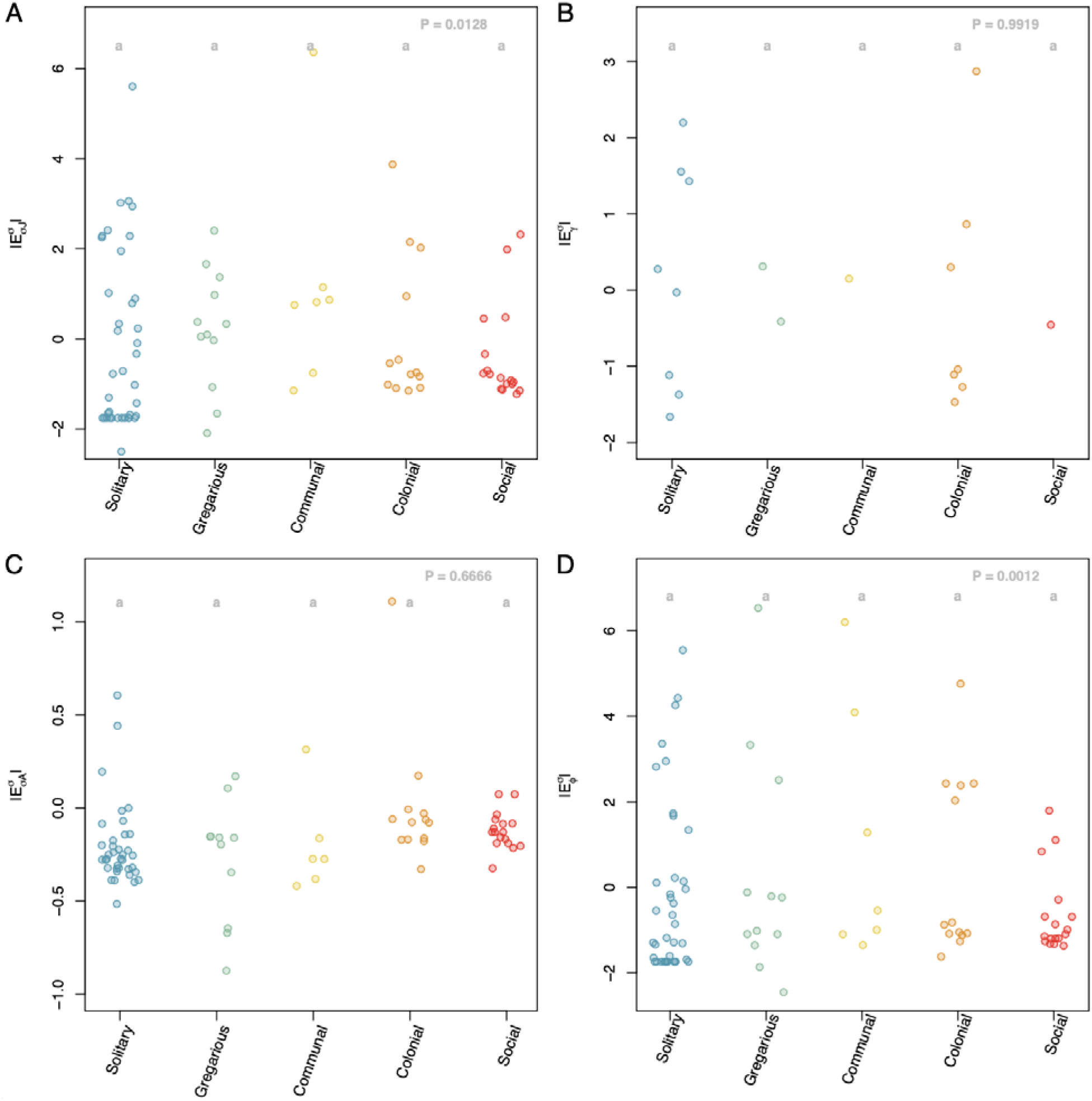
As species become more social, their population performance becomes less sensitive to changes in juvenile survival and reproduction. Stochastic elasticity of population growth rate (λ*_s_*) to changes in the variance of four vital rates (*vr*) (|*E_vr_*^σ^|): (|*E_vr_*^σ^|): juvenile survival (|*E* ^σ^|), maturation (|*E* ^σ^|), adult survival (|*E* ^σ^|), and reproduction (|*E* ^σ^|), grouped by degree of sociality. Response variables are the residuals of *E* ^σ^ against adult body mass of each animal species, and log - transformed. P values correspond to a phylogenetic ANOVA, and post-hoc Tukey test letters positioned on top of each group indicate whether groups are significantly different, after phylogenetic corrections.

We found support for our hypothesis (H3) that more social species invest more in maximising the mean of adult vital rates than more solitary species. Indeed, total stochastic elasticity to the mean values (|*T*_μ_|) differ significantly across sociality levels (P = 0.0062; Figure 3B), with social species having the lowest values. The vital-rate-specific analyses also reveal that social species exhibit significantly higher stochastic elasticity values to changes in the mean of adult survival (|*E*^μ^ |; Figure 5C, P = 0.0017) and reproduction (|*E*^μ^ |; Figure 5D, P = 0.0125). However, the stochastic elasticity to mean juvenile survival is also lowest in more social species (|*E*^μ^ |; Figure 5A, P = 0.0046). There is no effect for mean maturation on the stochastic population growth rate (λ*_s_*) (|*E*^μ^_γ_|; Figure 5B, P = 0.5481). The differences in vital-rate specific stochastic elasticities are clearest between solitary and social species in post-hoc comparisons. These patterns imply that more social species achieve population stability not only by buffering variance, as discussed above, but also by enhancing the average performance of adult and juvenile vital rates.

**Figure 5.**
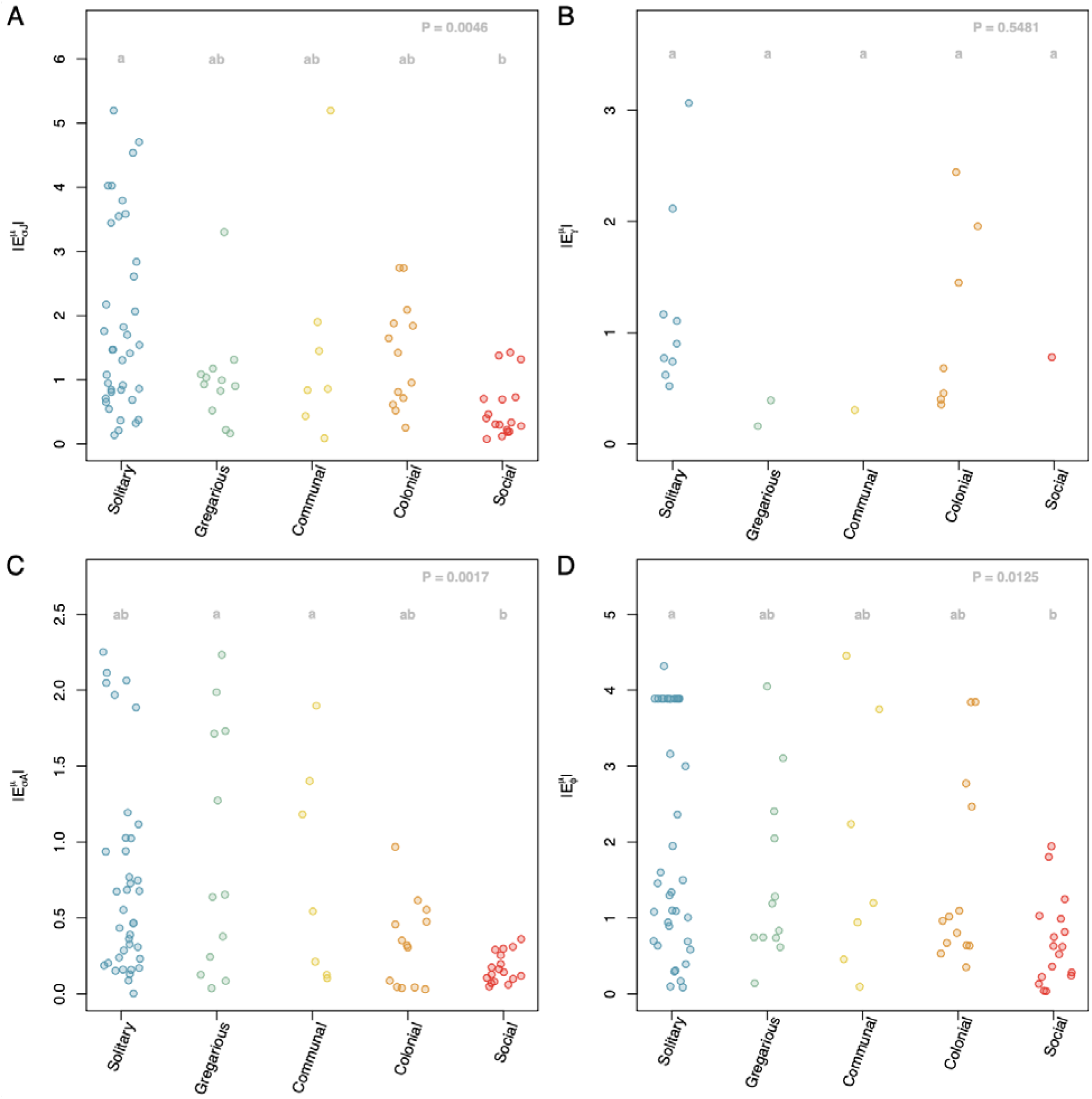
In more social species, vital rates are more tightly regulated. Stochastic elasticity of population growth rate (λ*_s_*) to changes in the mean of four vital rates (*vr*) (|*E_vr_*^μ^|): juvenile survival (|*E* ^μ^|), maturation (|*E* ^μ^|), adult survival (|*E* ^μ^|), and reproduction (|*E* ^μ^|), grouped by degree of sociality. Response variables are the absolute values of the residuals of *E_vr_*^σ^ against adult body mass of each animal species. P values correspond to a phylogenetic ANOVA, and post-hoc Tukey test letters positioned on top of each group indicate whether groups are significantly different, after phylogenetic corrections.

We found evidence that (H4) the benefits of social buffering are climatically-dependent. However, we did not find evidence to support that these potential benefits break down under highly unpredictable climatic regimes, but in fact increase non-linearly. We found a negative, statistically significant coefficient for the interaction between sociality and precipitation predictability (PC1 in Figure 2) in the PGLS model predicting *T*_σ_ (P = 0.027; Table 1A). This means that, as environments become less climatically predictable (left to right in PC1; *e.g*., more erratic rainfall), highly social species diminish their stochastic elasticity to variance even further compared to highly social species in more predictable environments, indicating enhanced demographic buffering in highly stochastic environments. The interaction between sociality and PC2 (Temperature constancy) did not significantly predict *T*_σ_ (Table 1). The fact that the positioning of solitary, gregarious, communal, colonial, and social species along this climatic PCA is not significantly associated with either axis (Table S3) indicates that this key finding is not driven by environmental filtering on each group in different regions of the climatic space.

**Table 1.** Populations located in climates with low predictability for precipitation regimes show a disproportionate increase in their demographic buffering abilities. Estimates and associated P-values (P < 0.05 in bold) for a battery of phylogenetic generalised least square (pgls) models evaluating the effects of the degree of sociality and climate on the elasticity of the stochastic population growth rate (λ*_s_*) to (**A**) the total change in mean and variance of all vital rates in each population, (**B**) changes in the mean of each vital rate, and (**C**) change in the variance of each vital rate, separately. PC1 and PC2 are defined in Figure 2, and correspond to a continuum of predictability in precipitation and of constancy of temperature, respectively. A significant interaction between sociality and either principal component axis indicates that the relationship between demographic buffering and sociality changes along climatic regimes.

| Stochastic elasticity<br>to |  | Predictor | Estimate | P | Predictor | Estimate | P |
| --- | --- | --- | --- | --- | --- | --- | --- |
| A. Total | Mean - $T_\mu$ | Sociality | 0.221 | 0.541 | Sociality | 0.052 | 0.878 |
|  |  | PC1 | 0.894 | 0.165 | PC2 | -0.268 | 0.770 |
|  |  | Soc x PC1 | -0.231 | 0.286 | Soc x PC2 | 0.22 | 0.472 |
| | Variance - $T_\sigma$ | Soc | 1.778 | **0.016** | Soc | 0.634 | 0.497 |
|  |  | PC1 | 4.233 | **0.026** | PC2 | -1.362 | 0.585 |
|  |  | Soc x PC1 | -1.431 | **0.027** | Soc x PC2 | 1.290 | 0.131 |
| **B.**<br>Changes<br>in mean | Juvenile<br>survival -<br>$\vert E_{\sigma J}^{\mu}\vert $ | Soc | 0.423 | 0.446 | Soc | 0.052 | 0.911 |
|  |  | PC1 | 0.768 | 0.424 | PC2 | -0.816 | 0.516 |
|  |  | Soc x PC1 | -0.411 | 0.216 | Soc x PC2 | 0.631 | 0.141 |
| | Maturation -<br>$\vert E_{\gamma}^{\mu}\vert $ | Soc | -0.503 | 0.538 | Soc | -0.724 | 0.337 |
|  |  | PC1 | 1.027 | 0.468 | PC2 | -1.278 | 0.524 |
|  |  | Soc x PC1 | -0.345 | 0.475 | Soc x PC2 | 0.551 | 0.408 |
| | Adult<br>survival -<br>$\vert E_{\sigma A}^{\mu}\vert $ | Soc | 0.183 | 0.751 | Soc | 0.028 | 0.960 |
|  |  | PC1 | 1.150 | 0.257 | PC2 | 0.032 | 0.983 |
|  |  | Soc x PC1 | -0.235 | 0.491 | Soc x PC2 | 0.069 | 0.889 |
| | Reproductio<br>n - $\vert E_{\phi}^{\mu}\vert $ | Soc | 0.382 | 0.656 | Soc | -0.216 | 0.778 |
|  |  | PC1 | 2.262 | 0.141 | PC2 | 0.216 | 0.916 |
|  |  | Soc x PC1 | -0.677 | 0.193 | Soc x PC2 | 0.482 | 0.481 |
| **C.** | Juvenile | Soc | 0.518 | 0.631 | Soc | 0.112 | 0.911 |
| Changes<br>in variance | survival -<br>$\vert E_{\sigma J}^{\sigma}\vert $ | PC1 | 1.791 | 0.342 | PC2 | -1.321 | 0.625 |
|  |  | Soc x PC1 | -0.607 | 0.346 | Soc x PC2 | 0.595 | 0.507 |
| | Maturation -<br>$\vert E_{\gamma}^{\sigma}\vert $ | Soc | -0.156 | 0.888 | Soc | -0.413 | 0.691 |
|  |  | PC1 | 1.790 | 0.356 | PC2 | -1.711 | 0.540 |
|  |  | Soc x PC1 | -0.497 | 0.451 | Soc x PC2 | 0.569 | 0.538 |
| | Adult<br>survival -<br>$\vert E_{\sigma A}^{\sigma}\vert $ | Soc | 0.283 | 0.384 | Soc | 0.150 | 0.613 |
|  |  | PC1 | 0.525 | 0.352 | PC2 | 0.151 | 0.849 |
|  |  | Soc x PC1 | -0.152 | 0.425 | Soc x PC2 | 0.056 | 0.832 |
| | Reproductio<br>n - $\vert E_{\phi}^{\sigma}\vert $ | Soc | 1.223 | 0.334 | Soc | 0.430 | 0.708 |
|  |  | PC1 | 3.401 | 0.129 | PC2 | -0.574 | 0.852 |
|  |  | Soc x PC1 | -0.976 | 0.197 | Soc x PC2 | 0.879 | 0.393 |

## Discussion

Understanding how sociality influences demographic performance under environmental stochasticity is essential for predicting population responses to ongoing global change. Although social behaviour is widely recognised as a key axis of animal life history (Clutton-Brock, 2009; Lukas & Clutton-Brock, 2012; Silk, 2007a), its role in modulating the demographic consequences of environmental variability remains poorly quantified across taxa. This gap is especially important given the increasing recognition that social traits can act as buffering mechanisms against ecological uncertainty (Albery et al., 2022; Doody et al., 2013; Rubenstein & Lovette, 2007). Our study provides the first macroecological and phylogenetically informed test of the social buffering hypothesis across the animal kingdom, using time-series demographic data from 87 populations of 66 animal species spanning 12 taxonomic classes. We find broad support for the idea that more social species exhibit stronger demographic buffering (H1), especially in adult stages (H2), and partially support the life history theory expectation that social species maximise mean demographic performance in adult vital rates (H3). Crucially, we show that these buffering benefits are conditional and become more important in highly stochastic environments (H4), thus adding a critical environmental contingency to the social buffering hypothesis.

Our results support the social buffering hypothesis (Rubenstein & Lovette, 2007), showing that population growth in more social species is less sensitive to interannual fluctuations in vital rates, as measured by lower total stochastic elasticity to variance (*T*_σ_; (Haridas & Tuljapurkar, 2005). This finding suggests that features of systems that are organised socially—such as alloparental care, food and information sharing, cooperative hunting, and defence—confer demographic stability by reducing the influence of environmental noise on fitness-related traits. These results echo and extend previous work in specific taxonomic groups (*e.g.*, birds: (Jetz et al., 2012); mammals: (Silk, 2007b); reptiles: (Doody et al., 2013)) and are consistent with theoretical predictions that selection should favour strategies that reduce the fitness costs of environmental variance (Hilde et al., 2020; Morris et al., 2008; Pfister, 1998). By demonstrating this pattern across 12 taxonomic classes, our findings indicate that the demographic benefits of sociality are widespread and likely not restricted to well-studied animal groups.

The ability of social species to buffer against environmental stochasticity is not uniformly distributed across their vital rates or life cycle stages. Rather, we provide evidence that their vital rates can exert different levels of demographic buffering. The most prominent buffering mechanism we show here in the context of sociality is related to the forces of natural selection that shape adult survival and reproduction through environmental canalisation. Our finding is consistent with theoretical predictions (Tuljapurkar, 1982) and empirical evidence (J.-M. Gaillard & Yoccoz, 2003; Morris et al., 2008; Santos et al., 2023) that traits contributing most to population growth rate are often the most canalised (*i.e.*, buffered). In this sense, it is not surprising that, in our study, juvenile survival and maturation show little evidence of buffering (*i.e*., greater values of |*E_vr_*^σ^|) in more social species, as juvenile vital rates are typically under weaker selection—and thus more variable—in long-lived species (McDonald et al., 2016; Stearns, 1976, 1999). Social systems likely reinforce this canalisation by promoting adult stability through mechanisms such as cooperative breeding, protection from predation, and shared resource acquisition (Bourke, 2011; Koenig & Dickinson, 2016).

In addition, we also provide evidence of a second buffering mechanism that acts on juvenile survival—where canalisation is less expected. Although juvenile survival is generally under weak natural selection, we show that increasing sociality significantly reduces the extent to which variation in this vital rate affects population growth. As sociality increases, juvenile survival becomes more buffered—a pattern not detected in vital rates that are already highly buffered, such as adult survival. This secondary buffering mechanism reinforces the idea that demographic buffering is life cycle stage dependent, but also challenges the assumption that demographic buffering occurs exclusively at vital rates where stability most enhances population growth (Hilde et al., 2020; Morris et al., 2008; Pfister, 1998).

In partial support of H3, we found that social species do not exhibit higher total stochastic elasticity to mean vital rates (*T*_μ_; (Haridas & Tuljapurkar, 2007) overall, but do show higher elasticity to the mean of adult survival and reproduction specifically compared to less social species. This finding suggests that, while sociality may not increase the overall importance of mean performance across all vital rates, it does amplify the role of key adult traits in determining population growth. This pattern is consistent with life-history theory predicting that species should maximise the performance of traits most important to fitness (Stearns, 1999). The elevated |*E*^μ^ | and |*E*^μ^ | values in more social species likely reflect adaptive strategies that enhance adult demographic performance in stable or cooperative contexts. However, this pattern did not hold across juvenile traits, again pointing to a stage-specific effect of sociality on demographic investment. Indeed, social buffering might be expected to disproportionately benefit adults over juveniles if social relationships establish over the life course <u>(Firth et al. 2024)</u>. Specifically, if adults typically hold stronger and more consistent social relationships compared to juveniles <u>(Woodman et al. 2024)</u>, this may enhance their ability to leverage group support under environmental stress. For instance, in cooperatively breeding mammalian species (such as meerkats, or baboons), dominant adult breeders often hold established social relationships to other group members and benefit from priority access to shared resources, reduced predation risk through increased vigilance, and assistance in offspring care; these mechanisms would all directly stabilise adult survival and fecundity more than they would affect juvenile traits (Clutton-Brock et al. 2016; Silk et al. 2016). Similarly, adult individuals in stable avian breeding groups often benefit from load-lightening (via helper contributions to territory defense, and joint nest provisioning), thus potentially reducing adult mortality and reproductive variability relative to juvenile birds, who are often less integrated into established social roles in cooperatively breeding systems (Rubenstein & Abbot 2017). These differences in social roles across the life-course suggest that demographic buffering by sociality might be particularly pronounced for adult vital rates.

Importantly, our results reveal that the demographic buffering effects of sociality are context-dependent and may be more important in highly stochastic environments (H4). This finding adds a novel macroecological insight to the social buffering hypothesis. Contrary to some work suggesting that demographic buffering mechanisms can break down under extreme environmental conditions (Rodriguez-Caro et al., 2021; Santos et al., 2024), our results indicate that sociality may become even more effective in buffering demographic processes against environmental stochasticity when conditions are highly unpredictable. This pattern echoes long-standing predictions about the evolution of cooperative systems in harsh environments, such as the development of eusociality in termites, (semi)social cockroaches, and naked mole rats, where group living is thought to have evolved in response to arid, variable conditions (Alexander, 1974; Faulkes & Bennett, 2013; Field & Toyoizumi, 2020; Lubin & Bilde, 2007). Rather than social organisation being strained or cooperative behaviours faltering under extreme variability (Hayes, 2017), our findings suggest that sociality can enhance demographic resilience precisely where environmental unpredictability is greatest. Thus, we highlight the importance of considering environmental context in assessments of social evolution and suggest that sociality may play a crucial role in mediating species’ demographic responses to ongoing climate change.

Together, our findings suggest that sociality promotes demographic stability by buffering the variance of adult vital rates, and that these benefits are environmentally dependent. In doing so, we bridge theoretical work on demographic buffering (Hilde et al., 2020; Pfister, 1998; Tuljapurkar, 1982) with recent calls to integrate social behaviour into eco-evolutionary frameworks (Albery et al., 2022; Firth et al., 2024; Salguero-Gómez, 2024). Our use of time-series demographic data across a broad range of taxa, combined with a structured classification of sociality and high-resolution climate products, offers a general framework to assess how social and environmental factors interact to shape population dynamics. In an era of increasing environmental unpredictability (Bathiany et al., 2018), understanding the limits of social buffering is vital for predicting which species are most vulnerable to demographic destabilisation— and why.

## Supporting information

SOM

## Acknowledgements

This work was supported by a NERC Pushing the Frontiers grant (NE/X013766/1) to RSG. Some ideas for this paper originated in discussions with colleagues during the Royal Society meeting “Understanding age and society using natural populations”.

## Author Contributions

RSG conceived the ideas and designed the methodological approach with input from KD and LB. RSG collected the pertinent data or accessed it via open-access repositories. RSG curates COMADRE. RSG performed the analyses with input from GS. RSG wrote the first draft with input from KD. All coauthors contributed to subsequent versions of the manuscript.

## Statement of inclusivity

The authorship is composed of researchers of all academic stages, genders, and several nationalities, including multiple languages.

## Supplementary Online Materials

**Table S1.**
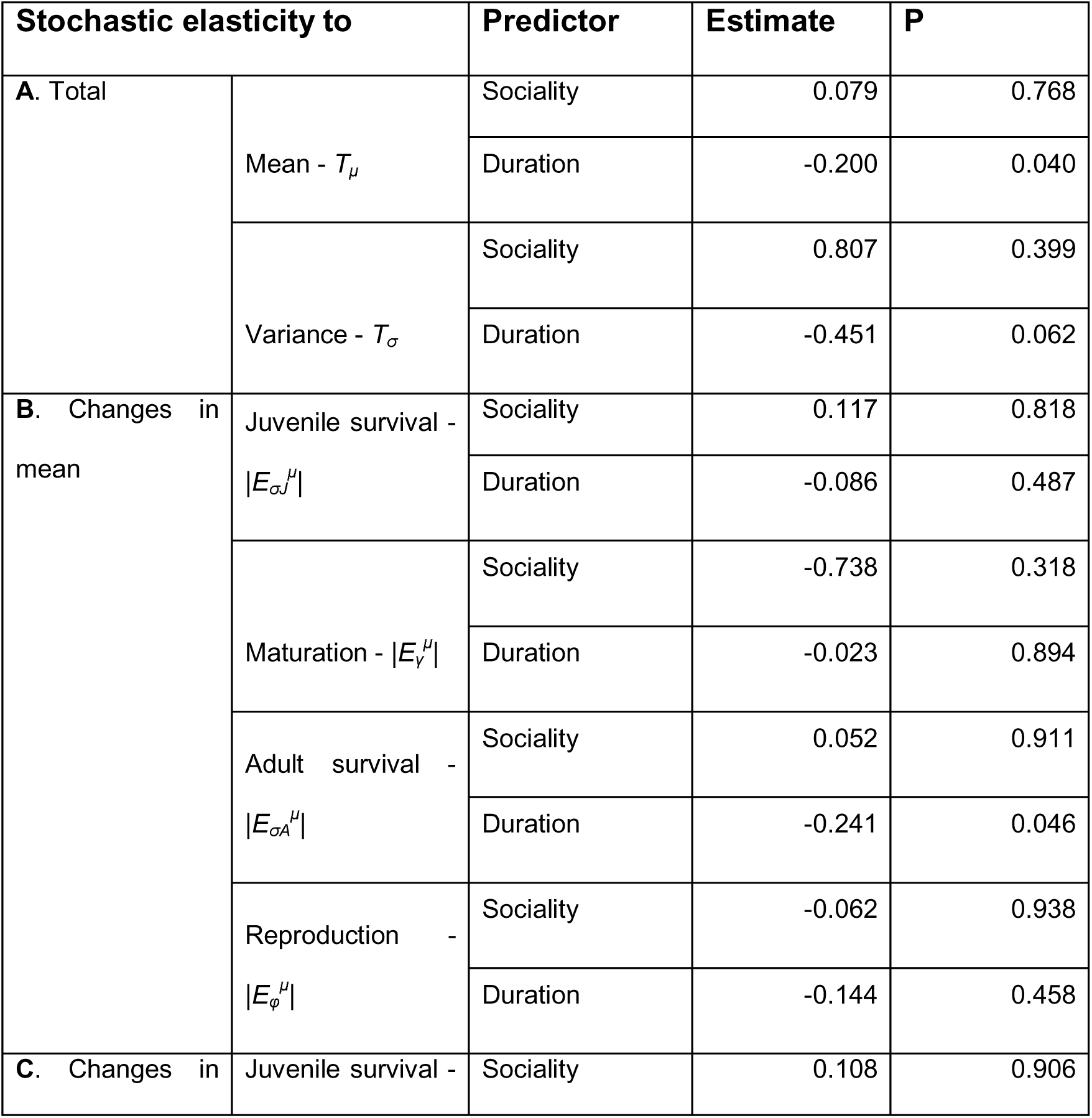
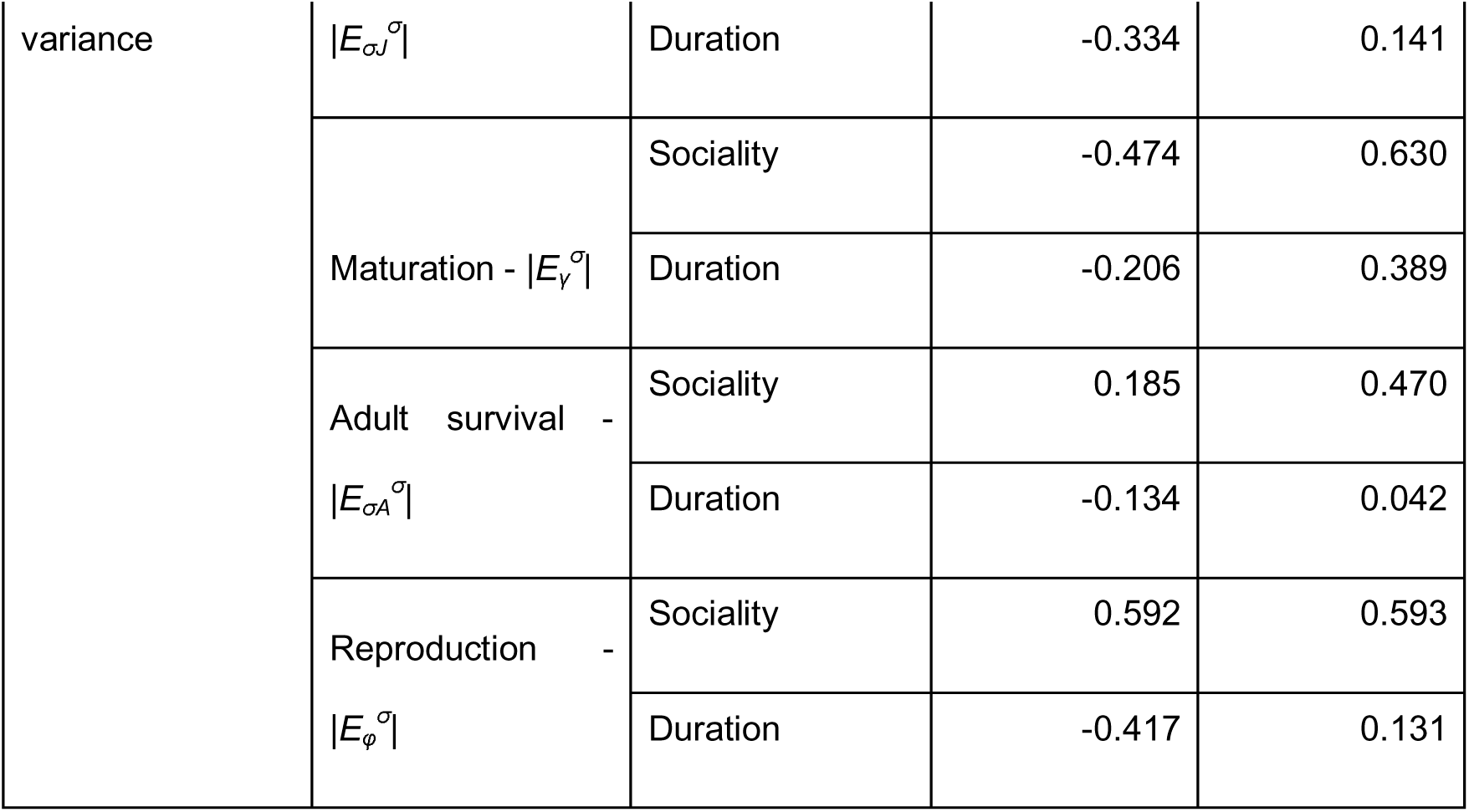
Sensitivity of results to temporal replication in studies Our overall results are mostly insensitive to the duration of the study. Here, study duration is defined as the number of matrix population models (MPMs) available in each examined population. Battery of pgls models examining the relationships between the different stochastic elasticities of stochastic population growth rate (λ*_s_*) and sociality, with study duration as a covariate. Note that the latter is only borderline significant in three occasions: *T*_σ_, |*E*_σ_*_A_*^μ^| and |*E*_σ_*_A_*^σ^|.

**Table S2.**
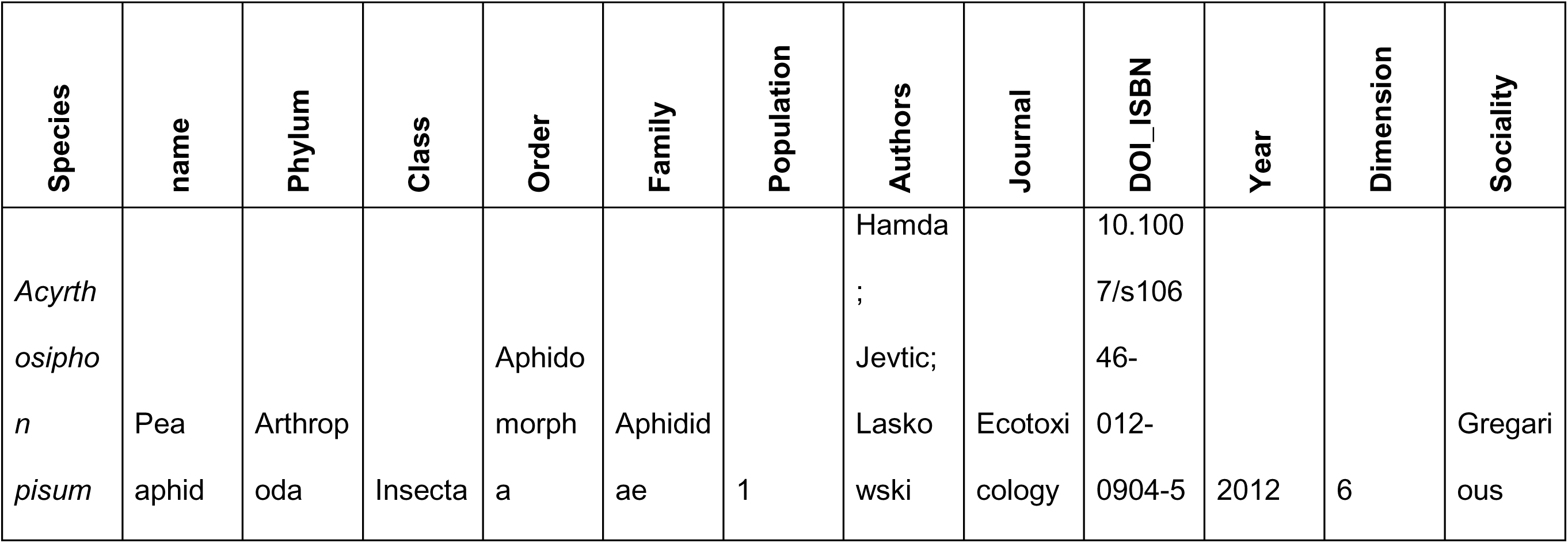
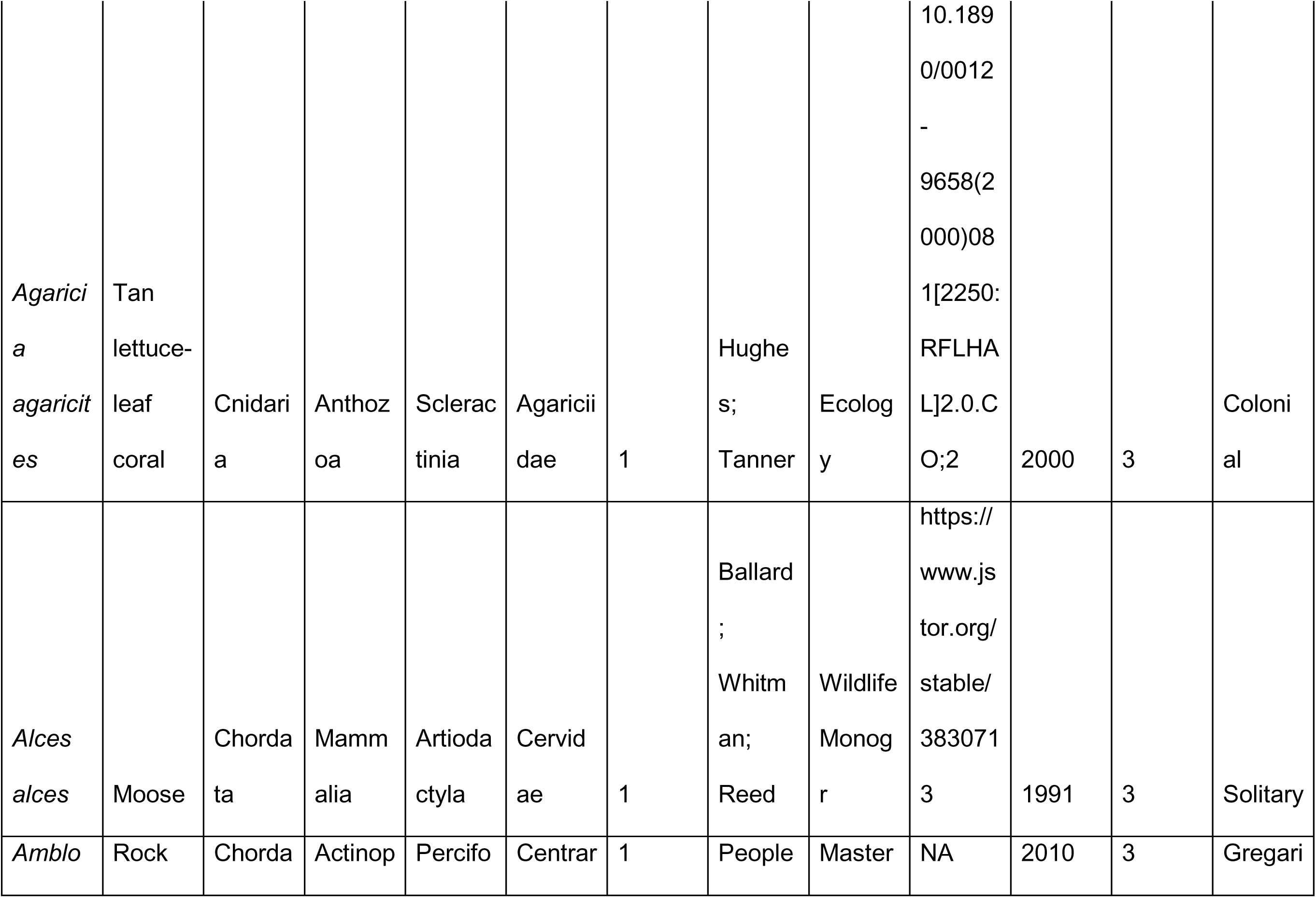
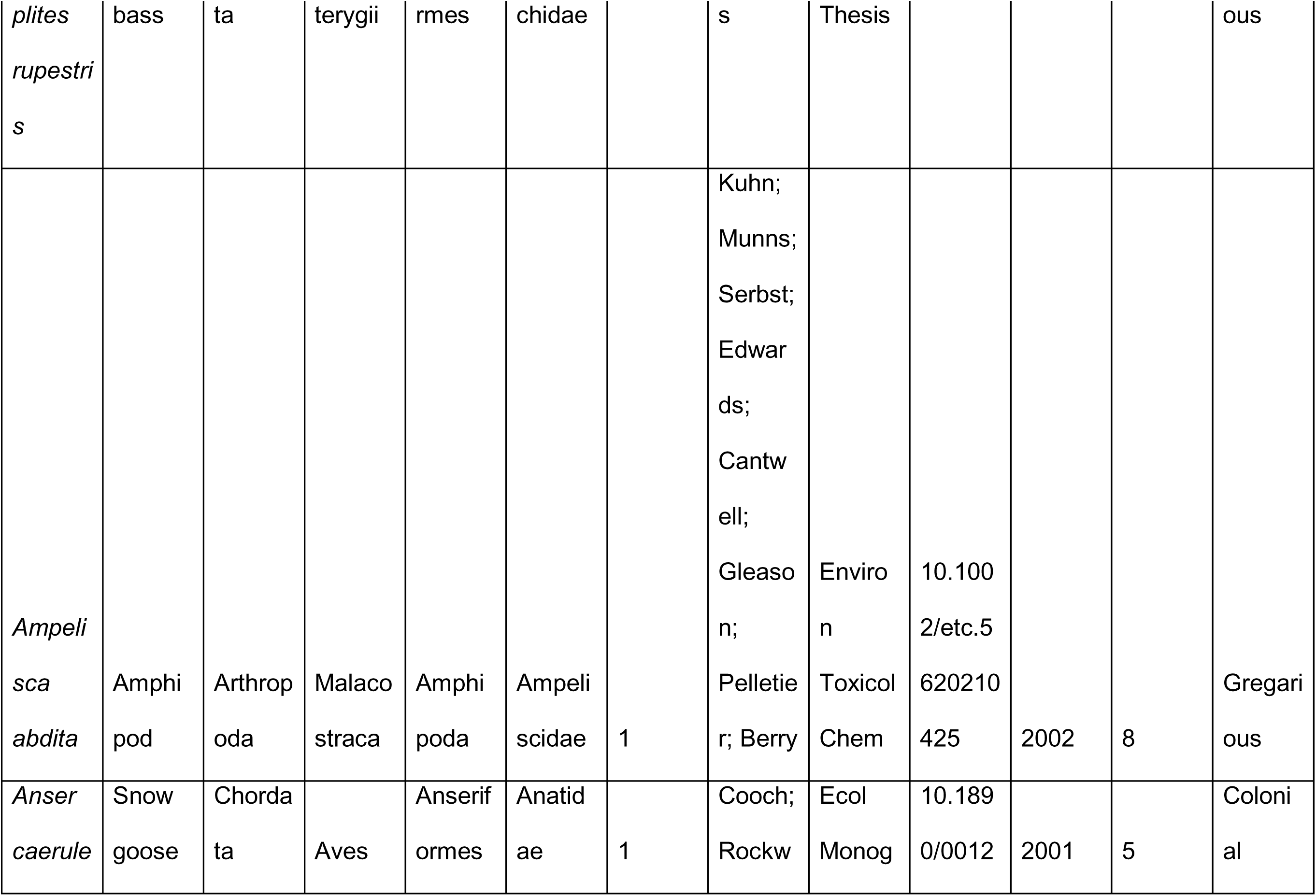
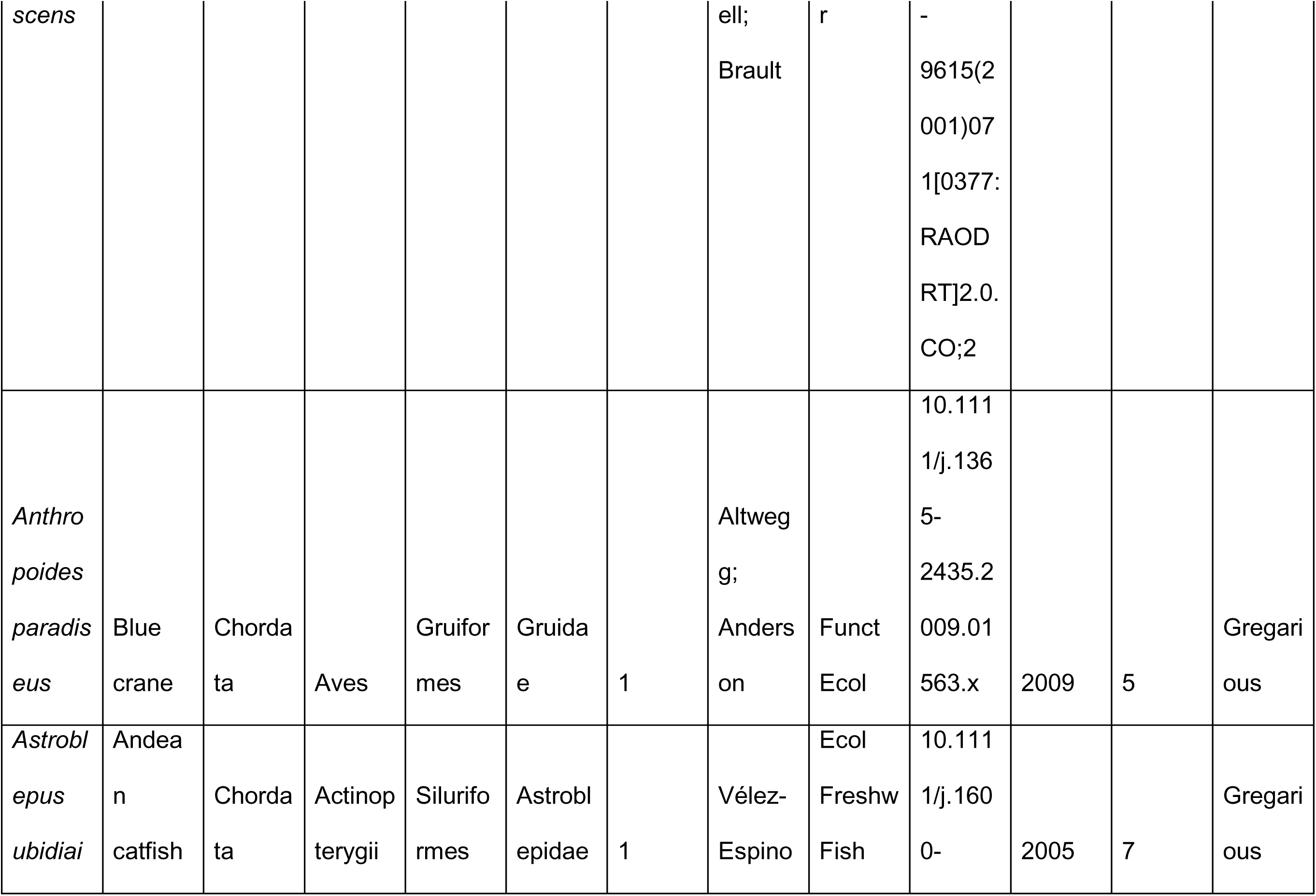
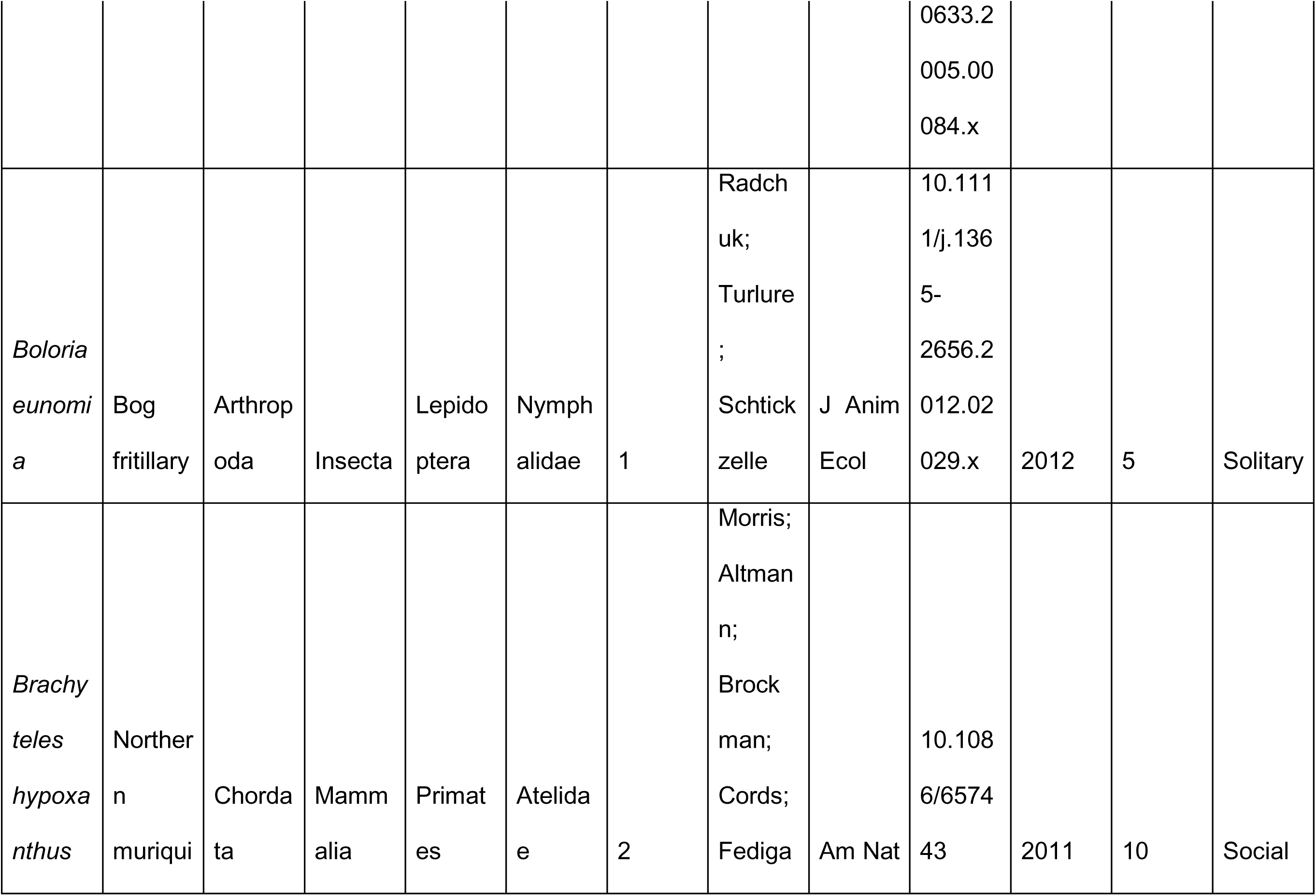
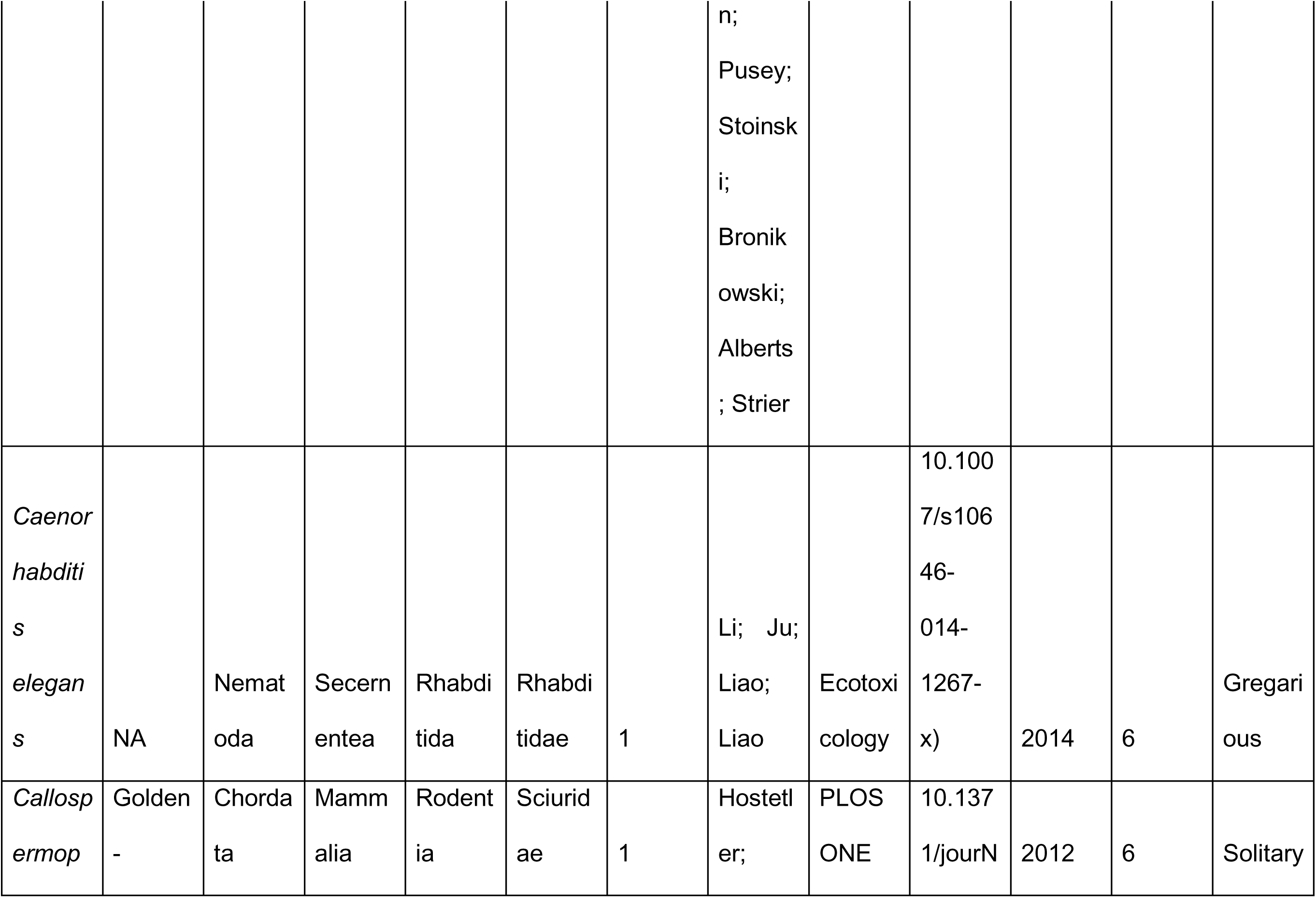
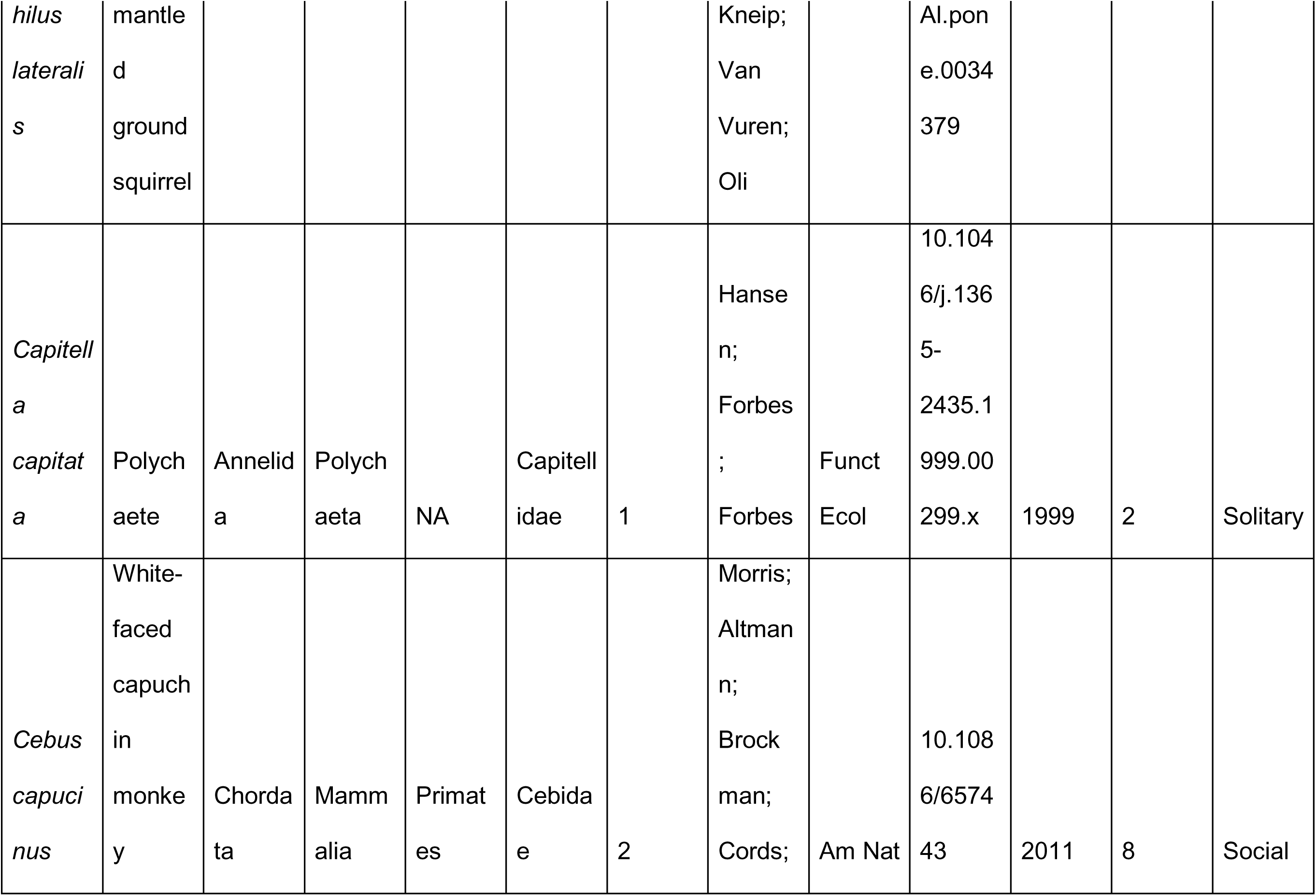
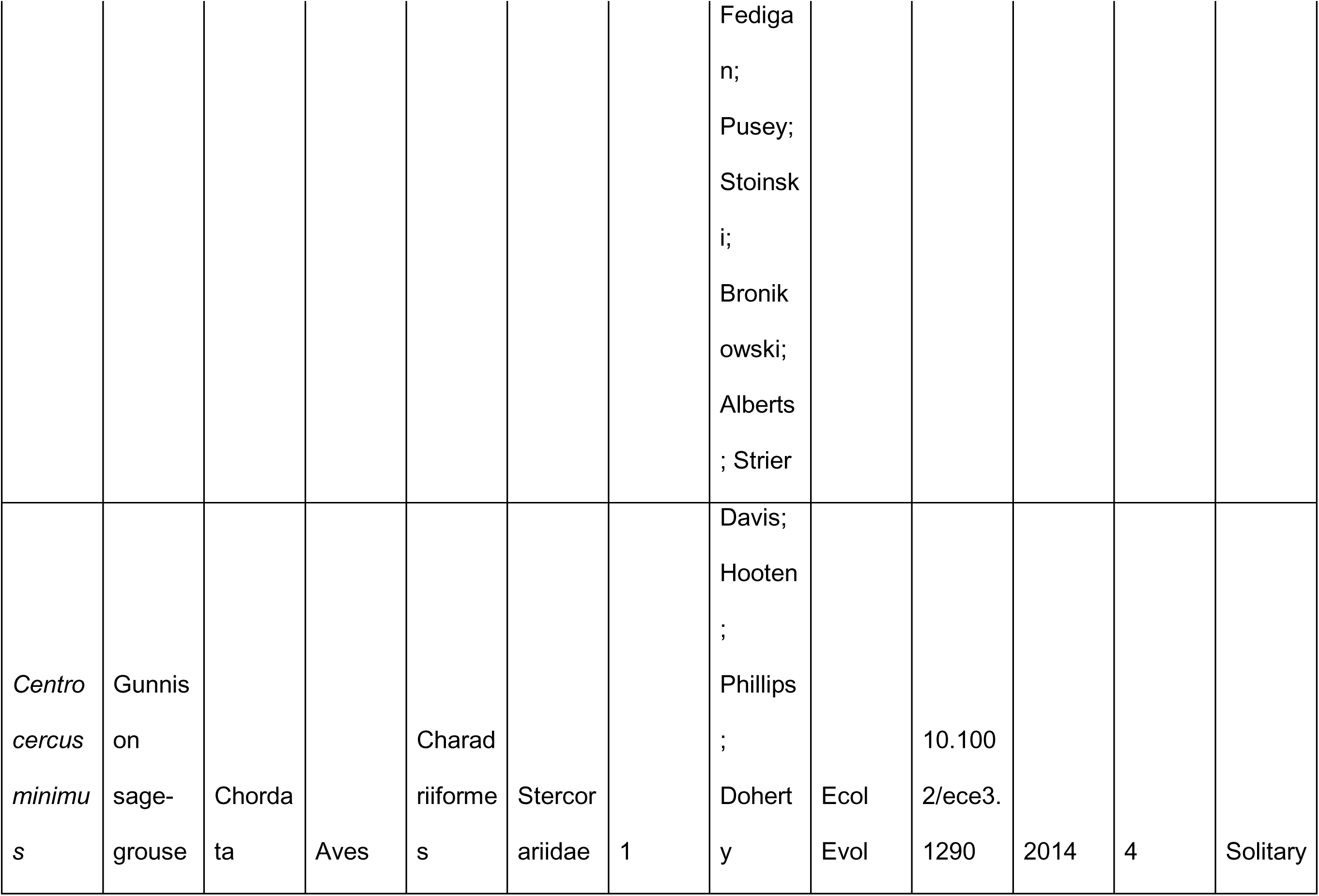
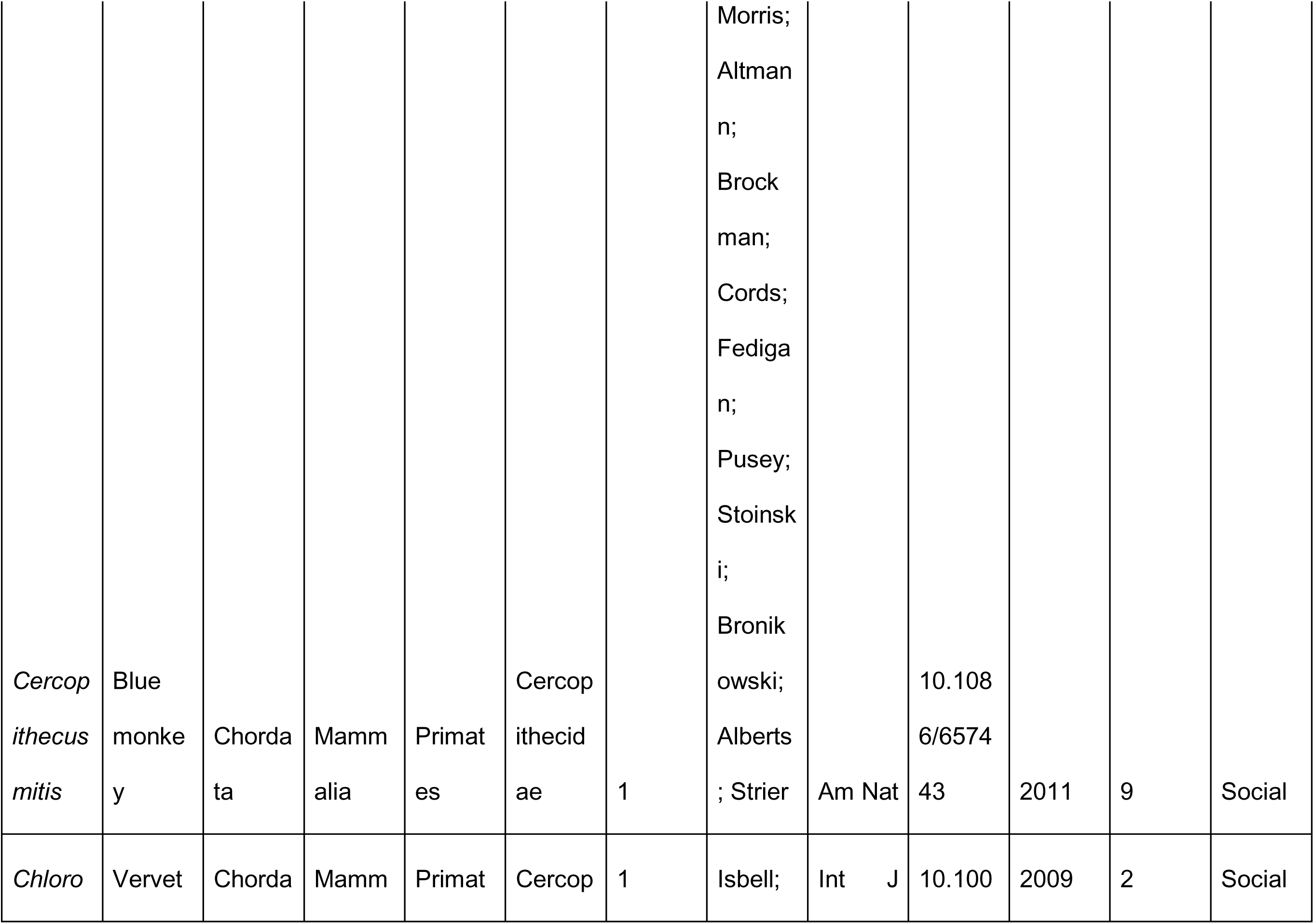
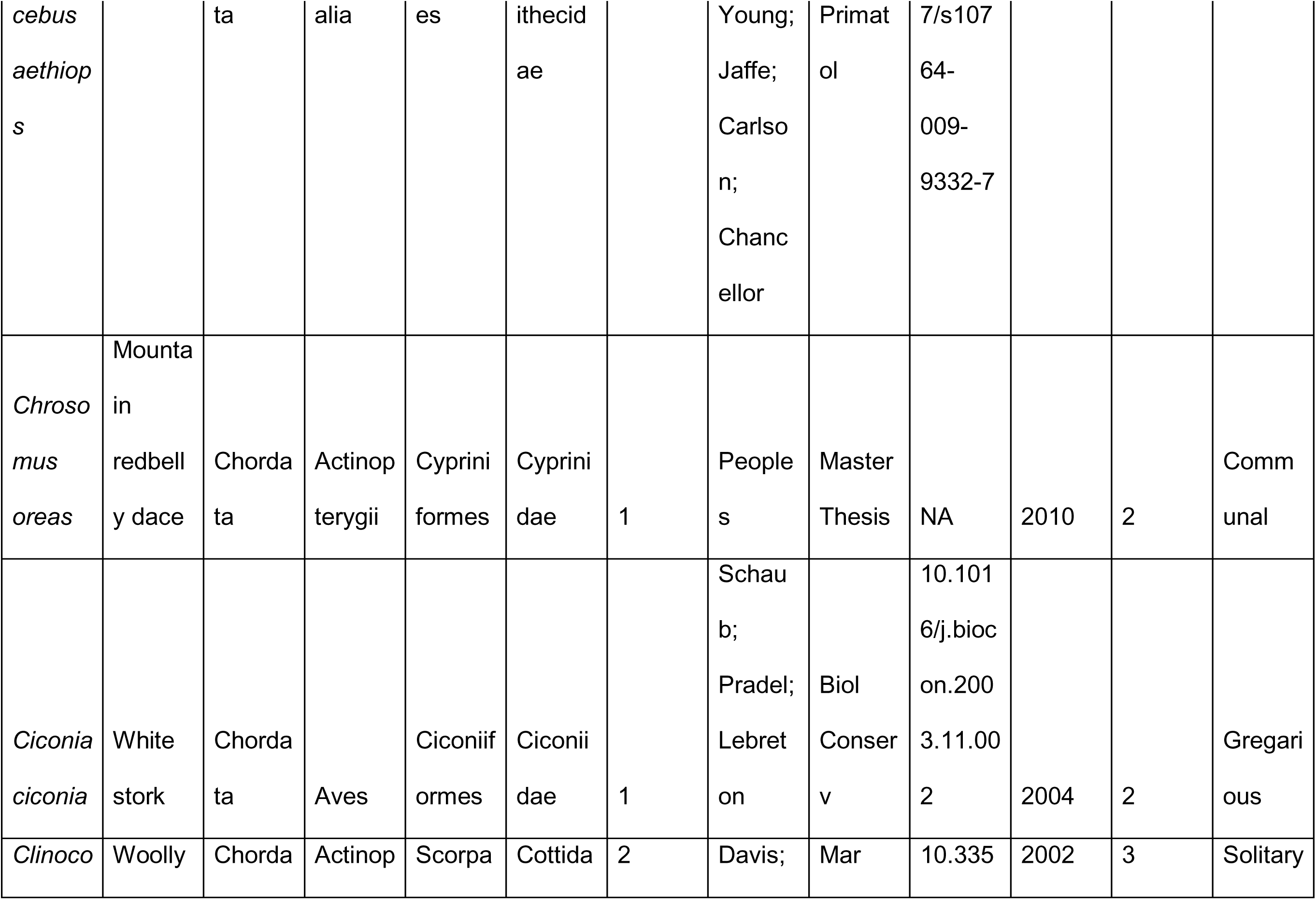
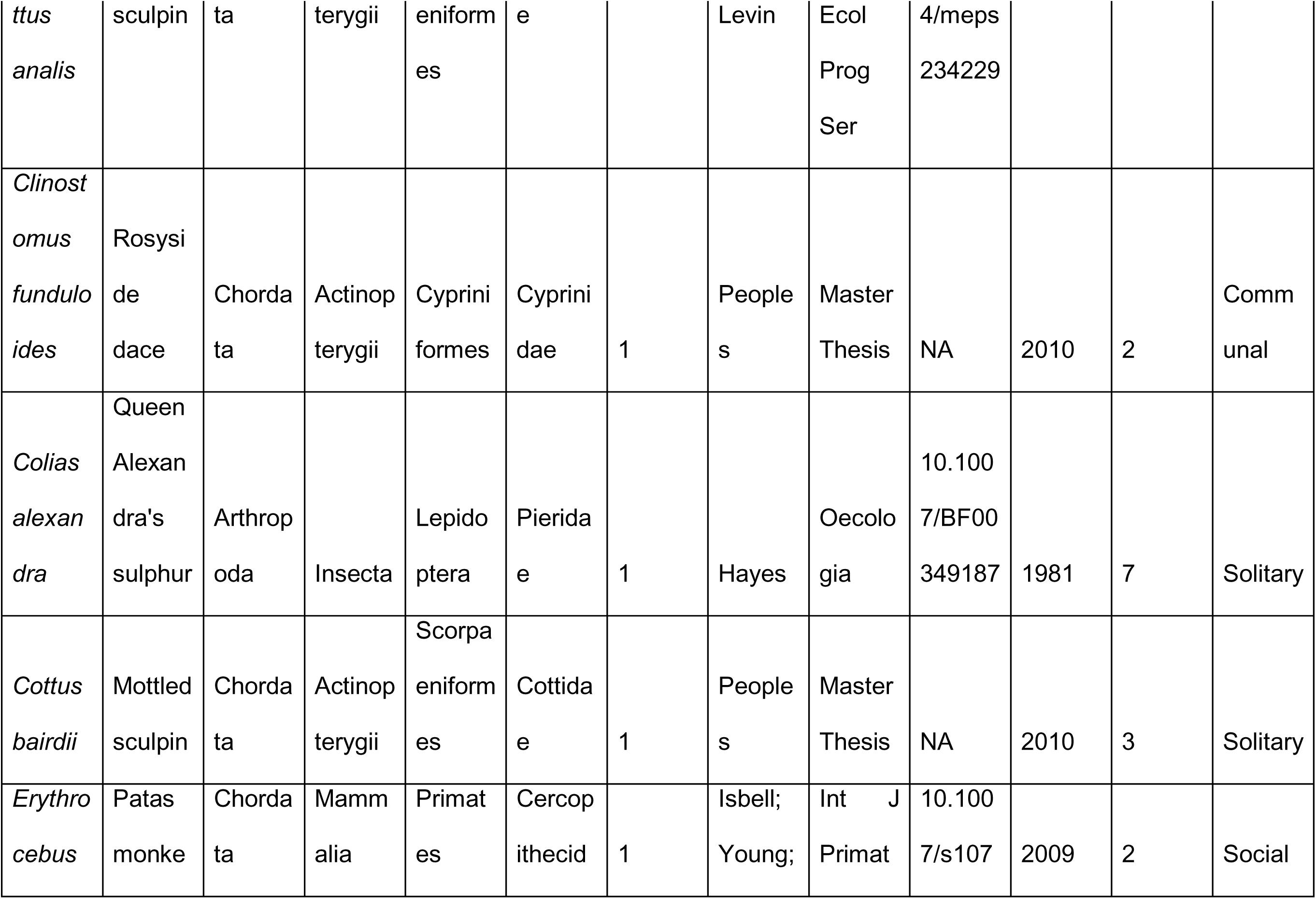
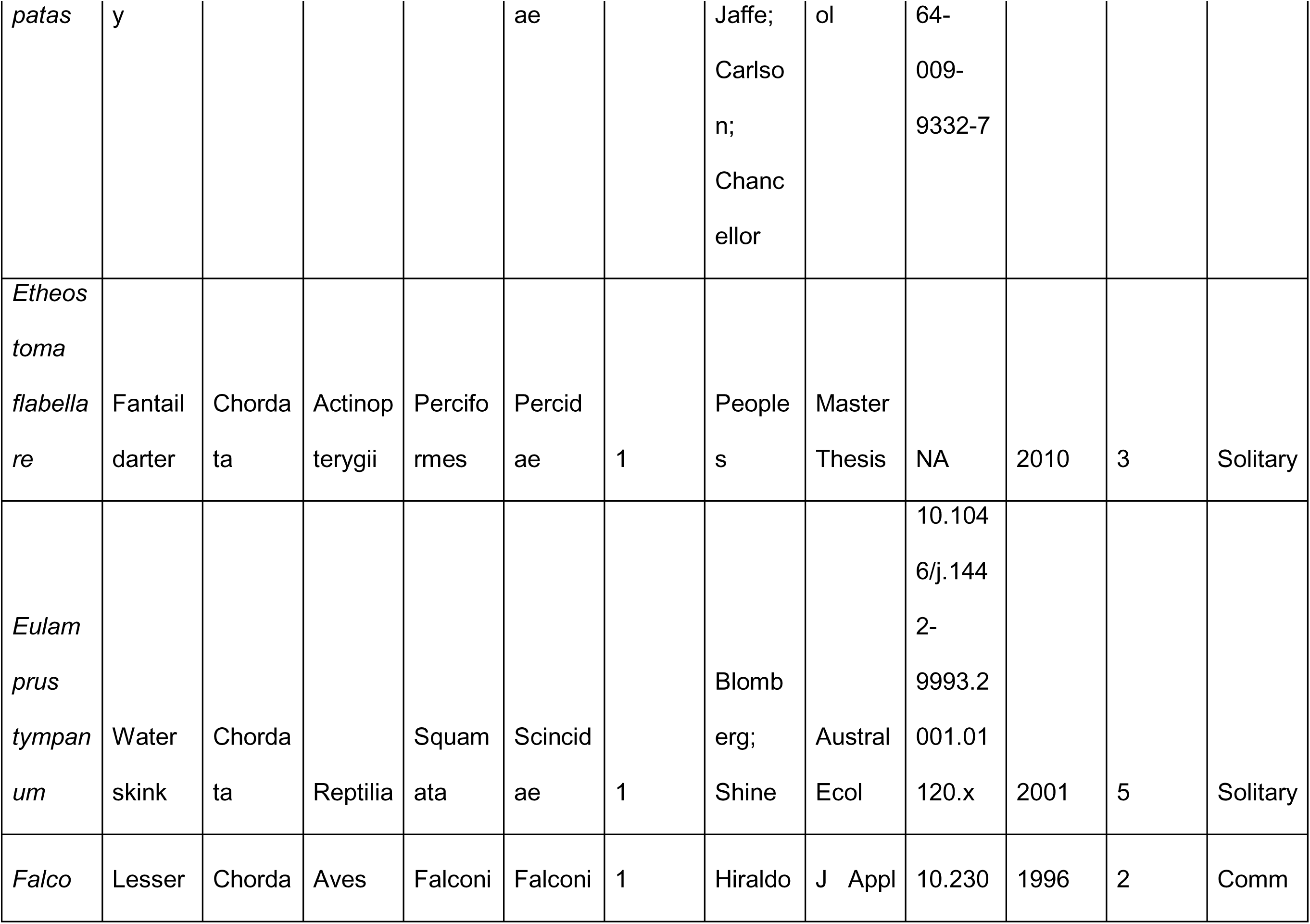
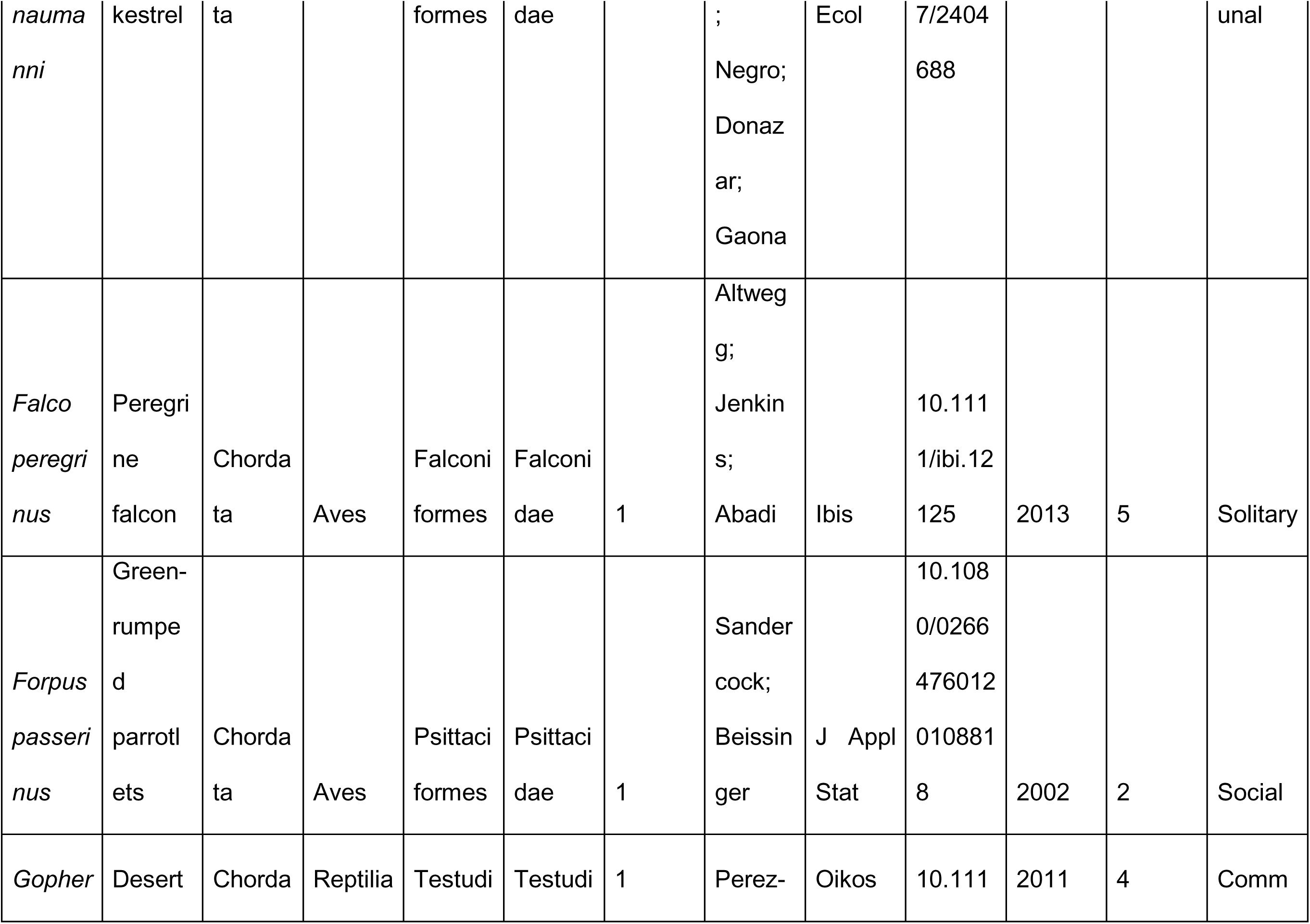
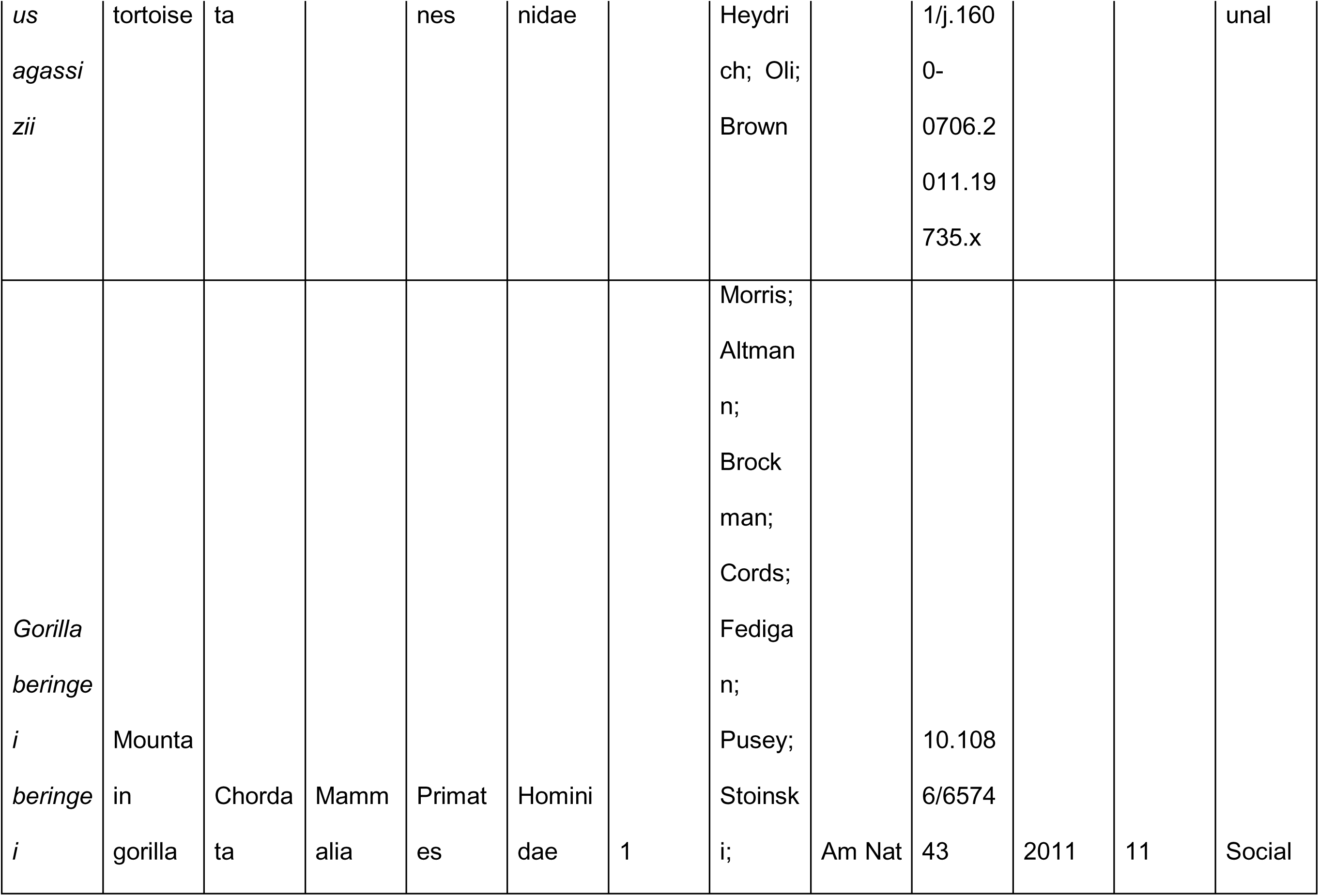
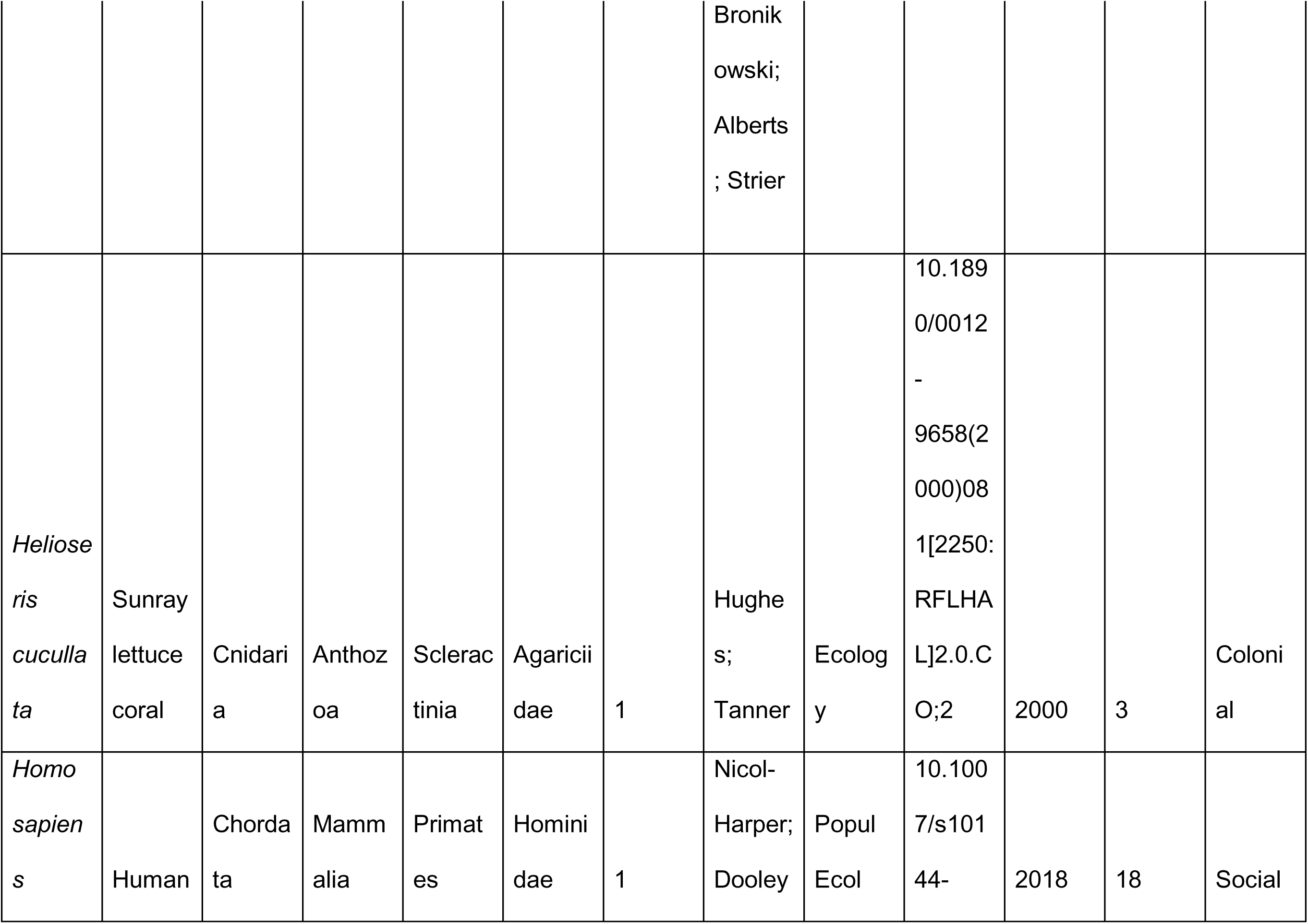
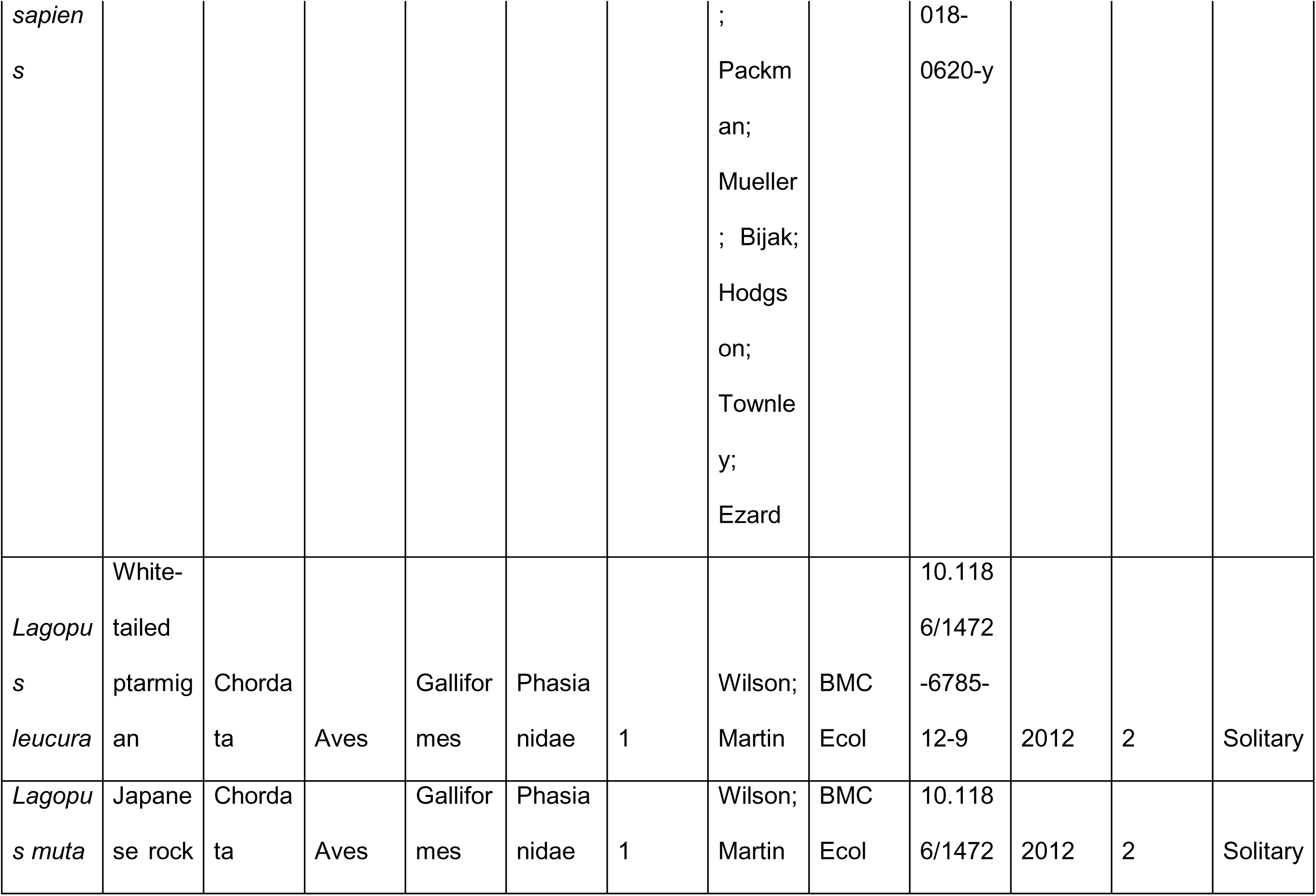
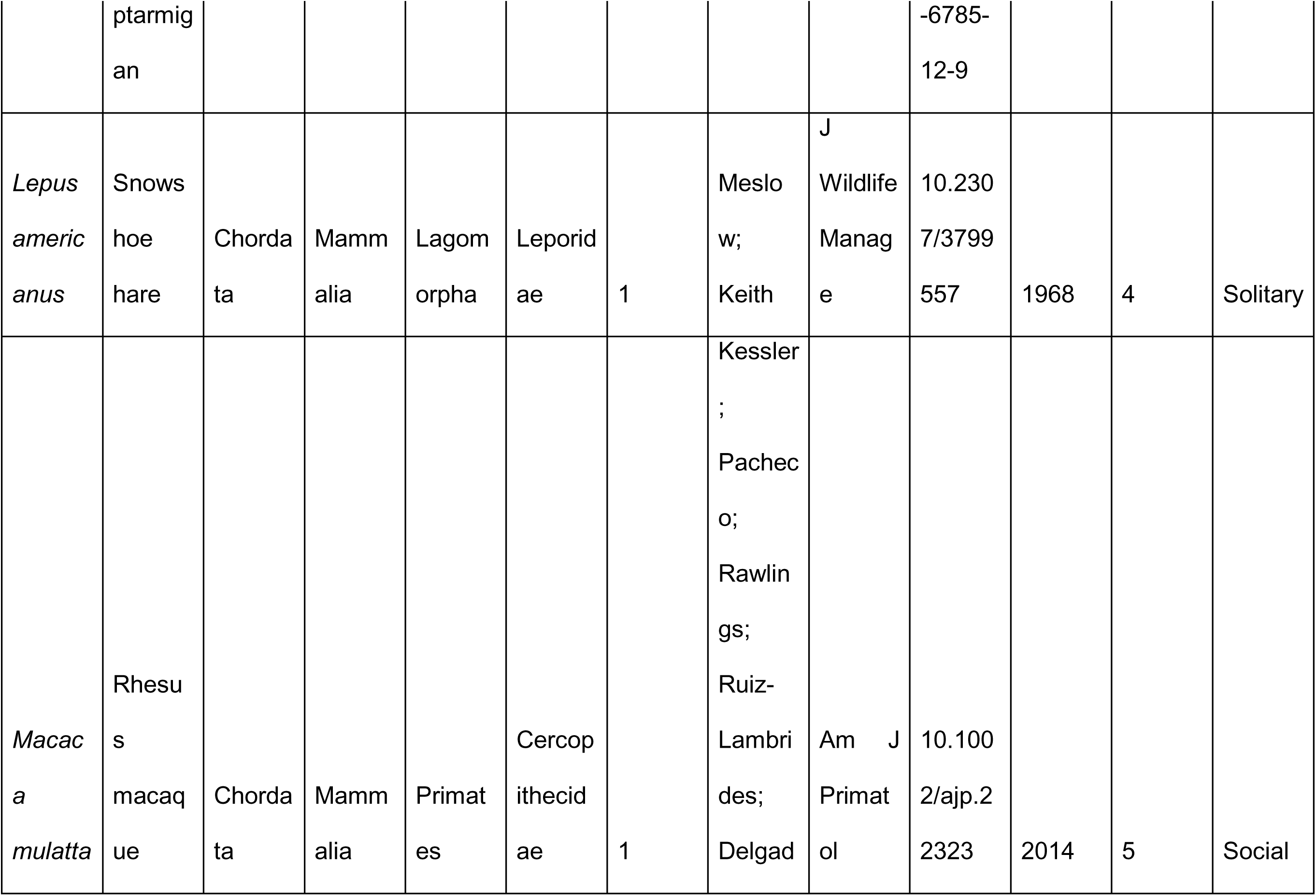
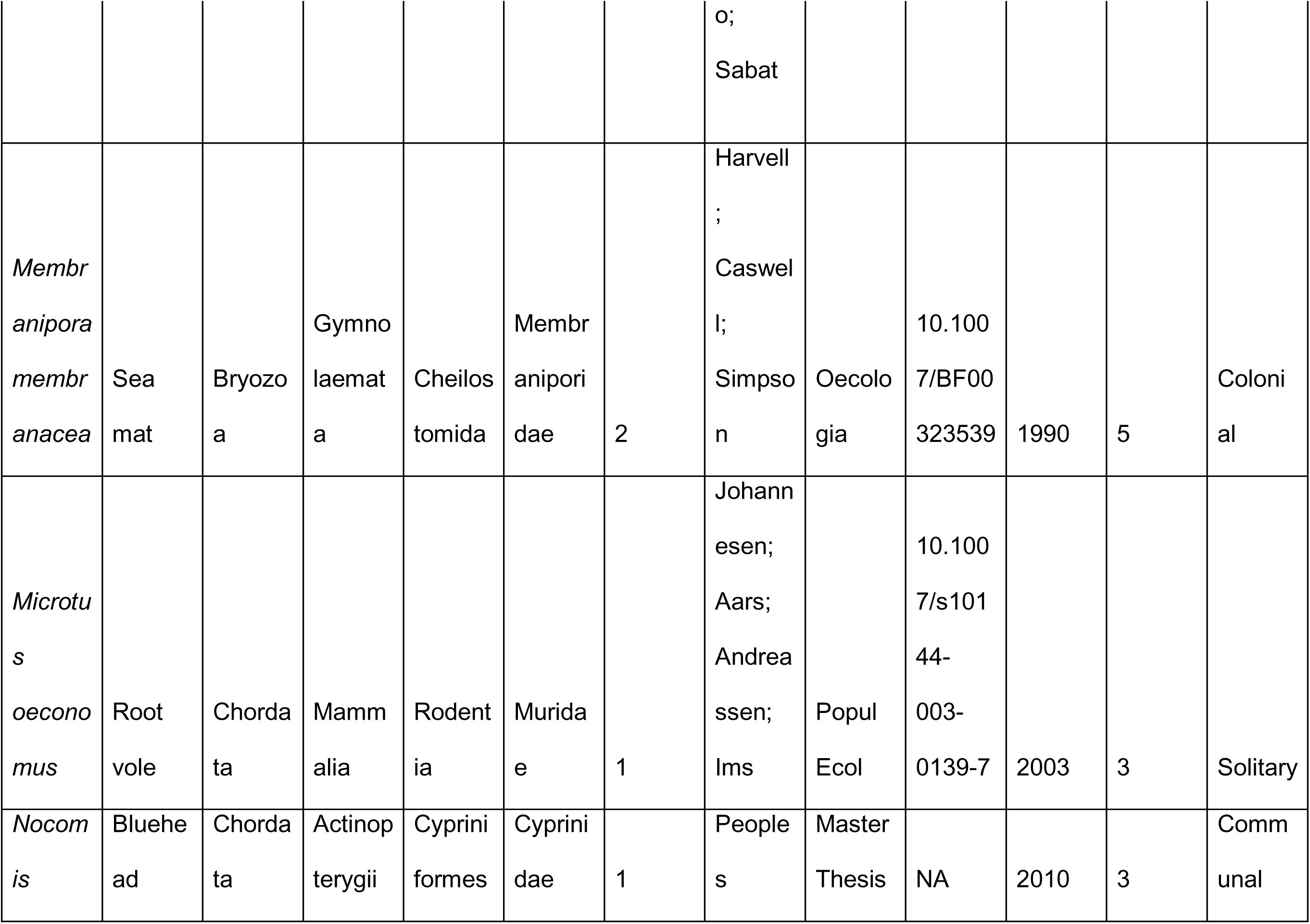
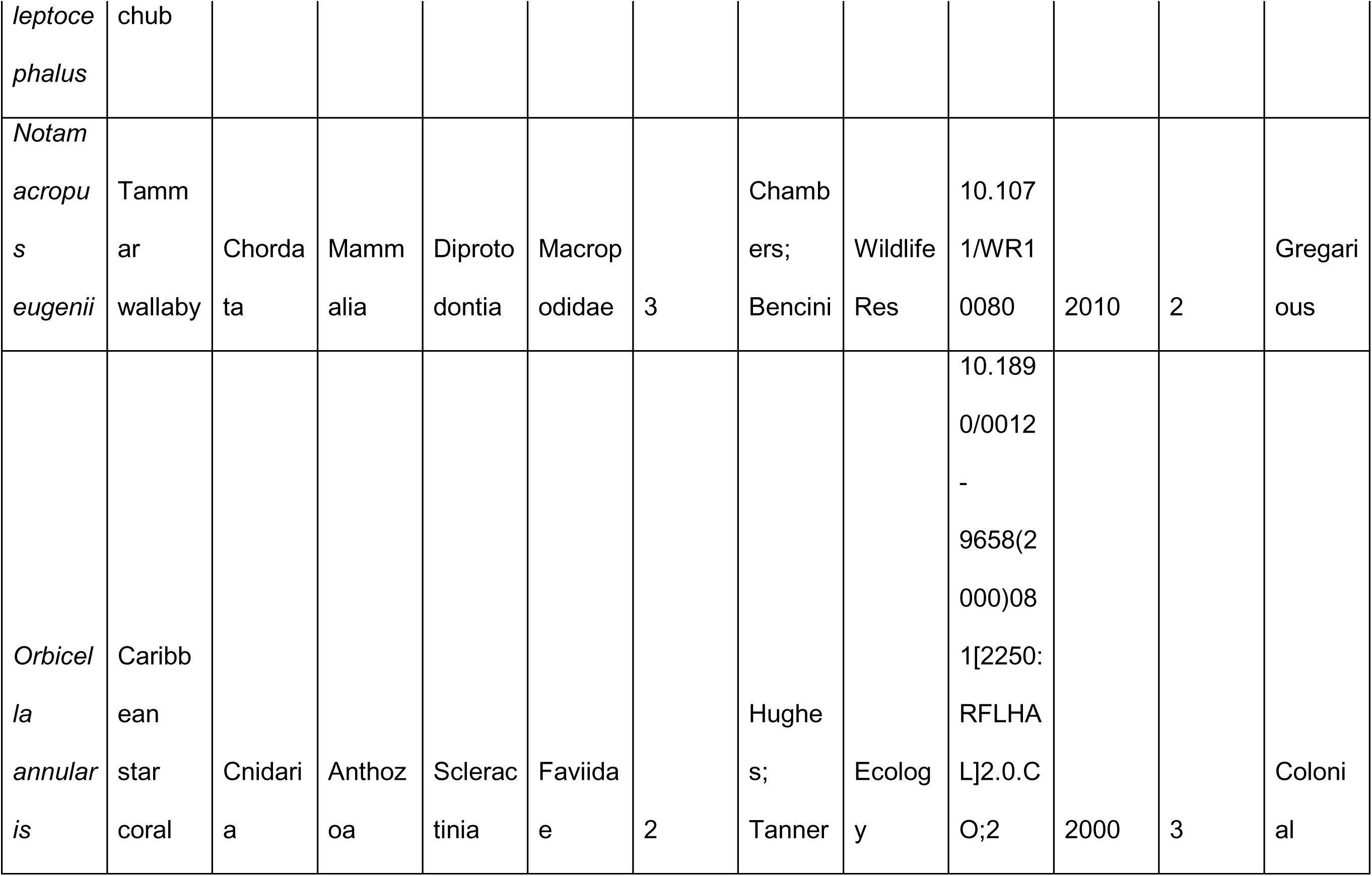
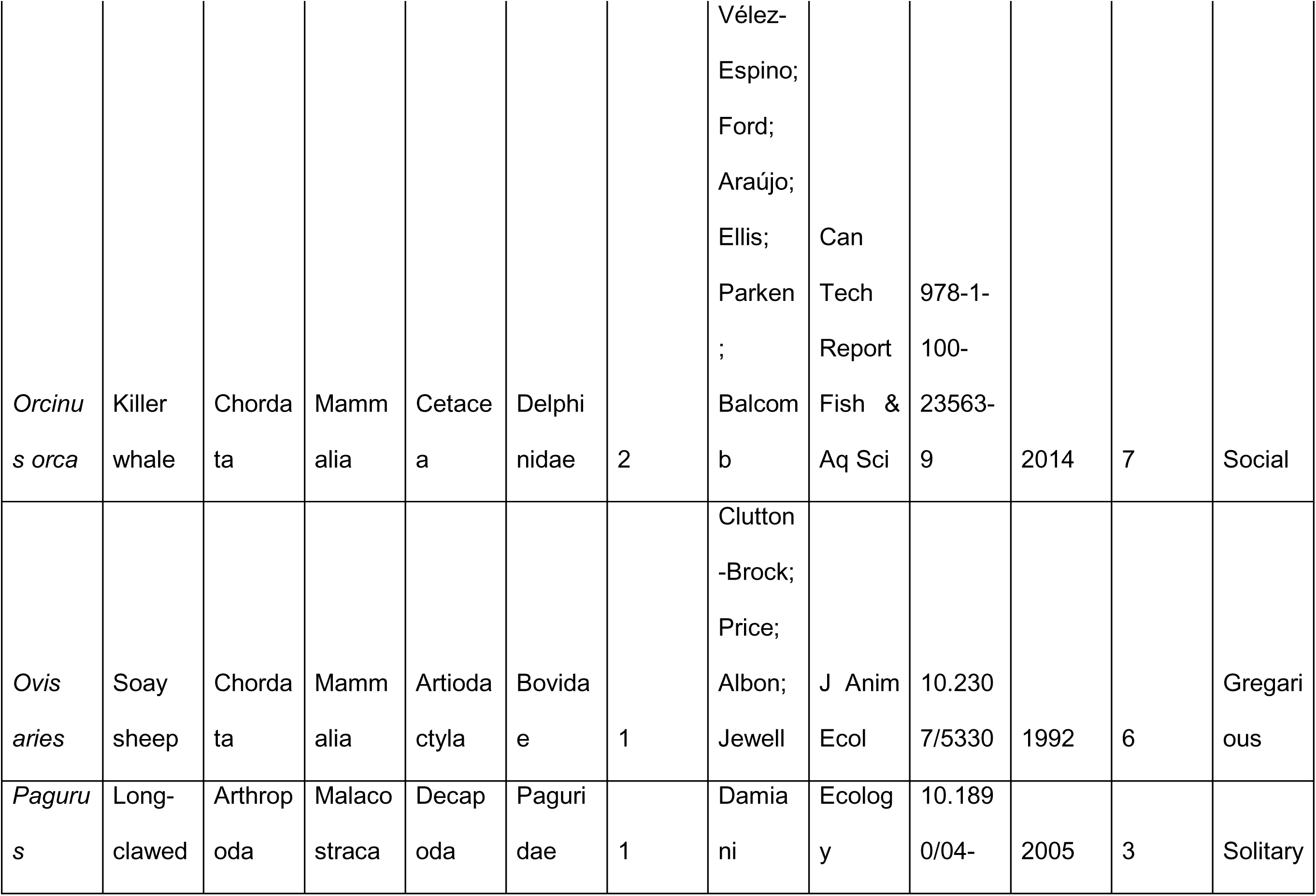
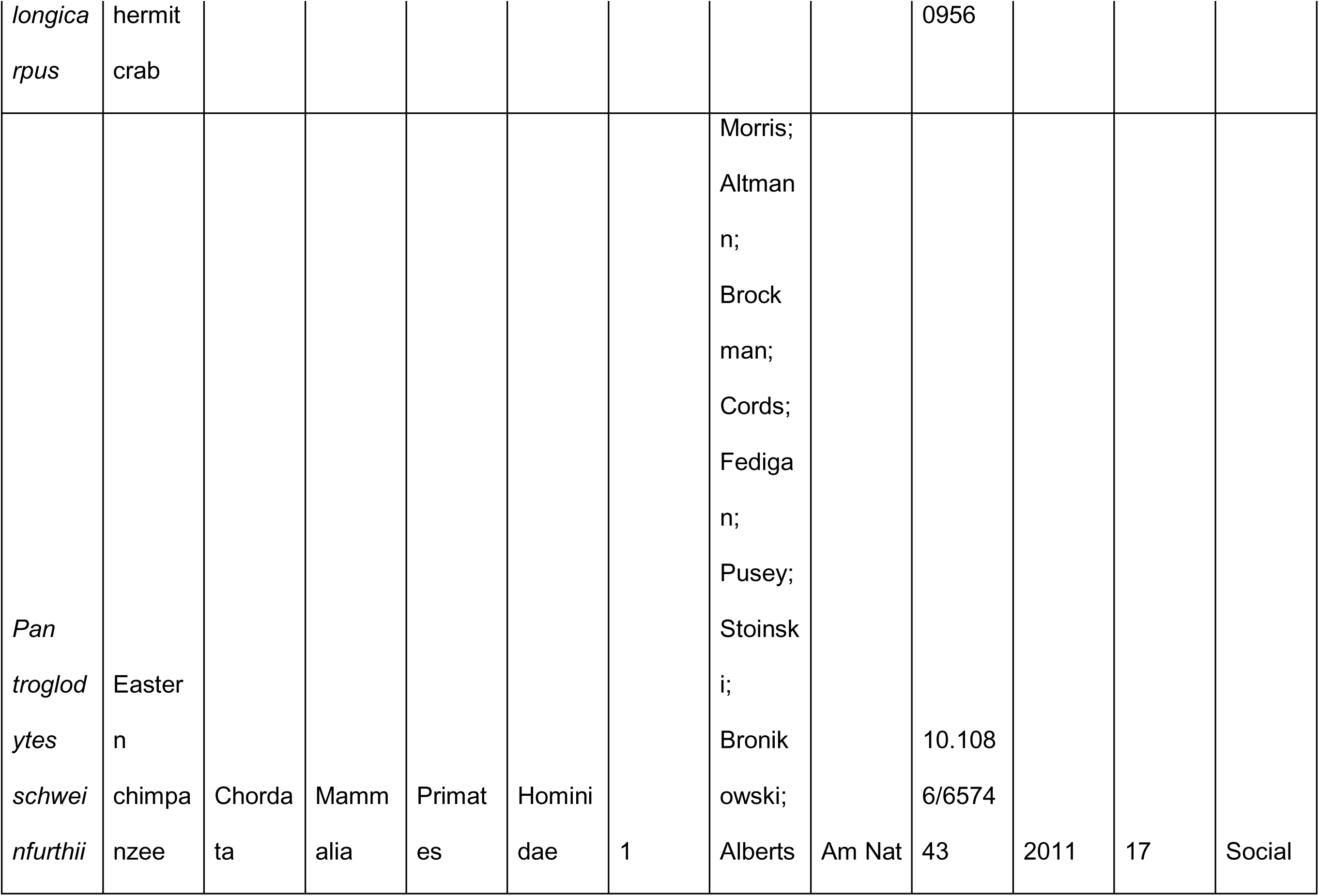
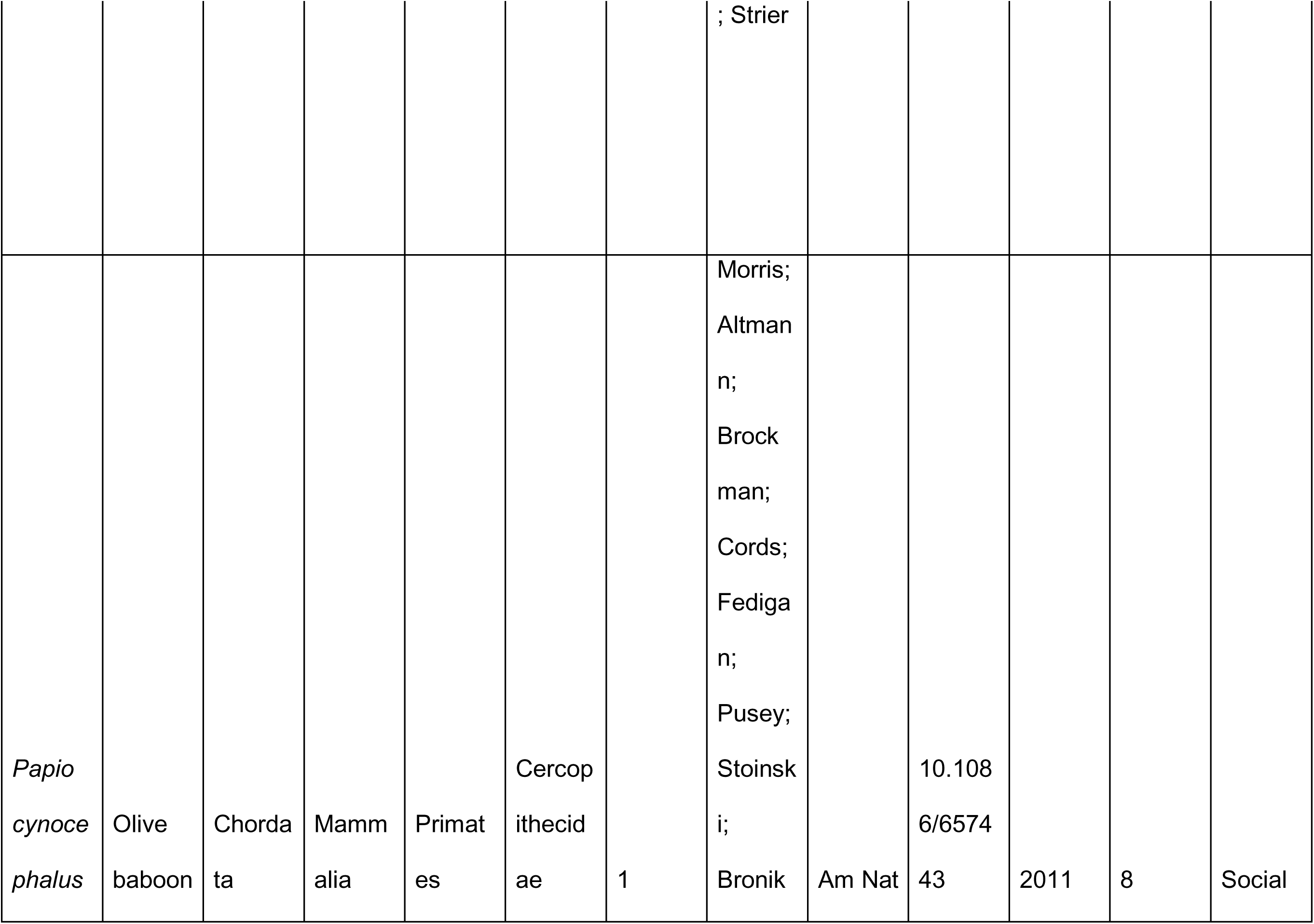
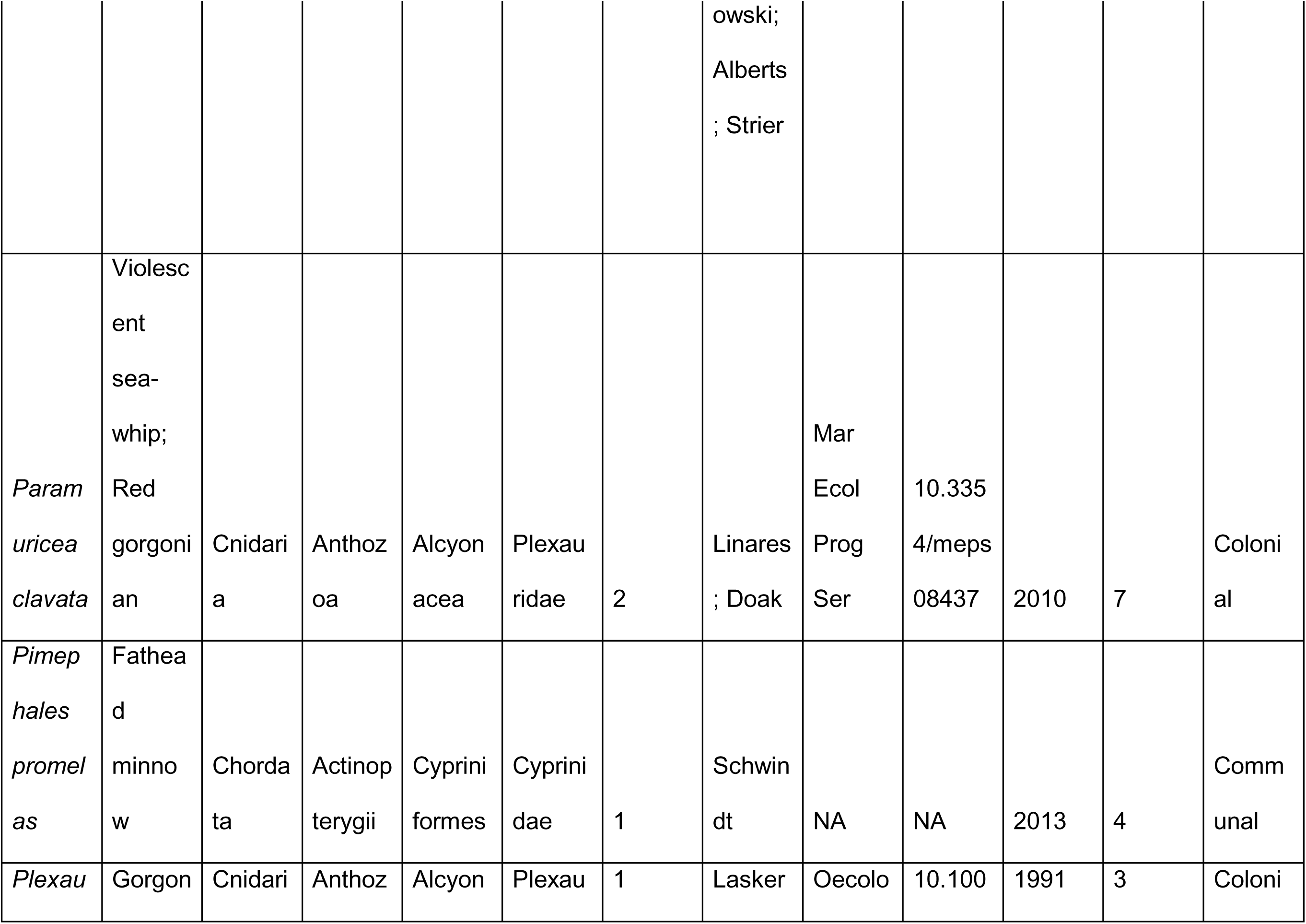
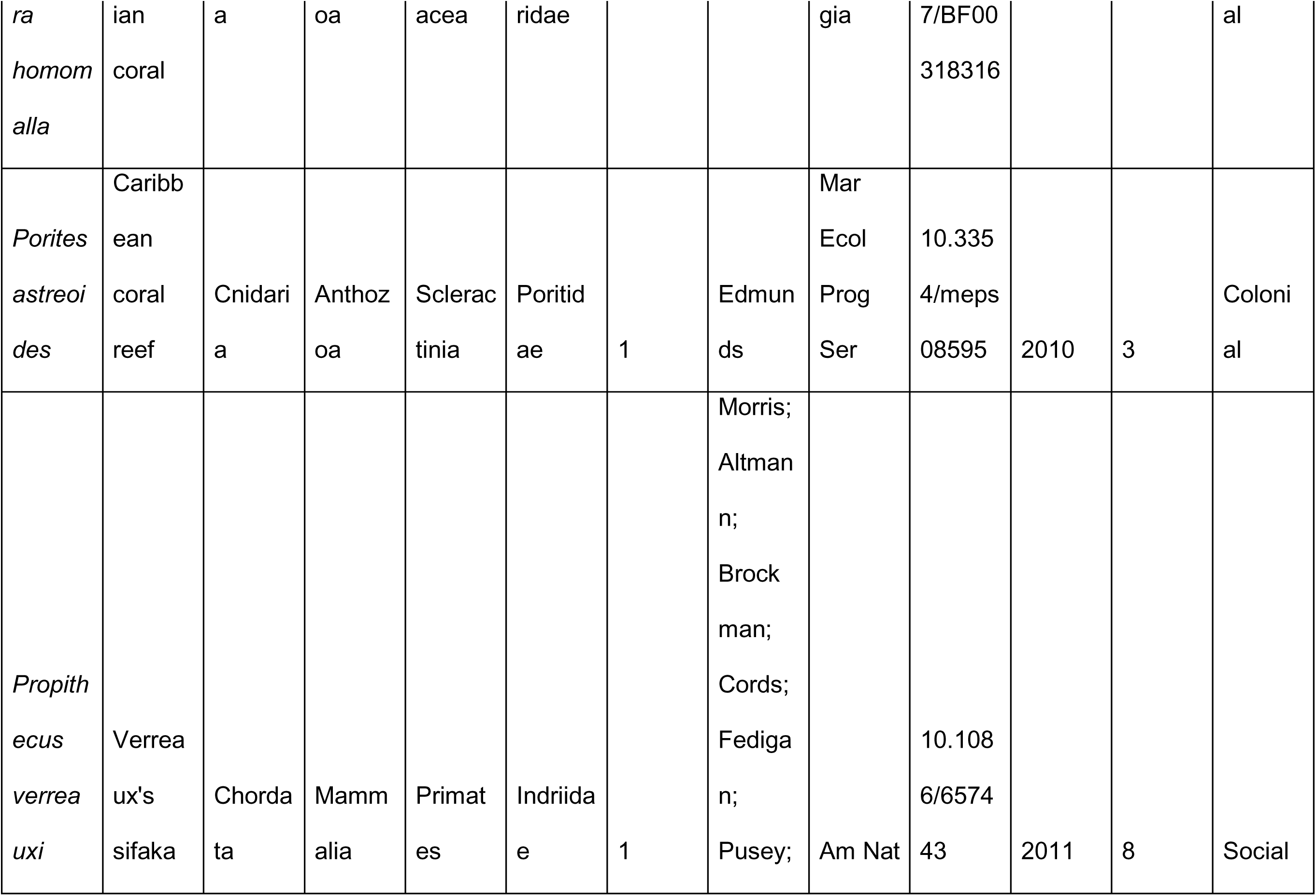
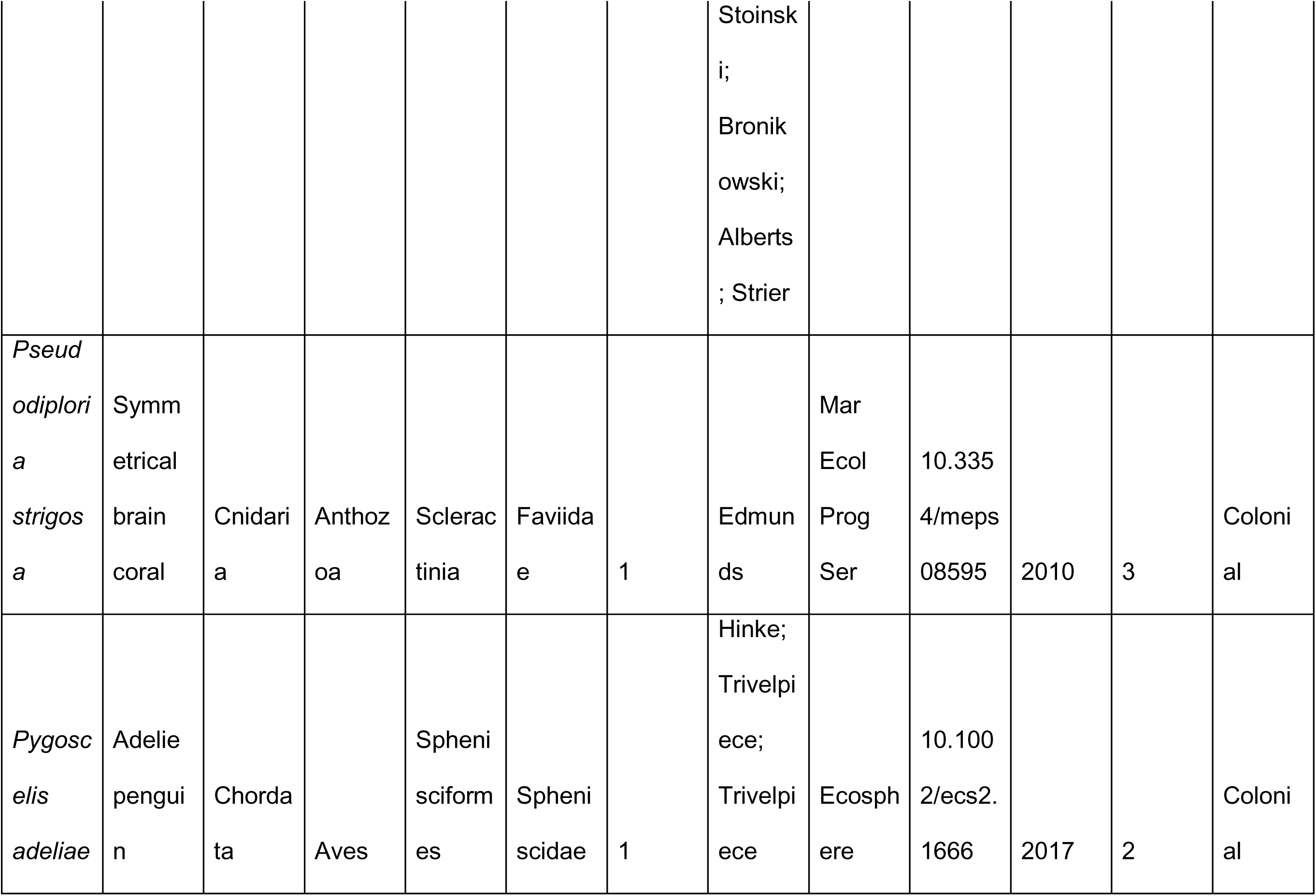
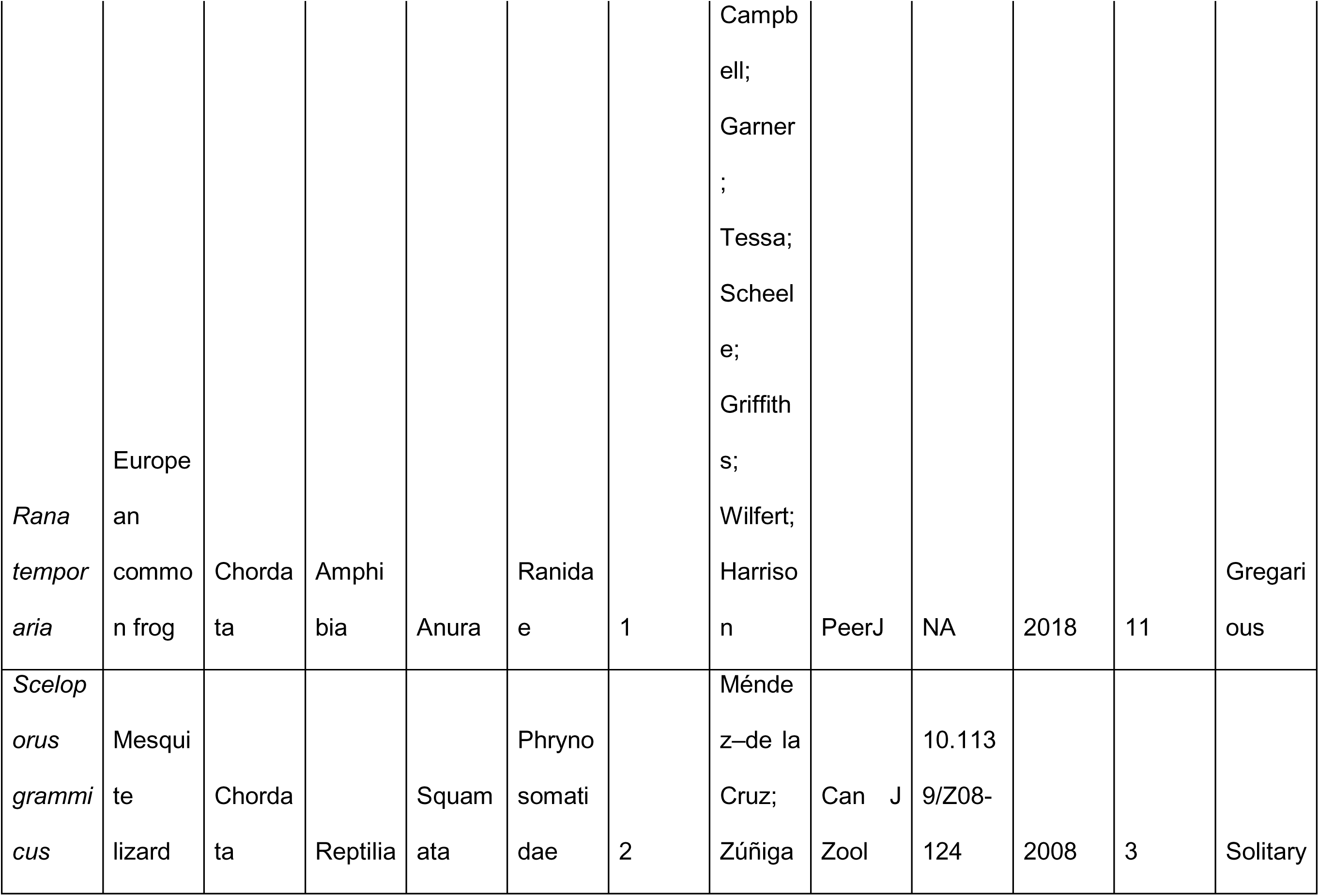
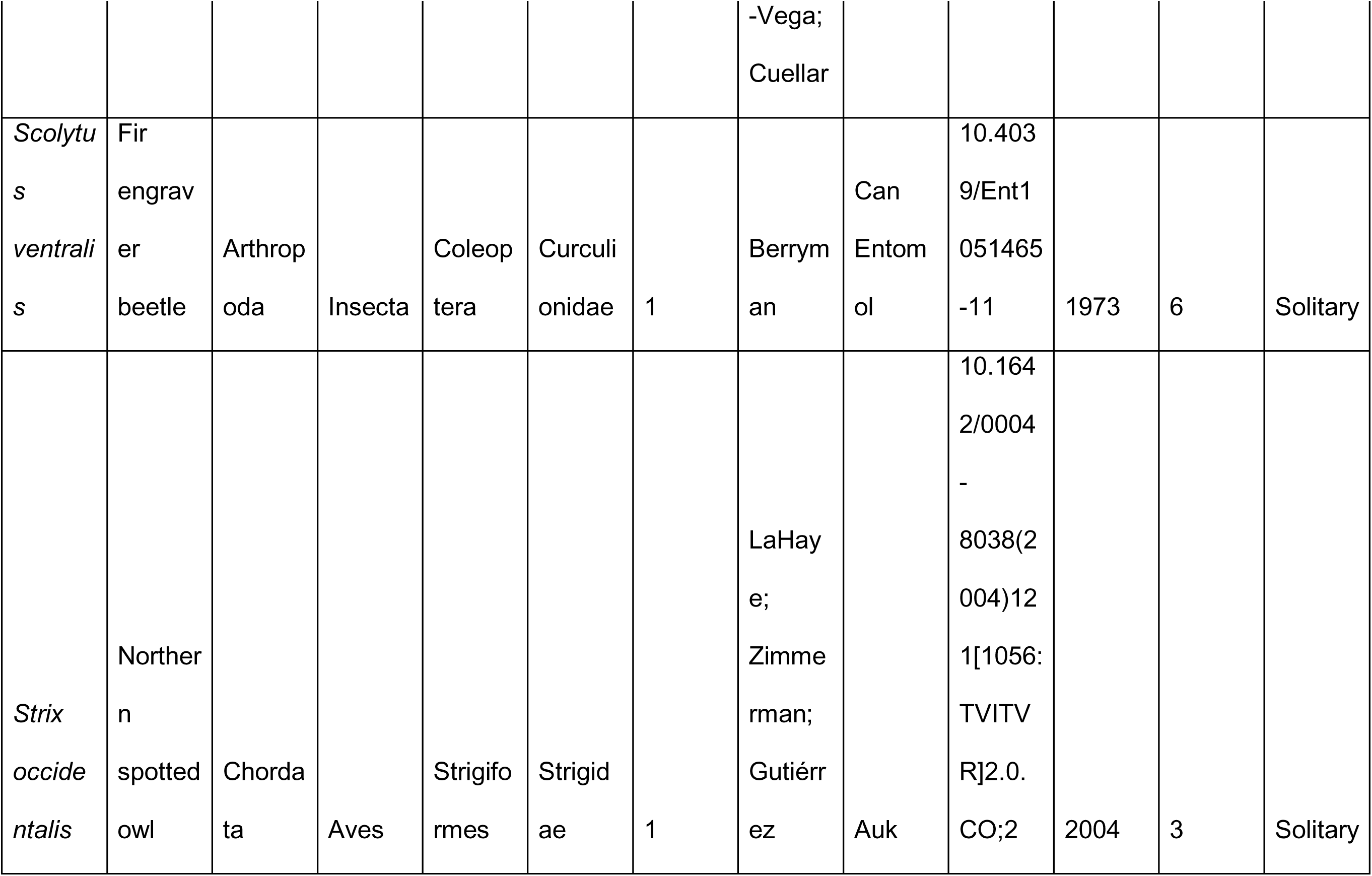
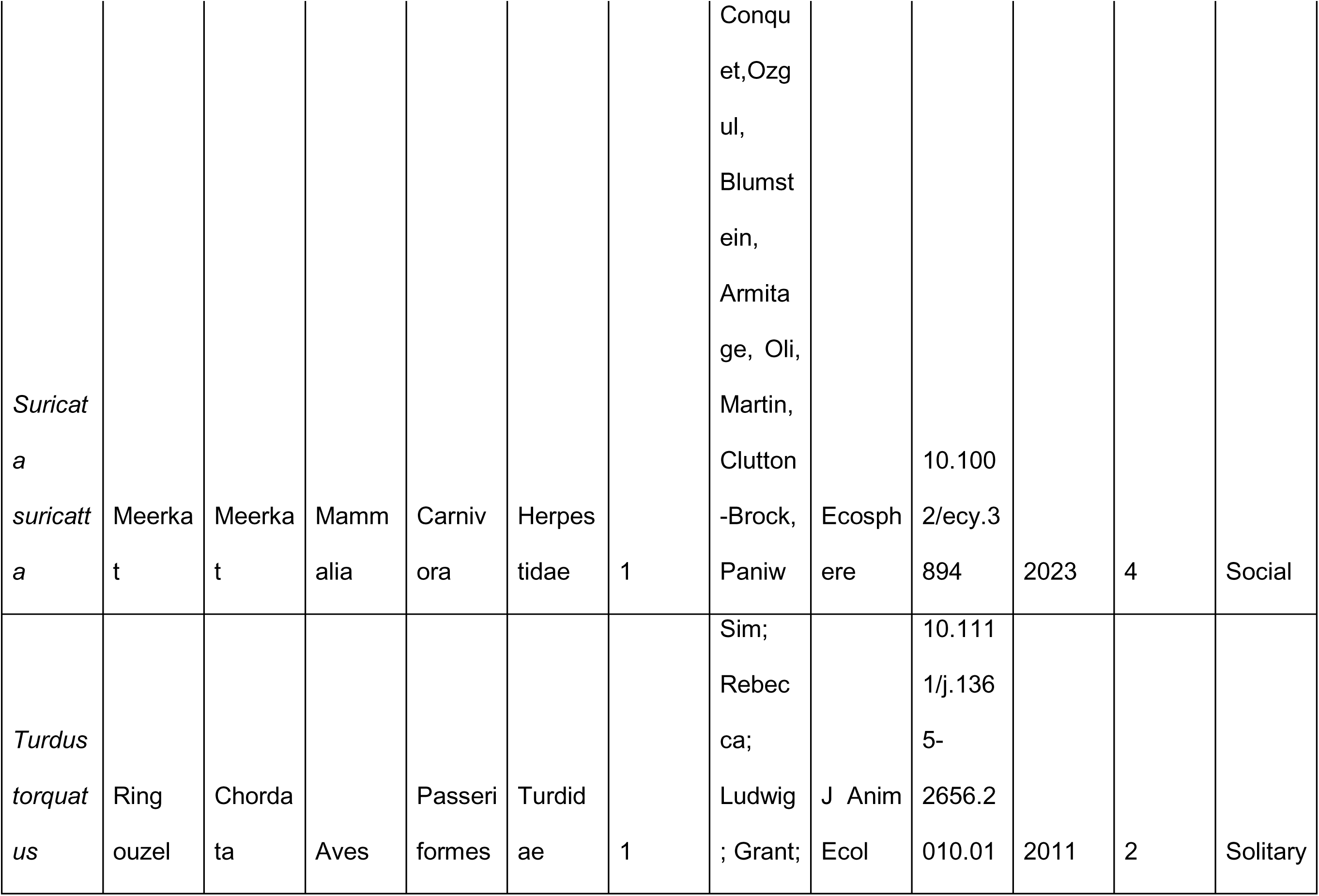
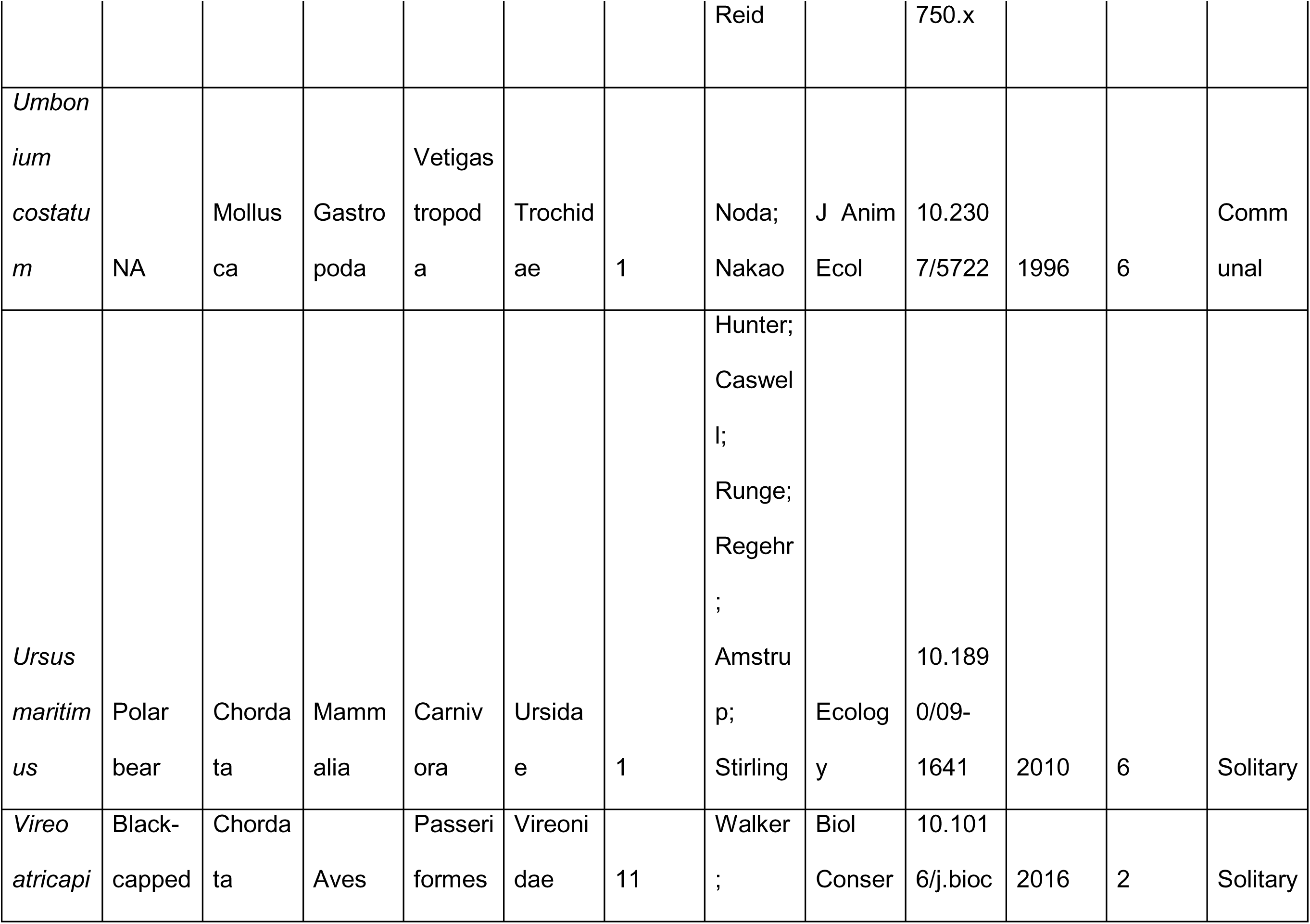
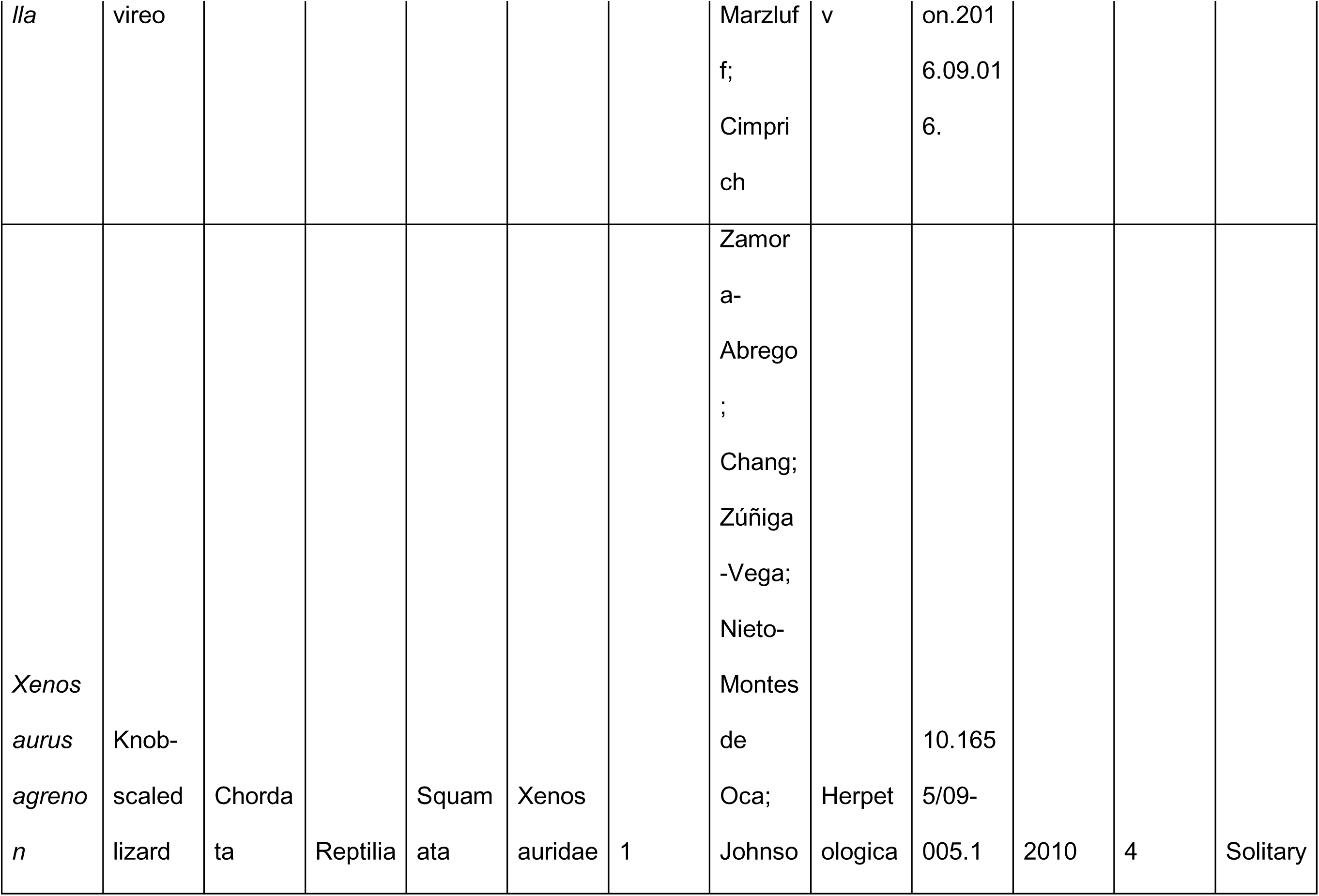
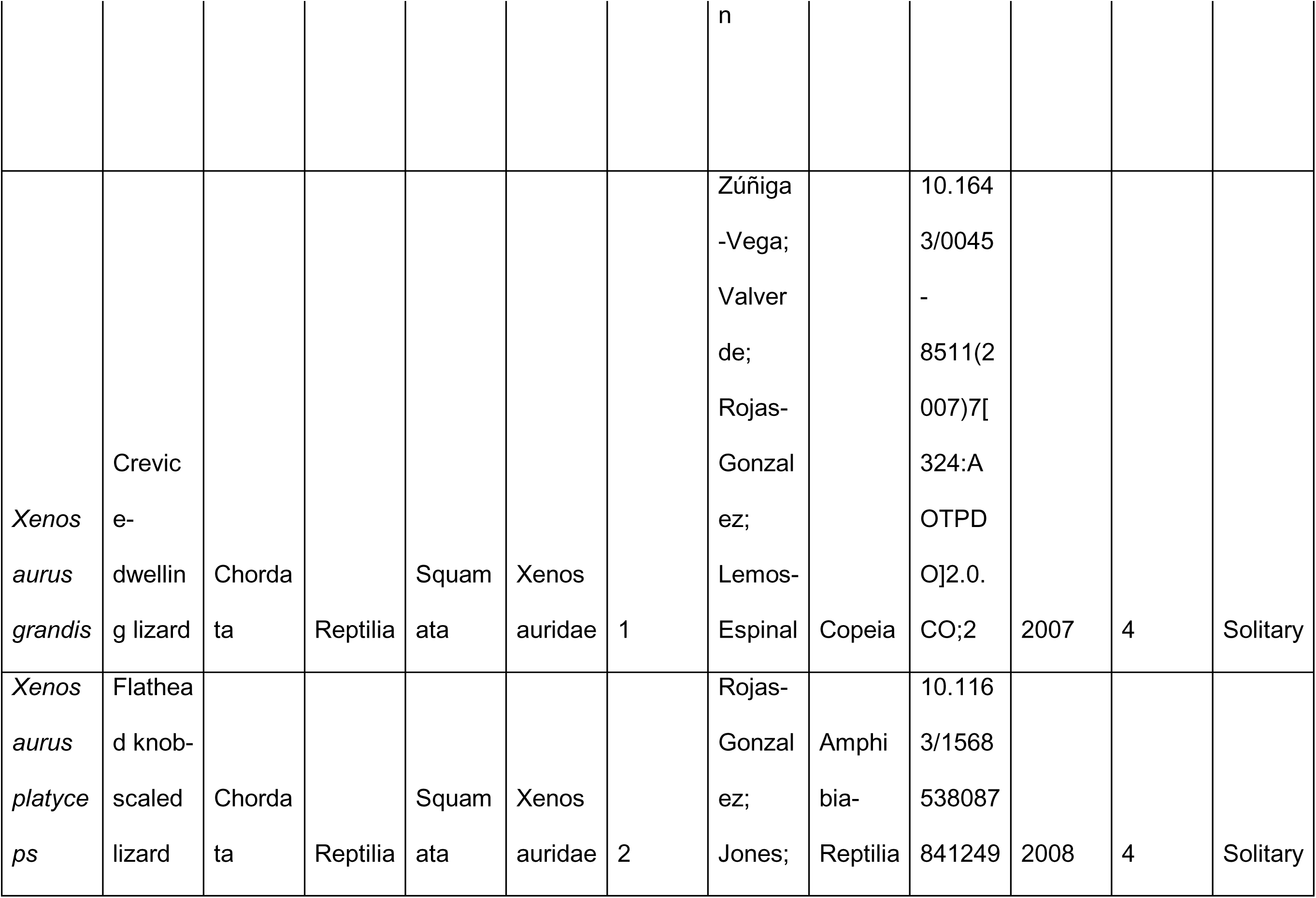
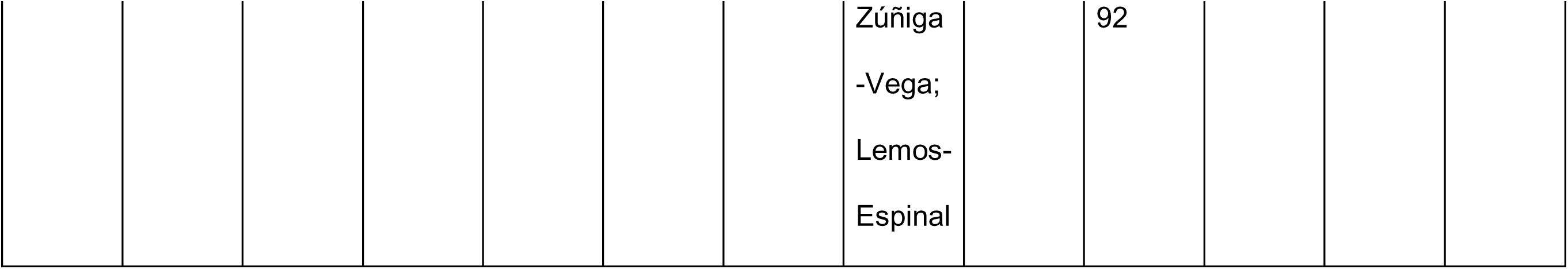
Metadata and source of matrix population models Sources of the demographic data obtained from COMADRE for the 66 examined species, together with other pertinent details. *Population* details the number of populations available per species in this study. *Dimension* details the number of stages the original matrix population models (MPMs) had before being collapsed to a set of 2×2 MPMs (see Methods). *Sociality* details the assigned level of our sociality continuum, with the explanation for each of the five levels provided in the Methods too.

**Table S3.**
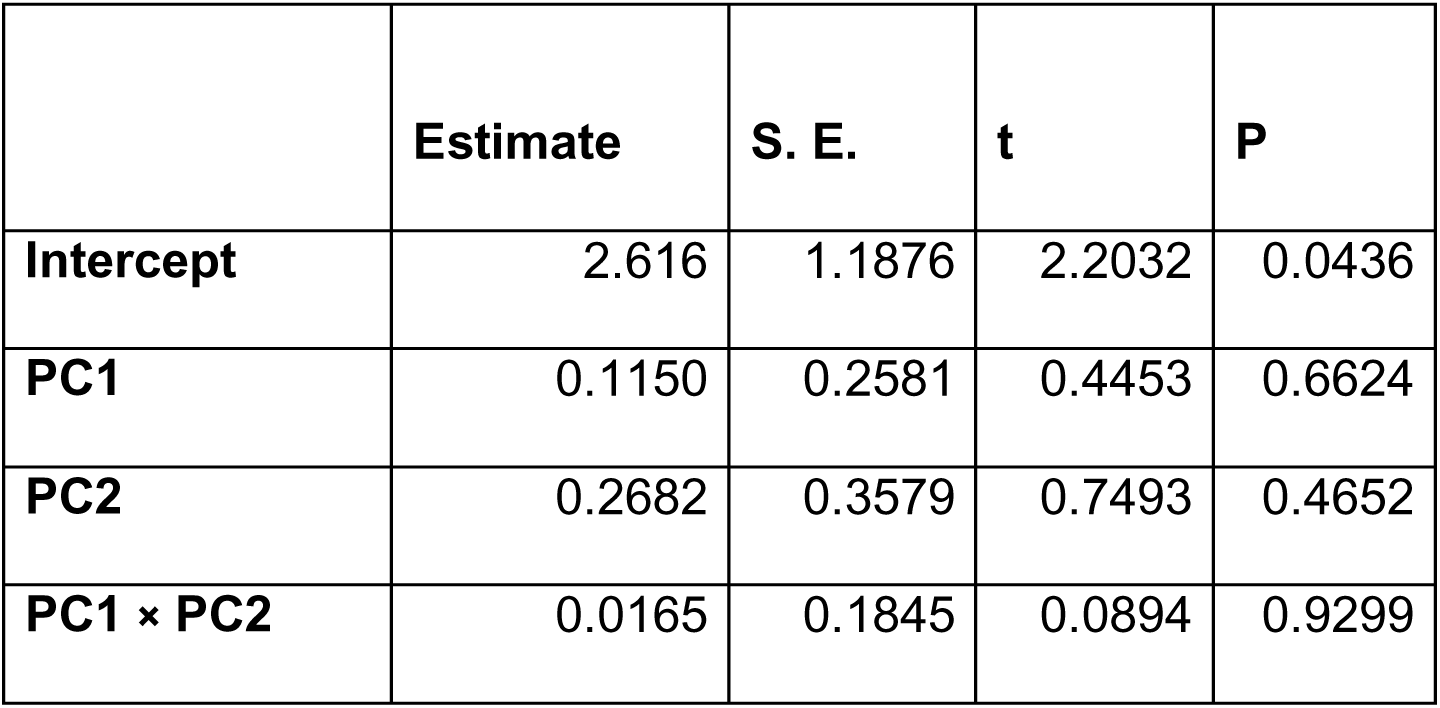
Degree of sociality across the climatic space The degree of sociality is not predicted by the climatic principal component axis. Results of a pgls model predicting the degree of sociality as a function of the positioning of each population along PC1 (Precipitation predictability; Figure 2) and PC2 (temperature constancy).

**Figure S1.**
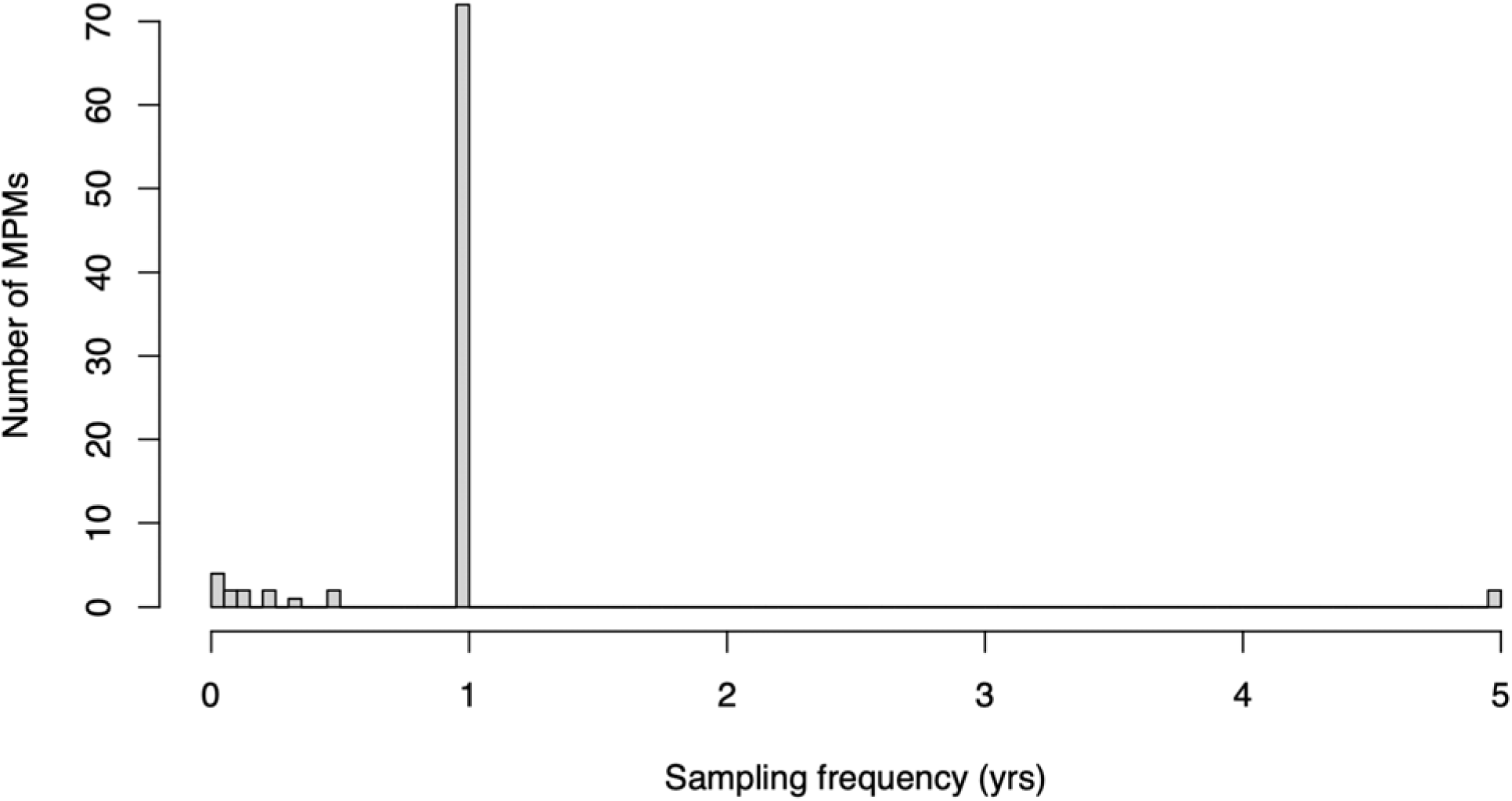
Sampling frequency of the matrix population models Most of the matrix population models (MPMs) in our selected set of matrix population models from the COMADRE Animal Matrix Database were sampled once a year. Species in our study with an original sampling frequency that is not annual: *Caenorhabditis elegans* (0.000114), *Capitella capitata* (0.0192), *Xenosaurus agrenon* (0.08), *Membranopira mebranaceae* (0.12), *Acyrthosiphon pisum* (0.02), *Clinocottus analis* (0.25), *Ampelisca abdita* (0.03), *Eulamprus tympanum* (0.08), *Sceloporus grammicus* (0.5), *Montastrae annularis* (5), and Homo sapiens sapiens (5).

**Figure S2.**
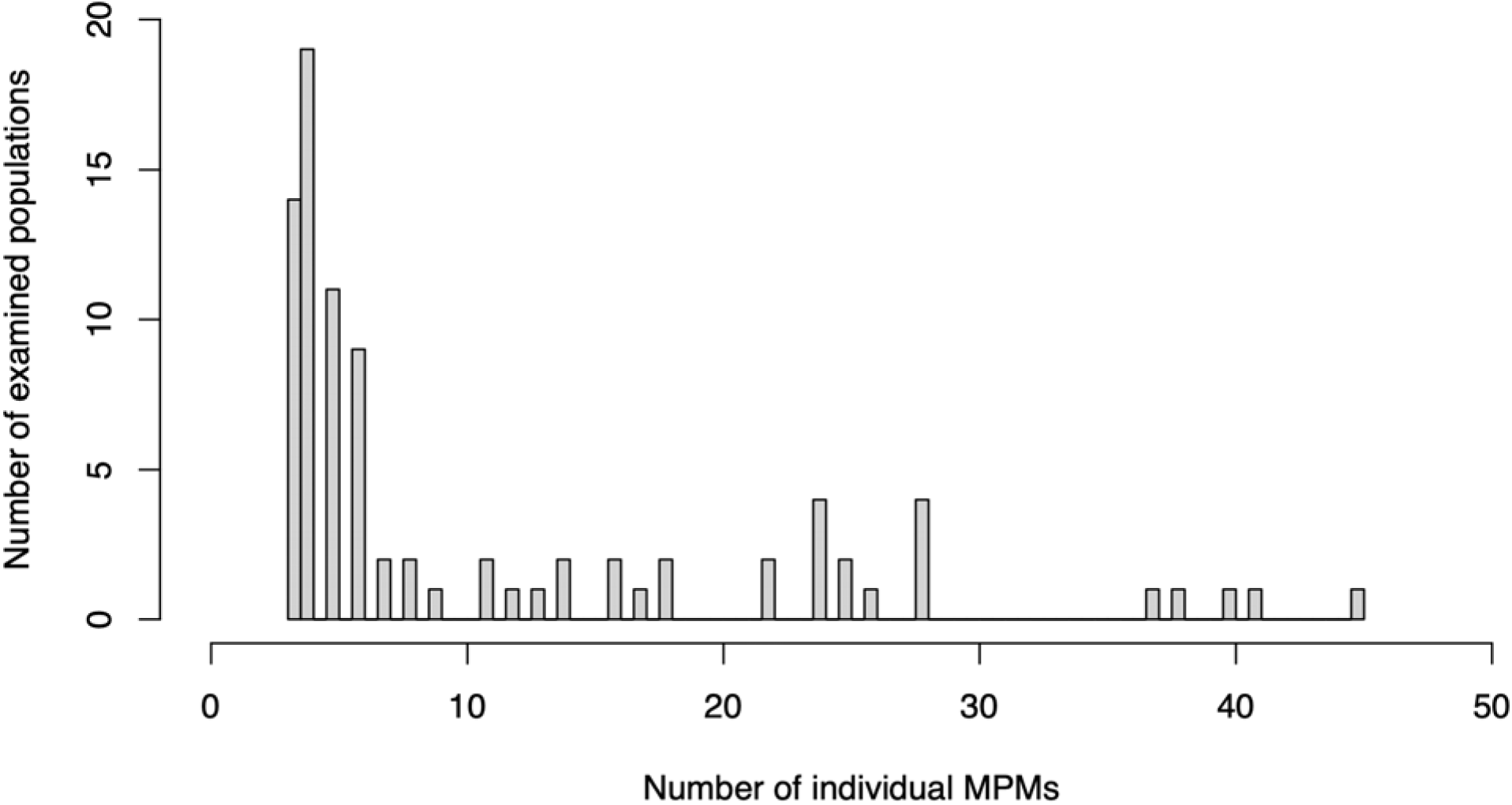
Matrix population model availability per study Number of matrix population models (MPMs) per studied population in our selected subset of 66 species from the COMADRE Animal Matrix. Total MPM = 955. Total number of separate populations = 87.

**Figure S3.**
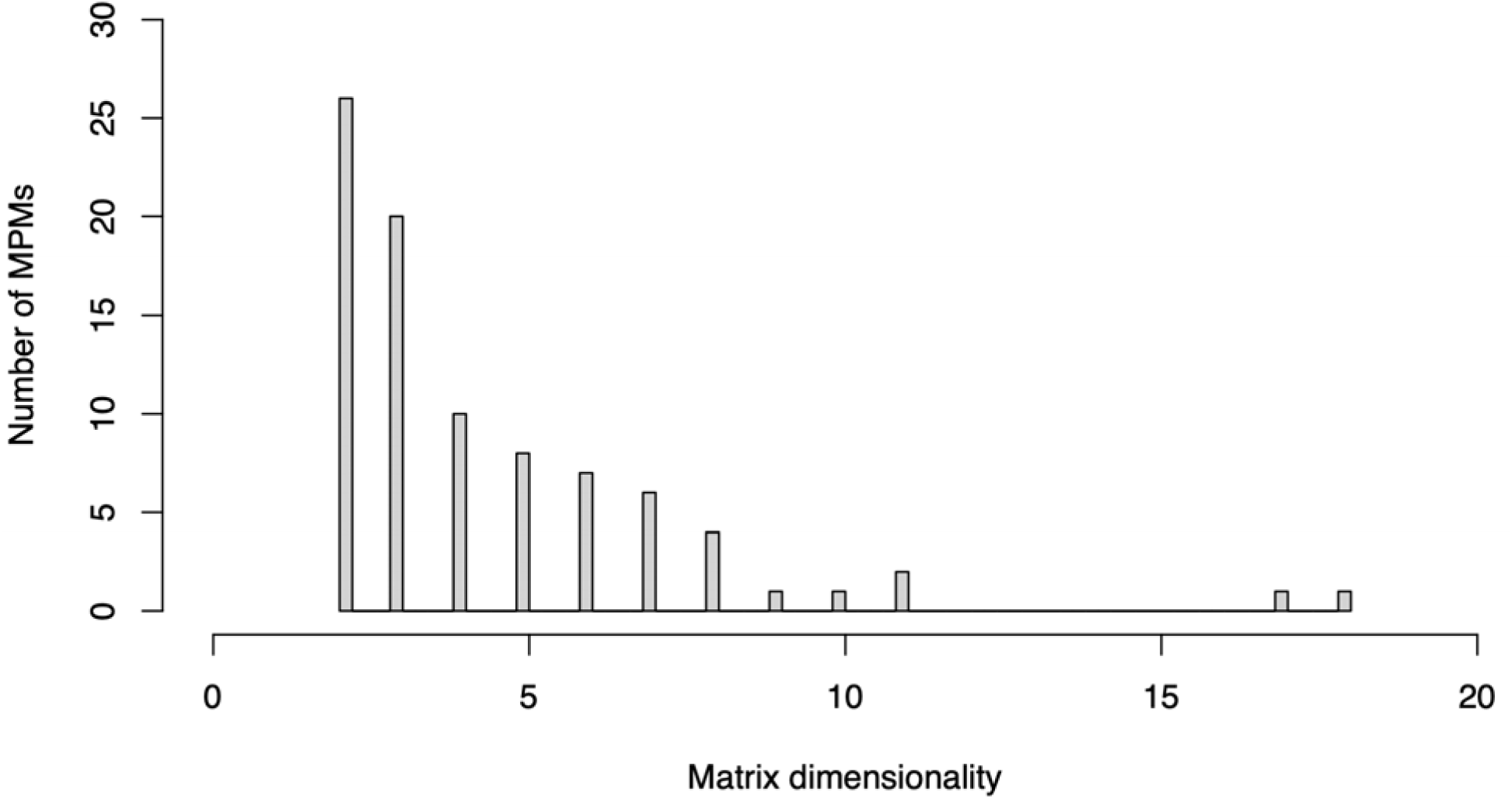
Matrix population model dimensionality The dimensionality of our examined matrix population models (MPMs) varied considerably between 2 and 18 stages. As such, we collapsed all models to a two-stage MPM to allow for stage comparability in vital rates.

**Figure S4.**
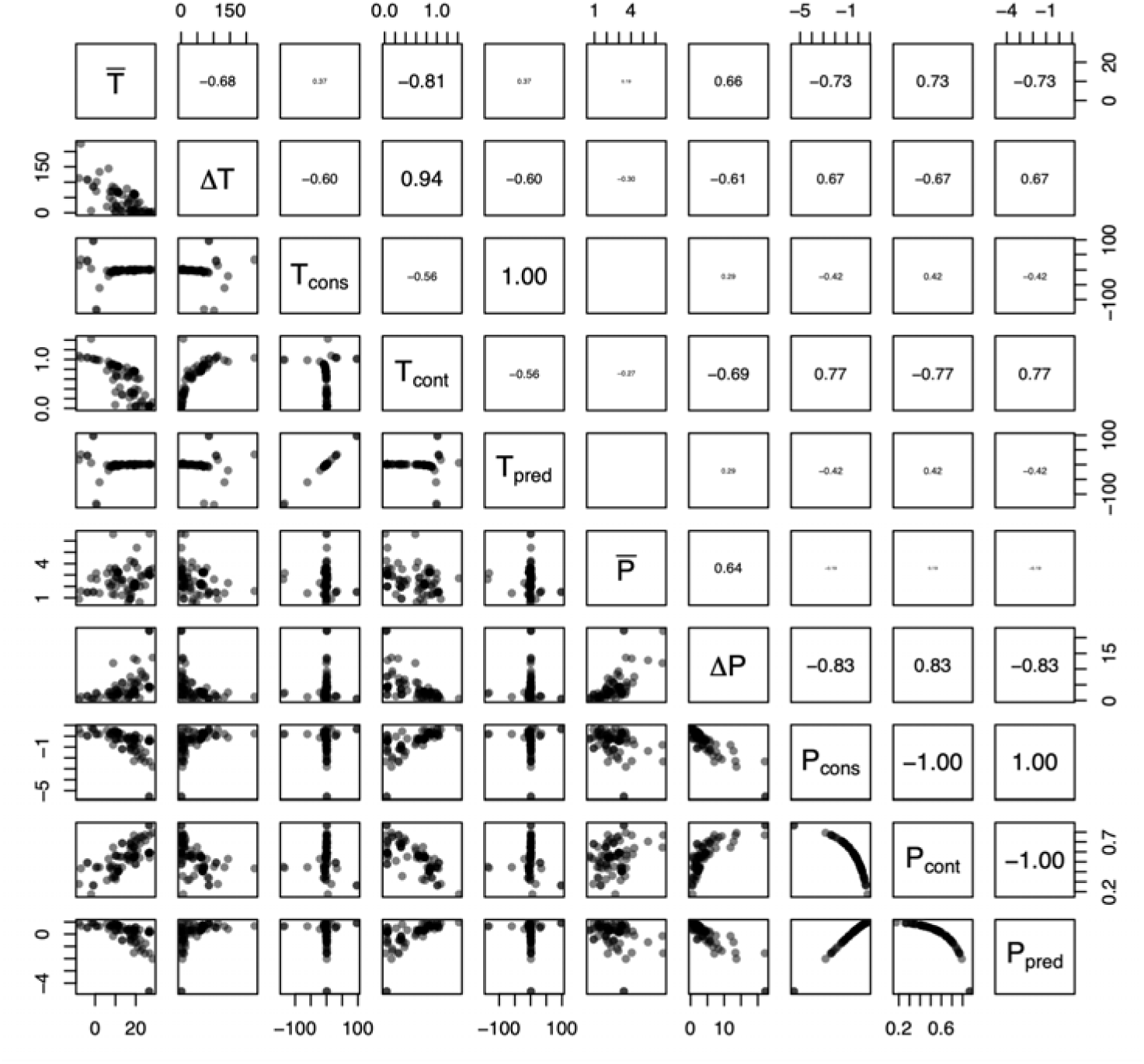
Climate driver collinearities Some of the examined climatic drivers are strongly collinear. Pairwise correlations (top-right) and scatterplots (bottom-left) of the statistics of monthly precipitation and temperature derived from the NASA POWER database to quantify the degree of environmental stochasticity to which our 87 natural populations have been exposed. Temperature statistics include: : mean; Δ*T*: variance: *T_cons_*: constancy; *T_cont_*: contingency; and *T_pred_*: predictability. Ditto for precipitation (*P*). Constancy quantifies the extent to which a climatic variable remains stable over time (1 - variance/mean); contingency quantifies the extent to which fluctuations are structured and recurrent; predictability = constancy + contingency, as per Colwell (Colwell, 1974). Font size of spearman correlation coefficient is proportional to its values.

**Figure S5.**
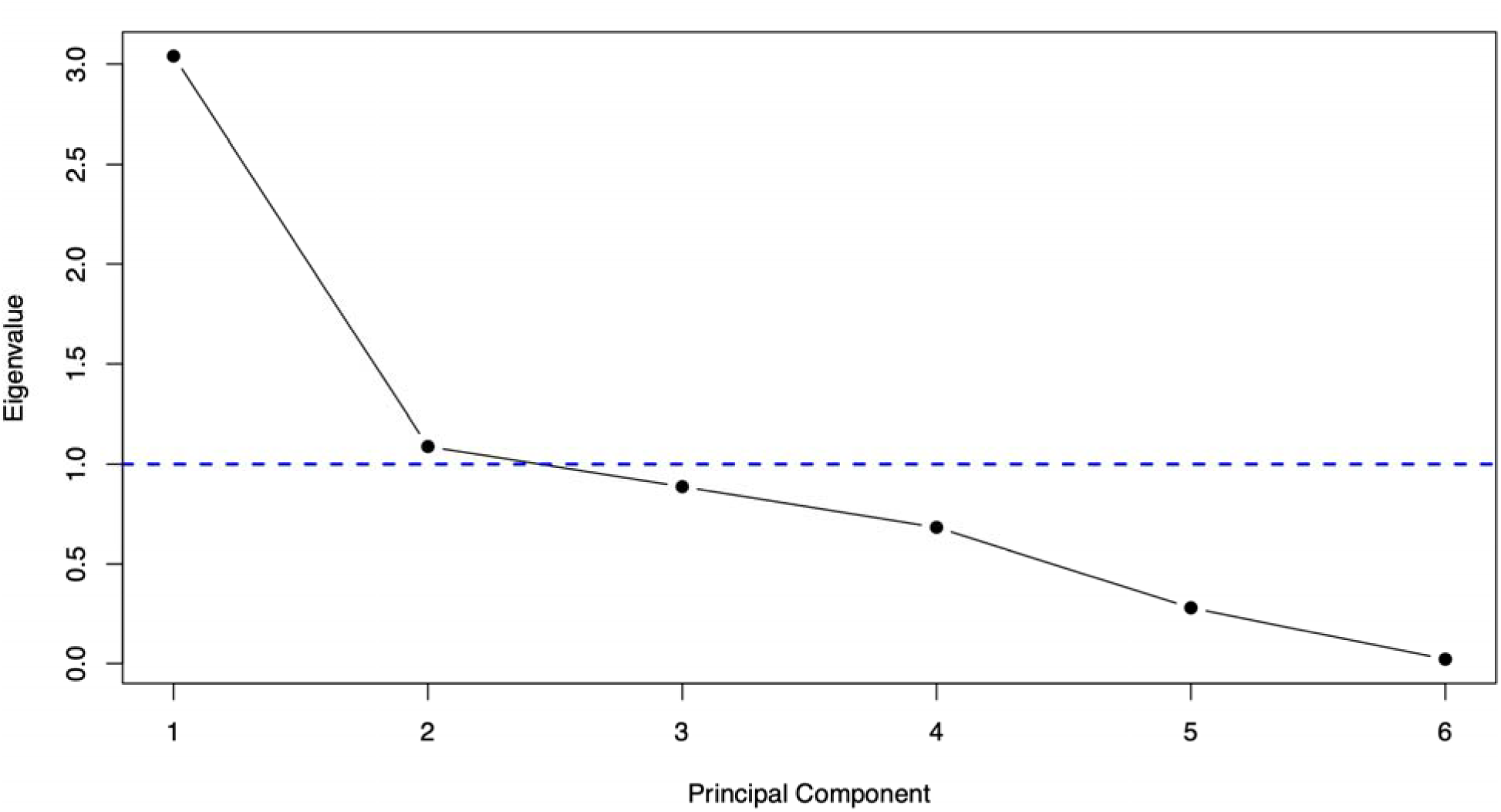
PCA screeplot Screeplot of the six principal components of the climatic PCA. The blue dashed line separates the principal components axis with associated eigenvalue > 1, which we retained for the next steps in our analyses, as per the Kaiser criterion.

**Figure S6.**
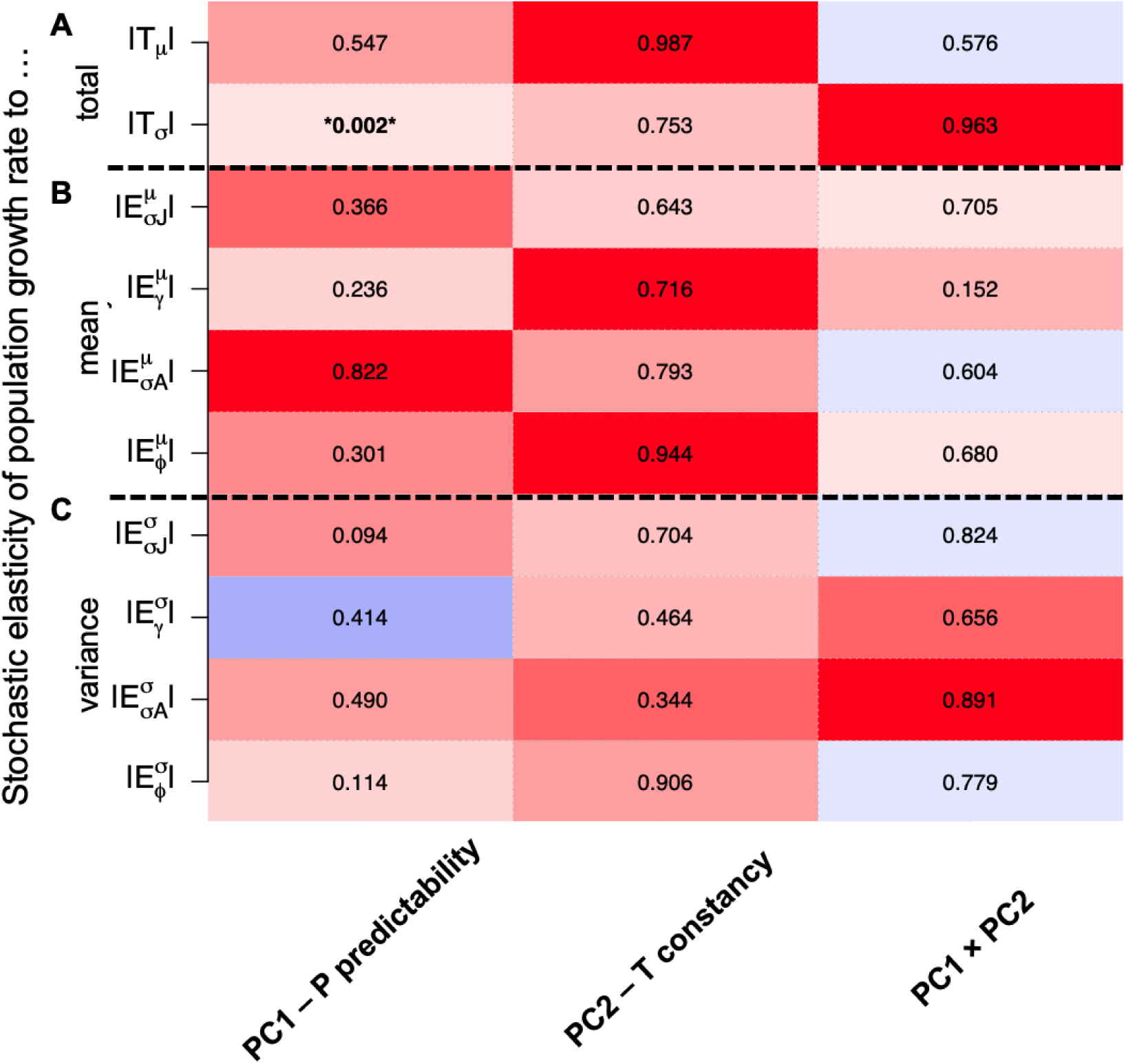
Climatic predictability of demographic buffering The predictability of annual precipitation patterns predicts the ability of animal populations to buffer against extreme climatic events. Scaled effect sizes (negative: red; positive: blue) and P values for the correlations between the climatic PCAs (Figure 2 - PC1: precipitation predictability, PC2: temperature constancy, and their interaction) and: (**A**) the sum of total stochastic elasticities to the mean (|*T*_μ_|) and the variance (|*T*_σ_|), (**B**) the stochastic elasticities to changes in vital rate means (|*E*^μ^|), and (**C**) to changes in vital rate variances (|*E*^σ^|). Vital rates are: juvenile survival (σ*_J_*), maturation (γ), adult survival (σ*_A_*), and reproduction (φ) (Eq. 1).

