## Supplementary material for "Support for the social buffering hypothesis, especially under unpredictable precipitation regimes": SOM

### **Table S1. Sensitivity of results to temporal replication in studies**

**Our overall results are mostly insensitive to the duration of the study.** Here, study duration is defined as the number of matrix population models (MPMs) available in each examined population. Battery of pgls models examining the relationships between the different stochastic elasticities of stochastic population growth rate (*λ_s_*) and sociality, with study duration as a covariate. Note that the latter is only borderline significant in three occasions: *T_σ_,* |*E_σA_^μ^*| and |*E_σA_^σ^*|.

| **Stochastic elasticity to** | | **Predictor** | **Estimate** | **P** |
| --- | --- | --- | --- | --- |
| **A**. Total | Mean - *T_μ_* | Sociality | 0.079 | 0.768 |
|  |  | Duration | -0.200 | 0.040 |
|  | Variance - *T_σ_* | Sociality | 0.807 | 0.399 |
|  |  | Duration | -0.451 | 0.062 |
| **B**. Changes in mean | Juvenile survival - \|*E_σJ_^μ^*\| | Sociality | 0.117 | 0.818 |
|  |  | Duration | -0.086 | 0.487 |
|  | Maturation - \|*E_γ_^μ^*\| | Sociality | -0.738 | 0.318 |
|  |  | Duration | -0.023 | 0.894 |
|  | Adult survival - \|*E_σA_^μ^*\| | Sociality | 0.052 | 0.911 |
|  |  | Duration | -0.241 | 0.046 |
|  | Reproduction - \|*E_φ_^μ^*\| | Sociality | -0.062 | 0.938 |
|  |  | Duration | -0.144 | 0.458 |
| **C**. Changes in variance | Juvenile survival - \|*E_σJ_^σ^*\| | Sociality | 0.108 | 0.906 |
|  |  | Duration | -0.334 | 0.141 |
|  | Maturation - \|*E_γ_^σ^*\| | Sociality | -0.474 | 0.630 |
|  |  | Duration | -0.206 | 0.389 |
|  | Adult survival - \|*E_σA_^σ^*\| | Sociality | 0.185 | 0.470 |
|  |  | Duration | -0.134 | 0.042 |
|  | Reproduction - \|*E_φ_^σ^*\| | Sociality | 0.592 | 0.593 |
|  |  | Duration | -0.417 | 0.131 |

### **Table S2. Metadata and source of matrix population models**

**Sources of the demographic data obtained from COMADRE for the 66 examined species, together with other pertinent details.** *Population* details the number of populations available per species in this study. *Dimension* details the number of stages the original matrix population models (MPMs) had before being collapsed to a set of 2×2 MPMs (see Methods). *Sociality* details the assigned level of our sociality continuum, with the explanation for each of the five levels provided in the Methods too.

| **Species** | **Common name** | **Phylum** | **Class** | **Order** | **Family** | **Population** | **Authors** | **Journal** | **DOI_ISBN** | **Year** | **Dimension** | **Sociality** |
| --- | --- | --- | --- | --- | --- | --- | --- | --- | --- | --- | --- | --- |
| *Acyrthosiphon pisum* | Pea aphid | Arthropoda | Insecta | Aphidomorpha | Aphididae | 1 | Hamda; Jevtic; Laskowski | Ecotoxicology | 10.1007/s10646-012-0904-5 | 2012 | 6 | Gregarious |
| *Agaricia agaricites* | Tan lettuce-leaf coral | Cnidaria | Anthozoa | Scleractinia | Agariciidae | 1 | Hughes; Tanner | Ecology | 10.1890/0012-9658(2000)081[2250:RFLHAL]2.0.CO;2 | 2000 | 3 | Colonial |
| *Alces alces* | Moose | Chordata | Mammalia | Artiodactyla | Cervidae | 1 | Ballard; Whitman; Reed | Wildlife Monogr | https://www.jstor.org/stable/3830713 | 1991 | 3 | Solitary |
| *Ambloplites rupestris* | Rock bass | Chordata | Actinopterygii | Perciformes | Centrarchidae | 1 | Peoples | Master Thesis | NA | 2010 | 3 | Gregarious |
| *Ampelisca abdita* | Amphipod | Arthropoda | Malacostraca | Amphipoda | Ampeliscidae | 1 | Kuhn; Munns; Serbst; Edwards; Cantwell; Gleason; Pelletier; Berry | Environ Toxicol Chem | 10.1002/etc.5620210425 | 2002 | 8 | Gregarious |
| *Anser caerulescens* | Snow goose | Chordata | Aves | Anseriformes | Anatidae | 1 | Cooch; Rockwell; Brault | Ecol Monogr | 10.1890/0012-9615(2001)071[0377:RAODRT]2.0.CO;2 | 2001 | 5 | Colonial |
| *Anthropoides paradiseus* | Blue crane | Chordata | Aves | Gruiformes | Gruidae | 1 | Altwegg; Anderson | Funct Ecol | 10.1111/j.1365-2435.2009.01563.x | 2009 | 5 | Gregarious |
| *Astroblepus ubidiai* | Andean catfish | Chordata | Actinopterygii | Siluriformes | Astroblepidae | 1 | Vélez-Espino | Ecol Freshw Fish | 10.1111/j.1600-0633.2005.00084.x | 2005 | 7 | Gregarious |
| *Boloria eunomia* | Bog fritillary | Arthropoda | Insecta | Lepidoptera | Nymphalidae | 1 | Radchuk; Turlure; Schtickzelle | J Anim Ecol | 10.1111/j.1365-2656.2012.02029.x | 2012 | 5 | Solitary |
| *Brachyteles hypoxanthus* | Northern muriqui | Chordata | Mammalia | Primates | Atelidae | 2 | Morris; Altmann; Brockman; Cords; Fedigan; Pusey; Stoinski; Bronikowski; Alberts; Strier | Am Nat | 10.1086/657443 | 2011 | 10 | Social |
| *Caenorhabditis elegans* | NA | Nematoda | Secernentea | Rhabditida | Rhabditidae | 1 | Li; Ju; Liao; Liao | Ecotoxicology | 10.1007/s10646-014-1267-x) | 2014 | 6 | Gregarious |
| *Callospermophilus lateralis* | Golden-mantled ground squirrel | Chordata | Mammalia | Rodentia | Sciuridae | 1 | Hostetler; Kneip; Van Vuren; Oli | PLOS ONE | 10.1371/jourNAl.pone.0034379 | 2012 | 6 | Solitary |
| *Capitella capitata* | Polychaete | Annelida | Polychaeta | NA | Capitellidae | 1 | Hansen; Forbes; Forbes | Funct Ecol | 10.1046/j.1365-2435.1999.00299.x | 1999 | 2 | Solitary |
| *Cebus capucinus* | White-faced capuchin monkey | Chordata | Mammalia | Primates | Cebidae | 2 | Morris; Altmann; Brockman; Cords; Fedigan; Pusey; Stoinski; Bronikowski; Alberts; Strier | Am Nat | 10.1086/657443 | 2011 | 8 | Social |
| *Centrocercus minimus* | Gunnison sage-grouse | Chordata | Aves | Charadriiformes | Stercorariidae | 1 | Davis; Hooten; Phillips; Doherty | Ecol Evol | 10.1002/ece3.1290 | 2014 | 4 | Solitary |
| *Cercopithecus mitis* | Blue monkey | Chordata | Mammalia | Primates | Cercopithecidae | 1 | Morris; Altmann; Brockman; Cords; Fedigan; Pusey; Stoinski; Bronikowski; Alberts; Strier | Am Nat | 10.1086/657443 | 2011 | 9 | Social |
| *Chlorocebus aethiops* | Vervet | Chordata | Mammalia | Primates | Cercopithecidae | 1 | Isbell; Young; Jaffe; Carlson; Chancellor | Int J Primatol | 10.1007/s10764-009-9332-7 | 2009 | 2 | Social |
| *Chrosomus oreas* | Mountain redbelly dace | Chordata | Actinopterygii | Cypriniformes | Cyprinidae | 1 | Peoples | Master Thesis | NA | 2010 | 2 | Communal |
| *Ciconia ciconia* | White stork | Chordata | Aves | Ciconiiformes | Ciconiidae | 1 | Schaub; Pradel; Lebreton | Biol Conserv | 10.1016/j.biocon.2003.11.002 | 2004 | 2 | Gregarious |
| *Clinocottus analis* | Woolly sculpin | Chordata | Actinopterygii | Scorpaeniformes | Cottidae | 2 | Davis; Levin | Mar Ecol Prog Ser | 10.3354/meps234229 | 2002 | 3 | Solitary |
| *Clinostomus funduloides* | Rosyside dace | Chordata | Actinopterygii | Cypriniformes | Cyprinidae | 1 | Peoples | Master Thesis | NA | 2010 | 2 | Communal |
| *Colias alexandra* | Queen Alexandra's sulphur | Arthropoda | Insecta | Lepidoptera | Pieridae | 1 | Hayes | Oecologia | 10.1007/BF00349187 | 1981 | 7 | Solitary |
| *Cottus bairdii* | Mottled sculpin | Chordata | Actinopterygii | Scorpaeniformes | Cottidae | 1 | Peoples | Master Thesis | NA | 2010 | 3 | Solitary |
| *Erythrocebus patas* | Patas monkey | Chordata | Mammalia | Primates | Cercopithecidae | 1 | Isbell; Young; Jaffe; Carlson; Chancellor | Int J Primatol | 10.1007/s10764-009-9332-7 | 2009 | 2 | Social |
| *Etheostoma flabellare* | Fantail darter | Chordata | Actinopterygii | Perciformes | Percidae | 1 | Peoples | Master Thesis | NA | 2010 | 3 | Solitary |
| *Eulamprus tympanum* | Water skink | Chordata | Reptilia | Squamata | Scincidae | 1 | Blomberg; Shine | Austral Ecol | 10.1046/j.1442-9993.2001.01120.x | 2001 | 5 | Solitary |
| *Falco naumanni* | Lesser kestrel | Chordata | Aves | Falconiformes | Falconidae | 1 | Hiraldo; Negro; Donazar; Gaona | J Appl Ecol | 10.2307/2404688 | 1996 | 2 | Communal |
| *Falco peregrinus* | Peregrine falcon | Chordata | Aves | Falconiformes | Falconidae | 1 | Altwegg; Jenkins; Abadi | Ibis | 10.1111/ibi.12125 | 2013 | 5 | Solitary |
| *Forpus passerinus* | Green-rumped parrotlets | Chordata | Aves | Psittaciformes | Psittacidae | 1 | Sandercock; Beissinger | J Appl Stat | 10.1080/02664760120108818 | 2002 | 2 | Social |
| *Gopherus agassizii* | Desert tortoise | Chordata | Reptilia | Testudines | Testudinidae | 1 | Perez-Heydrich; Oli; Brown | Oikos | 10.1111/j.1600-0706.2011.19735.x | 2011 | 4 | Communal |
| *Gorilla beringei beringei* | Mountain gorilla | Chordata | Mammalia | Primates | Hominidae | 1 | Morris; Altmann; Brockman; Cords; Fedigan; Pusey; Stoinski; Bronikowski; Alberts; Strier | Am Nat | 10.1086/657443 | 2011 | 11 | Social |
| *Helioseris cucullata* | Sunray lettuce coral | Cnidaria | Anthozoa | Scleractinia | Agariciidae | 1 | Hughes; Tanner | Ecology | 10.1890/0012-9658(2000)081[2250:RFLHAL]2.0.CO;2 | 2000 | 3 | Colonial |
| *Homo sapiens sapiens* | Human | Chordata | Mammalia | Primates | Hominidae | 1 | Nicol-Harper; Dooley; Packman; Mueller; Bijak; Hodgson; Townley; Ezard | Popul Ecol | 10.1007/s10144-018-0620-y | 2018 | 18 | Social |
| *Lagopus leucura* | White-tailed ptarmigan | Chordata | Aves | Galliformes | Phasianidae | 1 | Wilson; Martin | BMC Ecol | 10.1186/1472-6785-12-9 | 2012 | 2 | Solitary |
| *Lagopus muta* | Japanese rock ptarmigan | Chordata | Aves | Galliformes | Phasianidae | 1 | Wilson; Martin | BMC Ecol | 10.1186/1472-6785-12-9 | 2012 | 2 | Solitary |
| *Lepus americanus* | Snowshoe hare | Chordata | Mammalia | Lagomorpha | Leporidae | 1 | Meslow; Keith | J Wildlife Manage | 10.2307/3799557 | 1968 | 4 | Solitary |
| *Macaca mulatta* | Rhesus macaque | Chordata | Mammalia | Primates | Cercopithecidae | 1 | Kessler; Pacheco; Rawlings; Ruiz-Lambrides; Delgado; Sabat | Am J Primatol | 10.1002/ajp.22323 | 2014 | 5 | Social |
| *Membranipora membranacea* | Sea mat | Bryozoa | Gymnolaemata | Cheilostomida | Membraniporidae | 2 | Harvell; Caswell; Simpson | Oecologia | 10.1007/BF00323539 | 1990 | 5 | Colonial |
| *Microtus oeconomus* | Root vole | Chordata | Mammalia | Rodentia | Muridae | 1 | Johannesen; Aars; Andreassen; Ims | Popul Ecol | 10.1007/s10144-003-0139-7 | 2003 | 3 | Solitary |
| *Nocomis leptocephalus* | Bluehead chub | Chordata | Actinopterygii | Cypriniformes | Cyprinidae | 1 | Peoples | Master Thesis | NA | 2010 | 3 | Communal |
| *Notamacropus eugenii* | Tammar wallaby | Chordata | Mammalia | Diprotodontia | Macropodidae | 3 | Chambers; Bencini | Wildlife Res | 10.1071/WR10080 | 2010 | 2 | Gregarious |
| *Orbicella annularis* | Caribbean star coral | Cnidaria | Anthozoa | Scleractinia | Faviidae | 2 | Hughes; Tanner | Ecology | 10.1890/0012-9658(2000)081[2250:RFLHAL]2.0.CO;2 | 2000 | 3 | Colonial |
| *Orcinus orca* | Killer whale | Chordata | Mammalia | Cetacea | Delphinidae | 2 | Vélez-Espino; Ford; Araújo; Ellis; Parken; Balcomb | Can Tech Report Fish & Aq Sci | 978-1-100-23563-9 | 2014 | 7 | Social |
| *Ovis aries* | Soay sheep | Chordata | Mammalia | Artiodactyla | Bovidae | 1 | Clutton-Brock; Price; Albon; Jewell | J Anim Ecol | 10.2307/5330 | 1992 | 6 | Gregarious |
| *Pagurus longicarpus* | Long-clawed hermit crab | Arthropoda | Malacostraca | Decapoda | Paguridae | 1 | Damiani | Ecology | 10.1890/04-0956 | 2005 | 3 | Solitary |
| *Pan troglodytes schweinfurthii* | Eastern chimpanzee | Chordata | Mammalia | Primates | Hominidae | 1 | Morris; Altmann; Brockman; Cords; Fedigan; Pusey; Stoinski; Bronikowski; Alberts; Strier | Am Nat | 10.1086/657443 | 2011 | 17 | Social |
| *Papio cynocephalus* | Olive baboon | Chordata | Mammalia | Primates | Cercopithecidae | 1 | Morris; Altmann; Brockman; Cords; Fedigan; Pusey; Stoinski; Bronikowski; Alberts; Strier | Am Nat | 10.1086/657443 | 2011 | 8 | Social |
| *Paramuricea clavata* | Violescent sea-whip; Red gorgonian | Cnidaria | Anthozoa | Alcyonacea | Plexauridae | 2 | Linares; Doak | Mar Ecol Prog Ser | 10.3354/meps08437 | 2010 | 7 | Colonial |
| *Pimephales promelas* | Fathead minnow | Chordata | Actinopterygii | Cypriniformes | Cyprinidae | 1 | Schwindt | NA | NA | 2013 | 4 | Communal |
| *Plexaura homomalla* | Gorgonian coral | Cnidaria | Anthozoa | Alcyonacea | Plexauridae | 1 | Lasker | Oecologia | 10.1007/BF00318316 | 1991 | 3 | Colonial |
| *Porites astreoides* | Caribbean coral reef | Cnidaria | Anthozoa | Scleractinia | Poritidae | 1 | Edmunds | Mar Ecol Prog Ser | 10.3354/meps08595 | 2010 | 3 | Colonial |
| *Propithecus verreauxi* | Verreaux's sifaka | Chordata | Mammalia | Primates | Indriidae | 1 | Morris; Altmann; Brockman; Cords; Fedigan; Pusey; Stoinski; Bronikowski; Alberts; Strier | Am Nat | 10.1086/657443 | 2011 | 8 | Social |
| *Pseudodiploria strigosa* | Symmetrical brain coral | Cnidaria | Anthozoa | Scleractinia | Faviidae | 1 | Edmunds | Mar Ecol Prog Ser | 10.3354/meps08595 | 2010 | 3 | Colonial |
| *Pygoscelis adeliae* | Adelie penguin | Chordata | Aves | Sphenisciformes | Spheniscidae | 1 | Hinke; Trivelpiece; Trivelpiece | Ecosphere | 10.1002/ecs2.1666 | 2017 | 2 | Colonial |
| *Rana temporaria* | European common frog | Chordata | Amphibia | Anura | Ranidae | 1 | Campbell; Garner; Tessa; Scheele; Griffiths; Wilfert; Harrison | PeerJ | NA | 2018 | 11 | Gregarious |
| *Sceloporus grammicus* | Mesquite lizard | Chordata | Reptilia | Squamata | Phrynosomatidae | 2 | Méndez–de la Cruz; Zúñiga-Vega; Cuellar | Can J Zool | 10.1139/Z08-124 | 2008 | 3 | Solitary |
| *Scolytus ventralis* | Fir engraver beetle | Arthropoda | Insecta | Coleoptera | Curculionidae | 1 | Berryman | Can Entomol | 10.4039/Ent1051465-11 | 1973 | 6 | Solitary |
| *Strix occidentalis* | Northern spotted owl | Chordata | Aves | Strigiformes | Strigidae | 1 | LaHaye; Zimmerman; Gutiérrez | Auk | 10.1642/0004-8038(2004)121[1056:TVITVR]2.0.CO;2 | 2004 | 3 | Solitary |
| *Suricata suricatta* | Meerkat | Meerkat | Mammalia | Carnivora | Herpestidae | 1 | Conquet,Ozgul, Blumstein, Armitage, Oli, Martin, Clutton-Brock, Paniw | Ecosphere | 10.1002/ecy.3894 | 2023 | 4 | Social |
| *Turdus torquatus* | Ring ouzel | Chordata | Aves | Passeriformes | Turdidae | 1 | Sim; Rebecca; Ludwig; Grant; Reid | J Anim Ecol | 10.1111/j.1365-2656.2010.01750.x | 2011 | 2 | Solitary |
| *Umbonium costatum* | NA | Mollusca | Gastropoda | Vetigastropoda | Trochidae | 1 | Noda; Nakao | J Anim Ecol | 10.2307/5722 | 1996 | 6 | Communal |
| *Ursus maritimus* | Polar bear | Chordata | Mammalia | Carnivora | Ursidae | 1 | Hunter; Caswell; Runge; Regehr; Amstrup; Stirling | Ecology | 10.1890/09-1641 | 2010 | 6 | Solitary |
| *Vireo atricapilla* | Black-capped vireo | Chordata | Aves | Passeriformes | Vireonidae | 11 | Walker; Marzluff; Cimprich | Biol Conserv | 10.1016/j.biocon.2016.09.016. | 2016 | 2 | Solitary |
| *Xenosaurus agrenon* | Knob-scaled lizard | Chordata | Reptilia | Squamata | Xenosauridae | 1 | Zamora-Abrego; Chang; Zúñiga-Vega; Nieto-Montes de Oca; Johnson | Herpetologica | 10.1655/09-005.1 | 2010 | 4 | Solitary |
| *Xenosaurus grandis* | Crevice-dwelling lizard | Chordata | Reptilia | Squamata | Xenosauridae | 1 | Zúñiga-Vega; Valverde; Rojas-Gonzalez; Lemos-Espinal | Copeia | 10.1643/0045-8511(2007)7[324:AOTPDO]2.0.CO;2 | 2007 | 4 | Solitary |
| *Xenosaurus platyceps* | Flathead knob-scaled lizard | Chordata | Reptilia | Squamata | Xenosauridae | 2 | Rojas-Gonzalez; Jones; Zúñiga-Vega; Lemos-Espinal | Amphibia-Reptilia | 10.1163/156853808784124992 | 2008 | 4 | Solitary |

### **Table S3. Degree of sociality across the climatic space**

### **The degree of sociality is not predicted by the climatic principal component axis.** Results of a pgls model predicting the degree of sociality as a function of the positioning of each population along PC1 (Precipitation predictability; Figure 2) and PC2 (temperature constancy).

|  | **Estimate** | **S. E.** | **t** | **P** |
| --- | --- | --- | --- | --- |
| **Intercept** | 2.616 | 1.1876 | 2.2032 | 0.0436 |
| **PC1** | 0.1150 | 0.2581 | 0.4453 | 0.6624 |
| **PC2** | 0.2682 | 0.3579 | 0.7493 | 0.4652 |
| **PC1 × PC2** | 0.0165 | 0.1845 | 0.0894 | 0.9299 |

### **Figure S1. Sampling frequency of the matrix population models**

**Most of the matrix population models (MPMs) in our selected set of matrix population models from the COMADRE Animal Matrix Database were sampled once a year.** Species in our study with an original sampling frequency that is not annual: *Caenorhabditis elegans* (0.000114), *Capitella capitata* (0.0192), *Xenosaurus agrenon* (0.08), *Membranopira mebranaceae* (0.12), *Acyrthosiphon pisum* (0.02), *Clinocottus analis* (0.25), *Ampelisca abdita* (0.03), *Eulamprus tympanum* (0.08), *Sceloporus grammicus* (0.5), *Montastrae annularis* (5), and Homo sapiens sapiens (5).

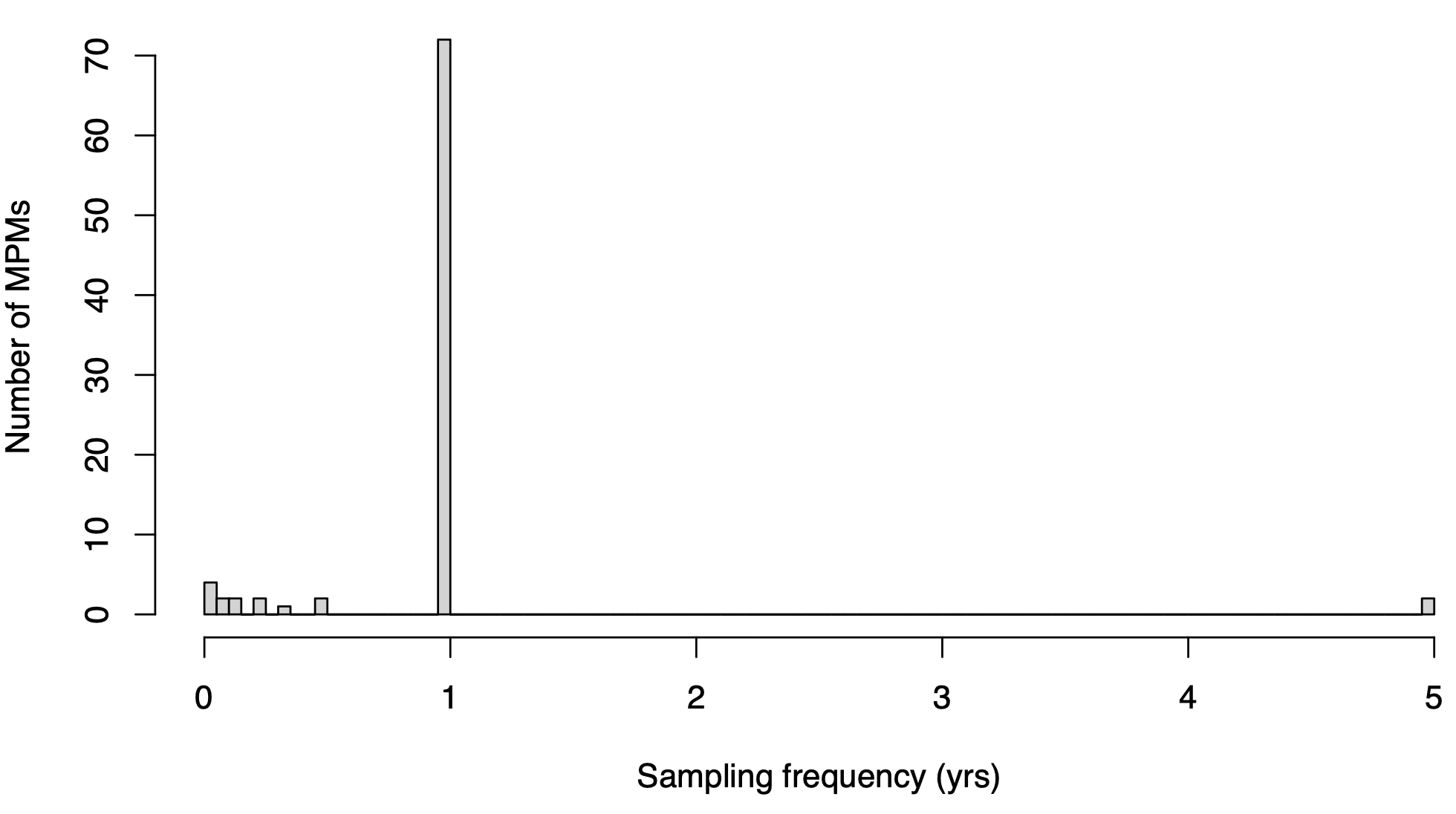

### **Figure S2. Matrix population model availability per study**

**Number of matrix population models (MPMs) per studied population in our selected subset of 66 species from the COMADRE Animal Matrix.** Total MPM = 955. Total number of separate populations = 87.

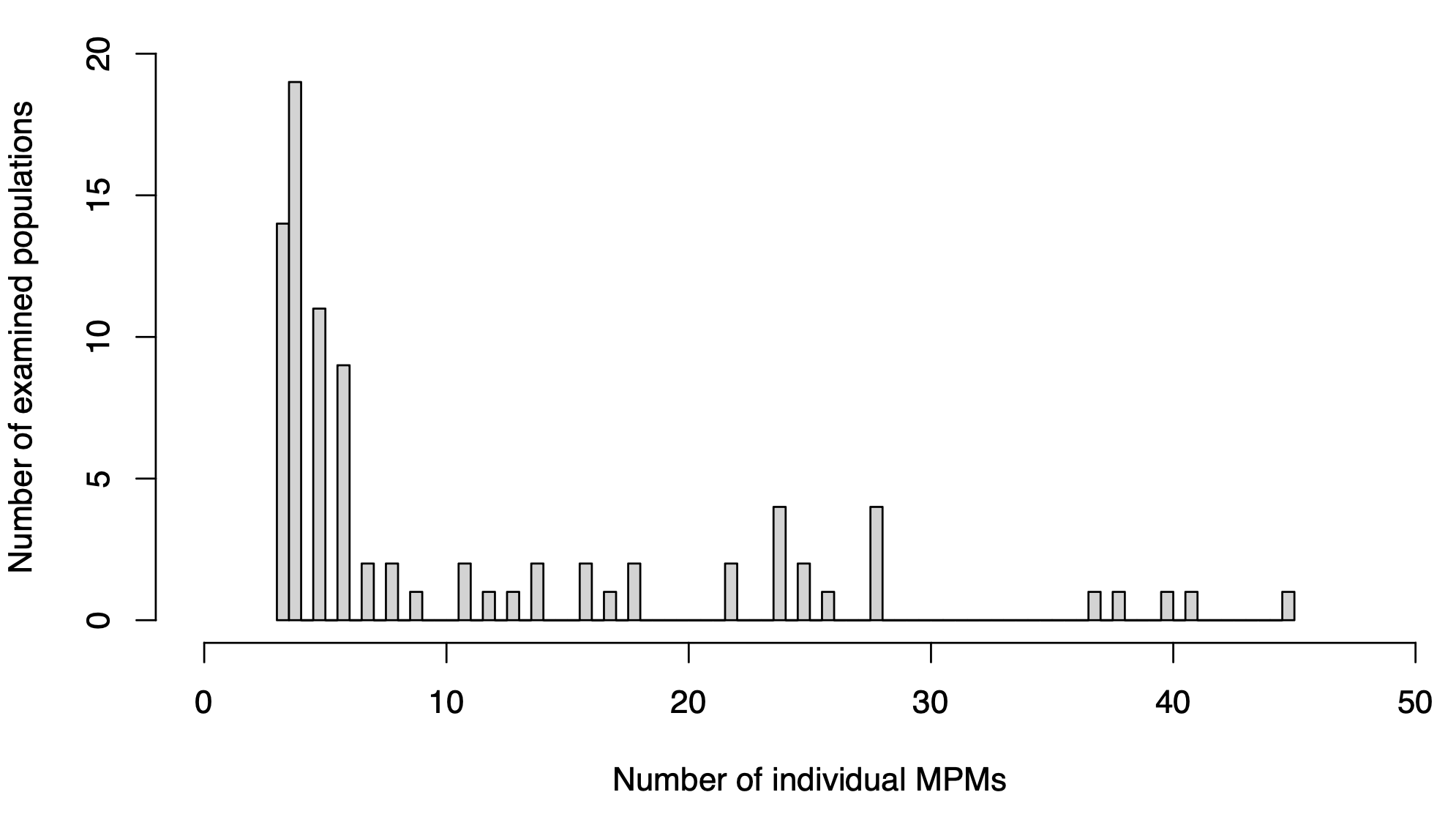

### **Figure S3. Matrix population model dimensionality**

**The dimensionality of our examined matrix population models (MPMs) varied considerably between 2 and 18 stages.** As such, we collapsed all models to a two-stage MPM to allow for stage comparability in vital rates.

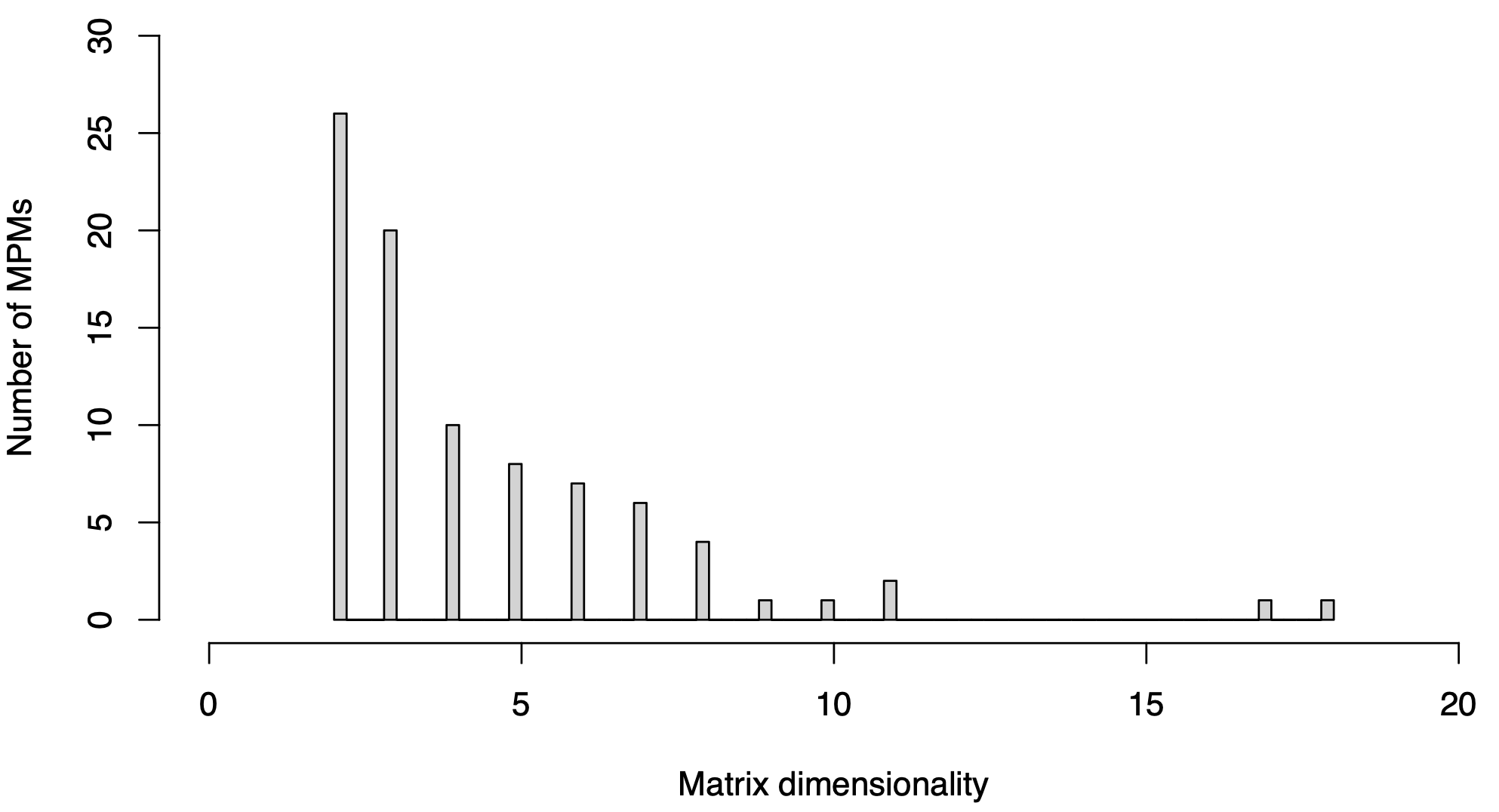

### **Figure S4. Climate driver collinearities**

**Some of the examined climatic drivers are strongly collinear.** Pairwise correlations (top-right) and scatterplots (bottom-left) of the statistics of monthly precipitation and temperature derived from the NASA POWER database to quantify the degree of environmental stochasticity to which our 87 natural populations have been exposed. Temperature statistics include: *T̄*: mean; *ΔT*: variance: *T_cons_*: constancy; *T_cont_*: contingency; and *T_pred_*: predictability. Ditto for precipitation (*P*). Constancy quantifies the extent to which a climatic variable remains stable over time (1 - variance/mean); contingency quantifies the extent to which fluctuations are structured and recurrent; predictability = constancy + contingency, as per Colwell (Colwell, 1974). Font size of spearman correlation coefficient is proportional to its values.

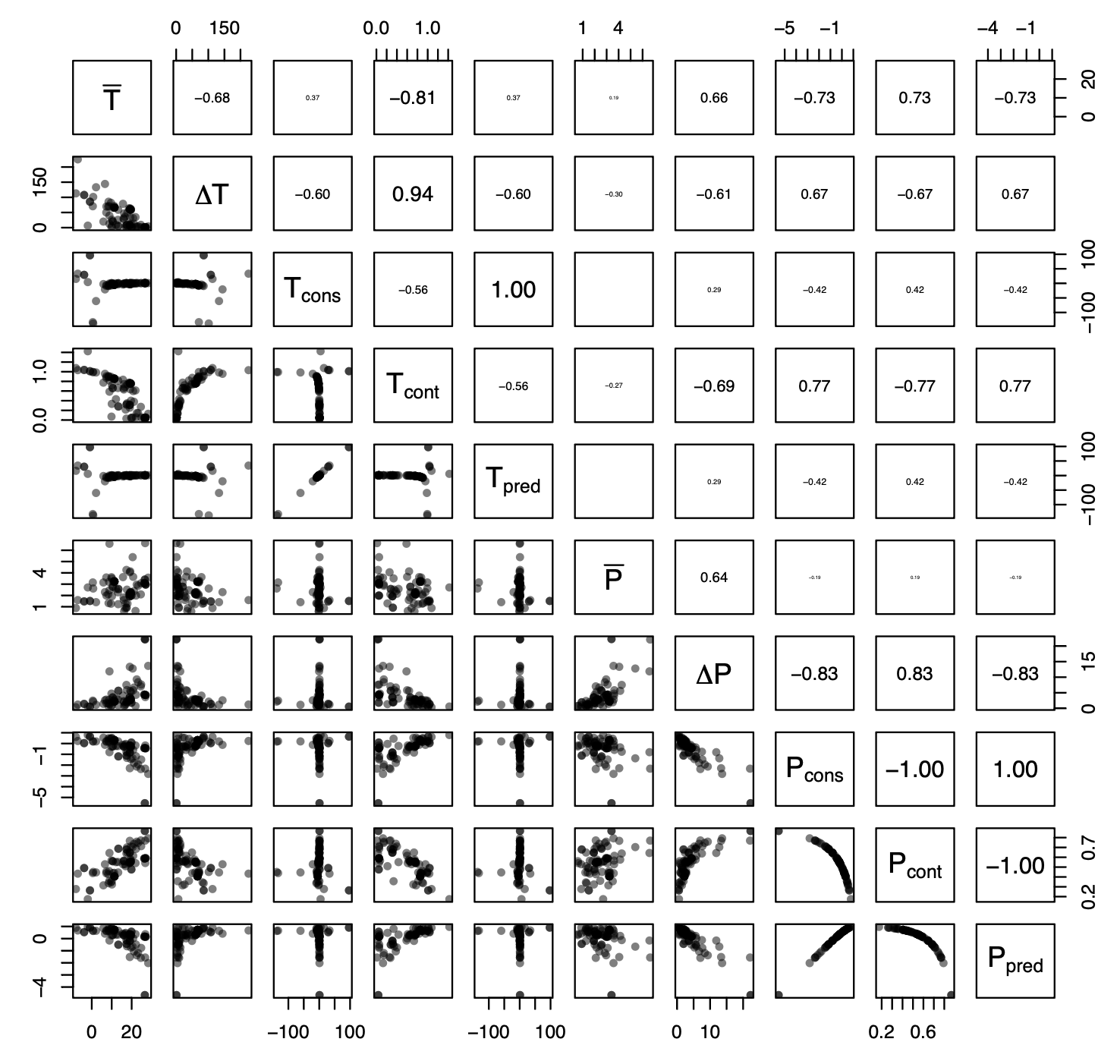

### **Figure S5. PCA screeplot**

**Screeplot of the six principal components of the climatic PCA.** The blue dashed line separates the principal components axis with associated eigenvalue > 1, which we retained for the next steps in our analyses, as per the Kaiser criterion.

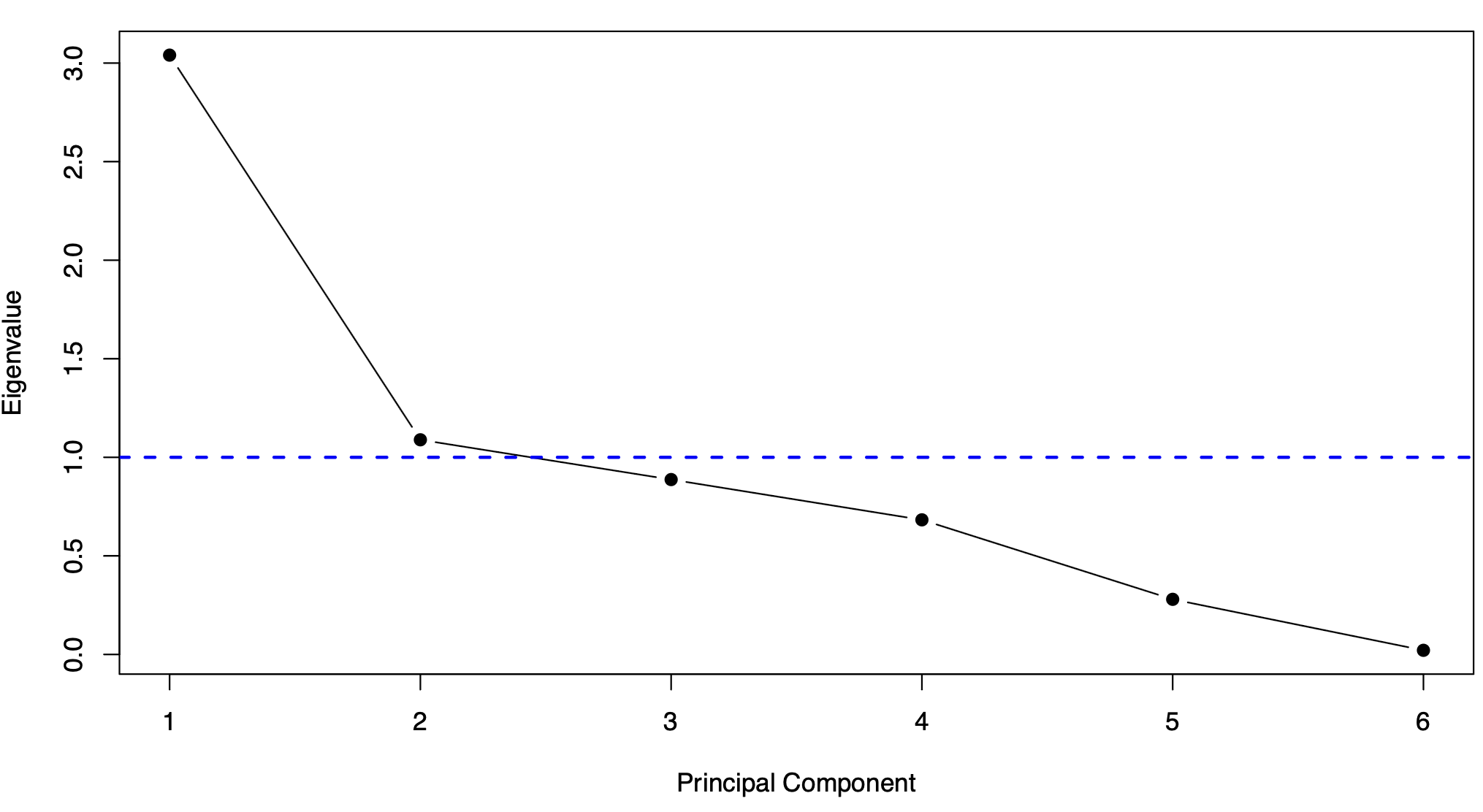

### **Figure S6. Climatic predictability of demographic buffering**

**The predictability of annual precipitation patterns predicts the ability of animal populations to buffer against extreme climatic events**. Scaled effect sizes (negative: red; positive: blue) and P values for the correlations between the climatic PCAs (Figure 2 - PC1: precipitation predictability, PC2: temperature constancy, and their interaction) and: (**A**) the sum of total stochastic elasticities to the mean (|*T_μ_*|) and the variance (|*T_σ_*|), (**B**) the stochastic elasticities to changes in vital rate means (|*E^μ^*|), and (**C**) to changes in vital rate variances (|*E^σ^*|). Vital rates are: juvenile survival (*σ_J_*), maturation (*γ*), adult survival (*σ_A_*), and reproduction (*φ*) (Eq. 1).

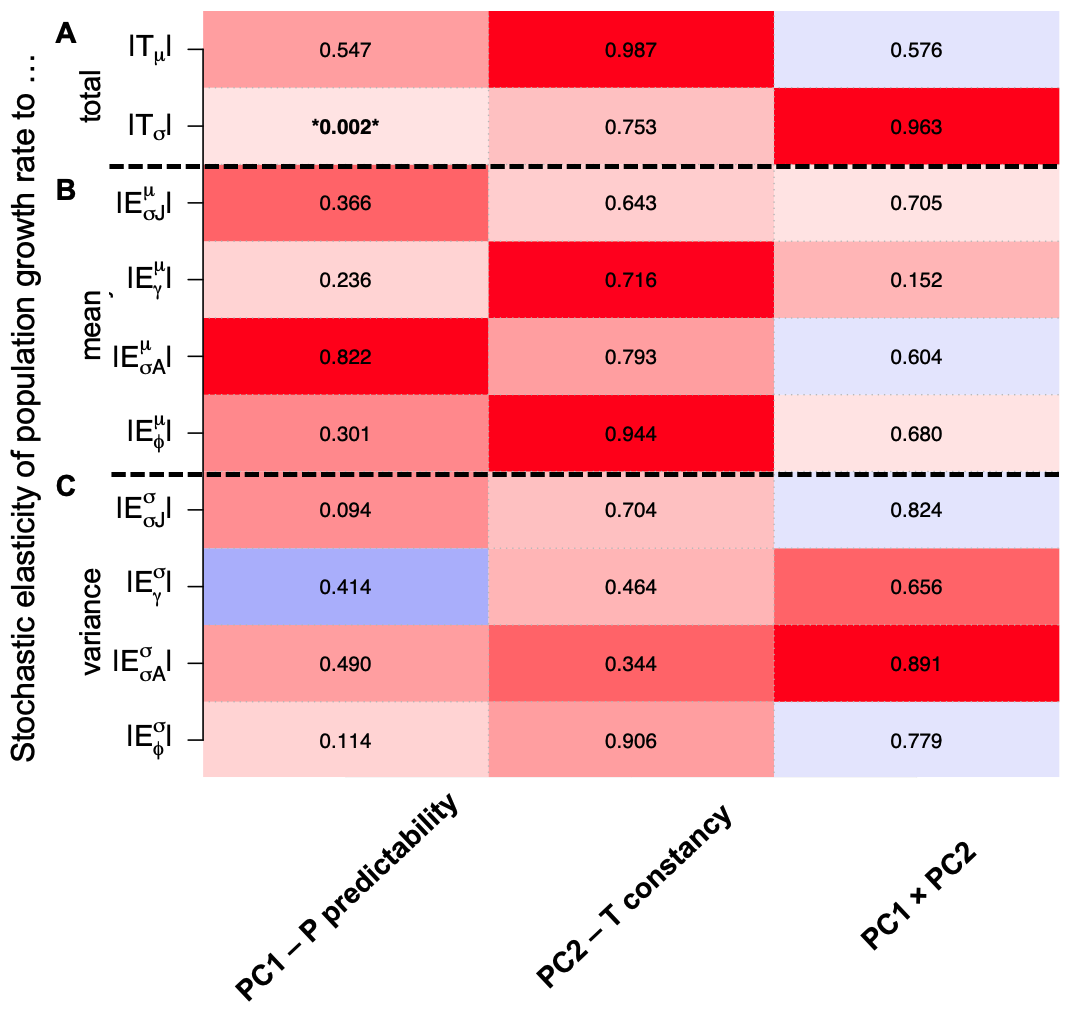
